# DeepCDS: *Ab initio* coding sequence prediction in prokaryotic metagenomic short reads

**DOI:** 10.64898/2026.06.17.732633

**Authors:** Line Sandvad Nielsen, Henrik Nielsen, Ole Winther

**Author notes:** Contributing authors. These authors contributed equally to this work (Henrik Nielsen, Ole Winther).

## Abstract

Accurate coding sequence prediction in short prokaryotic metagenomic sequences, such as Illumina reads, remains challenging due to sequence fragmentation, unknown sequence origins, and sequencing errors. Here we introduce DeepCDS, a deep learning-based *ab initio* coding sequence predictor trained on short prokaryotic sequences with and without simulated Illumina-like sequencing errors. DeepCDS integrates embeddings from the protein language model ESM-2 with nucleotide-level information to predict complete and partial coding sequences. Benchmarking on 215 phylogenetically diverse prokaryotic organisms demonstrates that DeepCDS consistently outperforms well-established and widely used methods in coding sequence detection across sequence lengths from 75 to 1,000 base pairs, including improved identification of both canonical and non-canonical start codons as well as stop codons, and greater robustness to different sequencing error profiles. DeepCDS also recovers more coding sequences from functionally uncharacterized genes and remains operational on sequences shorter than existing tools support. We additionally evaluate DeepCDS on a simulated metagenomic dataset from the Critical Assessment of Metagenome Interpretation initiative, where it likewise achieves the strongest performance among tested tools, and on a real human gut metagenome, where its predictions are more frequently supported by known proteins. These findings suggest that amino acid-level context, here provided by a protein language model, contributes complementary information for nucleotide-level coding sequence detection, especially at very short lengths. Ultimately, DeepCDS may help uncover the functional potential of the vast microbial diversity that remains genomically uncharacterized.

## 1 Introduction

Identification of protein-coding genes and their corresponding coding sequences (CDSs) is a well-established task in prokaryotic genomics. Prokaryotic genomes are typically small, ranging from 0.1 to 16 million base pairs (bps) [1], lack introns, and exhibit high gene density, with protein-coding regions comprising 80-90% of most genomes (Supplementary Figure A1A-B) [2–4]. This compact architecture is thought to reflect selection for fast replication [4]. Consequently, gene finding in prokaryotes often concerns identifying the correct reading frame (RF).

Over the past decades, numerous computational methods have achieved high accuracy in prokaryotic gene finding for whole-genome sequence data [5–8], typically relying on assembled genomes as input [2, 3]. However, assembly is not always feasible, and even when possible, it can still result in incomplete, misassembled, or contaminated genomes, which complicates downstream annotation [3, 9]. In metagenomic samples, these challenges are amplified by the high microbial diversity of mixed communities, where species can occur in notably different abundances (e.g., in soil, the human gut, sea water, and fermented foods), leading to low effective sequencing depth per species. This uneven sequencing depth can make assembly from short reads challenging, and even when successful, many reads may not be part of the final assemblies, as reads from low-abundance species may not reach the depth required to form contigs [6, 10–13]. Homology-based alignment methods such as BLASTX [14] are widely used to map such short sequences to reference databases, but they often yield limited results when dealing with error-prone sequences or sequences originating from previously uncharacterized organisms [6, 11]. For example, despite continuous expansion of prokaryotic reference genome catalogs, more than 20% of sequencing reads from human faecal metagenomes remain unmapped [15], representing a potentially large set of novel genes. Another challenge in metagenomic contexts is that the genomes from which sequences originate are unknown, and some may follow non-standard genetic codes [13]. For example, bacteria belonging to the orders *Mycoplasmatales* and *Entomoplasmatales* use NCBI genetic code 4 (translation table 4), where the codon TGA encodes tryptophan (W) rather than functioning as a stop codon [16, 17]. This motivates the development of *ab initio*-based models that can predict CDS directly from short, unassembled sequencing reads without prior knowledge of the sequence origin [12, 18].

This task is, however, challenging for several reasons: First, most prokaryotic CDSs are substantially longer than typical short reads (150 to 300 bps) generated by standard Illumina sequencing platforms (see Supplementary Figure A1D). Fragmented sequences covering only part of a coding region are inherently difficult to predict as they lack the context of full genes, and performance generally declines as input sequence length decreases [2, 6, 12, 13, 19]. Second, sequencing errors, in the form of artifactual substitutions, insertions, and deletions (indels), further complicate prediction by disrupting RFs, introducing premature stop codons, or altering stop codons to non-stop codons. While modern Illumina platforms (e.g., HiSeq, MiSeq, NextSeq) exhibit low error rates (indel errors in the order of 10*^−^*^6^ per base, substitution errors in the order 10*^−^*^3^ per base [20–22]), CDS prediction remains sensitive to such errors [6, 19]. Throughout this work, substitutions and indels refer to errors introduced during sequencing and not genuine biological variation between genomes.

From a historical perspective, the aim of predicting CDSs from error-prone short sequences is not new. As early as 1999, ESTScan [23] was proposed, based on a hidden Markov model that explicitly accounted for sequencing errors and was trained to predict coding regions from low-quality human expressed sequence tags (ESTs). Since then, several gene prediction tools have been developed to handle short and error-prone sequences [6, 11, 13, 24, 25]. FragGeneScan (FGS) [6], introduced in 2010, employs a hidden Markov model that integrates explicit sequencing error models based on data from 139 bacterial genomes. As it remains widely used for CDS prediction from short and error-prone reads, the tool was reimplemented as FragGeneScanRs [12] in 2022 to improve computational efficiency. MetaProdigal [13] from 2012 is an adaptation of the whole-genome gene finder Prodigal [5] but optimized for short metagenomic contigs of unknown origin. It uses a dynamic programming approach with 50 training files derived from clustering 1,415 prokaryotic RefSeq genomes. The successor of MetaGeneMark from 2010 [25], MetaGeneMark-2 [26], was released in 2022. It employs a generalized hidden Markov model based on data from 5,491 prokaryotic RefSeq genomes, and was evaluated on metagenomic contigs of 250 bp and longer. Neither MetaProdigal nor the MetaGeneMark models explicitly account for sequencing errors. Despite this, these models are widely used for CDS prediction in metagenomic sequence data.

Compared to the usage of traditional sequence-based features, protein language models (pLMs) such as ESM-2 [27], ESM C [28], and ProtT5 [29] have substantially enhanced protein sequence representations by learning evolutionary, structural, and functional dependencies across large corpora of unlabeled protein sequences [27, 29, 30]. By capturing these contextual relationships across amino acids in a sequence, these models have led to improved performance across a range of downstream prediction tasks. Recent work on eukaryotic start codon prediction [31] suggests that the utility of pLMs extends beyond encoding complete proteins, as their learned understanding of universal protein features can also be leveraged to distinguish which features do not resemble those of a protein, motivating their use in nucleotide-level prediction tasks. An alternative approach is to model the nucleotide sequence directly using language models trained on DNA sequences, which have been applied to a range of genomic prediction tasks [32]. However, applying a pLM to the translated RFs makes it possible to leverage the protein-level knowledge captured during pretraining, which is relevant for distinguishing coding from non-coding sequences. Building on this idea, we introduce DeepCDS: an *ab initio* deep learning model that integrates nucleotide-level codon encodings with fine-tuned pLM representations of translated RFs to predict complete and partial CDSs from short prokaryotic DNA sequences.

DeepCDS was trained on simulated sequences of 300 bps from 813 prokaryotic genomes, and takes only a DNA sequence as input without prior knowledge of its genomic origin. The model is available in three variants trained under increasing levels of sequencing noise: error-free, substitution errors only, and both substitution and indel errors, making it adaptable to a wide range of sequence contexts and data qualities. We benchmarked DeepCDS against FGS and MetaProdigal across diverse phylogenetic groups on test sets of varying sequence lengths and error rates, evaluating CDS detection performance and the ability to accurately identify start codon and stop codon positions. Integrating protein-level context with nucleotide-level information, DeepCDS substantially improves CDS detection performance, and is able to identify CDSs in fragments shorter than what current well-established tools support. DeepCDS can be used as a stand-alone tool or to complement existing whole-genome annotation pipelines by recovering partial CDS regions that assembly-based predictors may miss.

## 2 Materials and Methods

### 2.1 Data Sources

We collected the raw datasets from NCBI’s Genome database [33], specifically retrieving reference genomes including plasmids along with their corresponding RefSeq and GenBank annotations in .gff-format. In total we collected 1,125 genomes, distributed on 117 archaeal organisms and 1,008 bacterial organisms. The genomes were selected based on the following criteria: (i) having assembly status as “complete”, (ii) belonging to a defined family in NCBI’s Taxonomy classification system [34] (used for genome partitioning, see Section 2.3), (iii) covering a broad range in GC-content. All data were obtained using the most recent genome assemblies and annotations available as of September 2025. Metadata for each genome are provided in Supplementary Tables A1 and A2.

### 2.2 Data Preprocessing

#### Short read simulation

We used Mason [35] for simulating short reads from the genomes because of its accurate resemblance to modern Illumina platforms, based on a benchmark study from 2023 [22]. This includes an approximately uniform sampling of reads across the full genome and a close mimicking of Illumina substitution and indel error profiles, the former increasing toward read ends due to reduced signal intensity, and the latter tending to be relatively evenly distributed along the read [20, 22, 36]. Furthermore, Mason outputs a file with the ground truth set of aligned reads including sequencing error positions and genomic reference coordinates (“golden” .bam file), which is an important attribute for the supervised learning setup employed in this project.

We ran Mason separately on the forward and complement strand, because this made subsequent data processing clearer compared to sequencing from both strands together. Because the objective of our model is to predict coding regions solely based on the information embedded within individual short read sequences, we chose to simulate single-end reads. For the complement strand, we reverse-complemented the genome sequence prior to simulating reads. For each genome, we simulated reads corresponding to a sequencing depth of 1x per strand, so that each base on average was “sequenced” once. The number of reads simulated for each strand of a genome was calculated as 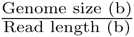. Although Mason was first introduced in 2010, the method is actively maintained (we used mason simulator v2.0.12, which was updated on the 2nd of April 2025).

For the training and validation sets, we simulated reads with a length of 300 bps with Mason, employing the default Illumina sequencing error profile (i.e., distribution of indel- and substitution error rates across the read). We set the average probability for a substitution error occurring to 5 *·* 10*^−^*^3^ per base (0.5% substitution error rate). Early experiments showed that our model was quick to learn to distinguish fully non-coding and coding sequences, whereas it was more challenged by learning the transitions between coding and non-coding regions, especially those caused by indel errors rather than start- and stop codons. As indel errors are rare at realistic Illumina error rates, training data simulated at such rates contained too few examples for the model to learn these transitions. For this reason, we increased the representation of indel errors in our training data, setting the rate of indel errors to be 2.5*·*10*^−^*^4^ per base each (0.05% indel error rate) in both the training and validation splits. We created two additional training and validation datasets with Mason, namely one only with substitution errors (0.5% substitution error rate), and one without any sequencing errors.

#### Short read processing and labeling

The simulated reads were labeled and processed to fit the input format of our model. We first collected CDS annotations from the .gff-files sourced from both RefSeq and GenBank for each reference genome. We removed duplicate CDSs annotated by both sources.

When two CDS annotations shared the same stop codon but differed in start codon positions, we kept the longer CDS to retain as much potential coding sequence as possible. Since this may have created a bias in the trained model for choosing the longer CDS (upstream start codon) we conducted a separate benchmark study of those reads that contained two differently annotated start codons (Results Section 3.3, “Model behavior on CDSs with conflicting annotated start positions”). In rare cases two CDS annotations shared start codon but different stop codon positions, where the longer CDS consistently encoded a selenocysteine at the position corresponding to the stop codon of the shorter sequence. In these cases, we kept the longer CDS.

The CDS annotations obtained from RefSeq and GenBank together with the reference alignment .bam-file produced by Mason, were used to identify coding regions and sequencing error positions within the simulated reads, by mapping each read’s position back to the reference genome using its start coordinate (Figure 1, “Data preprocessing”). We aligned each read to overlapping CDS annotations (when present in the corresponding region) and extracted the coordinates and RF of each CDS fragment relative to the read’s start position. Throughout, we refer to the part of a CDS contained in a sequence as a CDS fragment, which may comprise a complete CDS or, more commonly, a partial CDS truncated by the sequence boundaries. A CDS fragment split by an indel error into two or more segments is referred to as an indel-interrupted CDS. We adjusted all CDS coordinates annotated on the complement strand to align correctly with the reads simulated from the reverse-complemented genomic sequence. If a CDS annotation was given as [CDS_start_, CDS_stop_], and the genomic sequence had length *L*, the corrected CDS coordinates were calculated as [*L* + 1 *−* CDS_stop_*, L* + 1 *−* CDS_start_] (where the +1 accounts for the 1-based indexing of GFF coordinates).

**Fig. 1:**
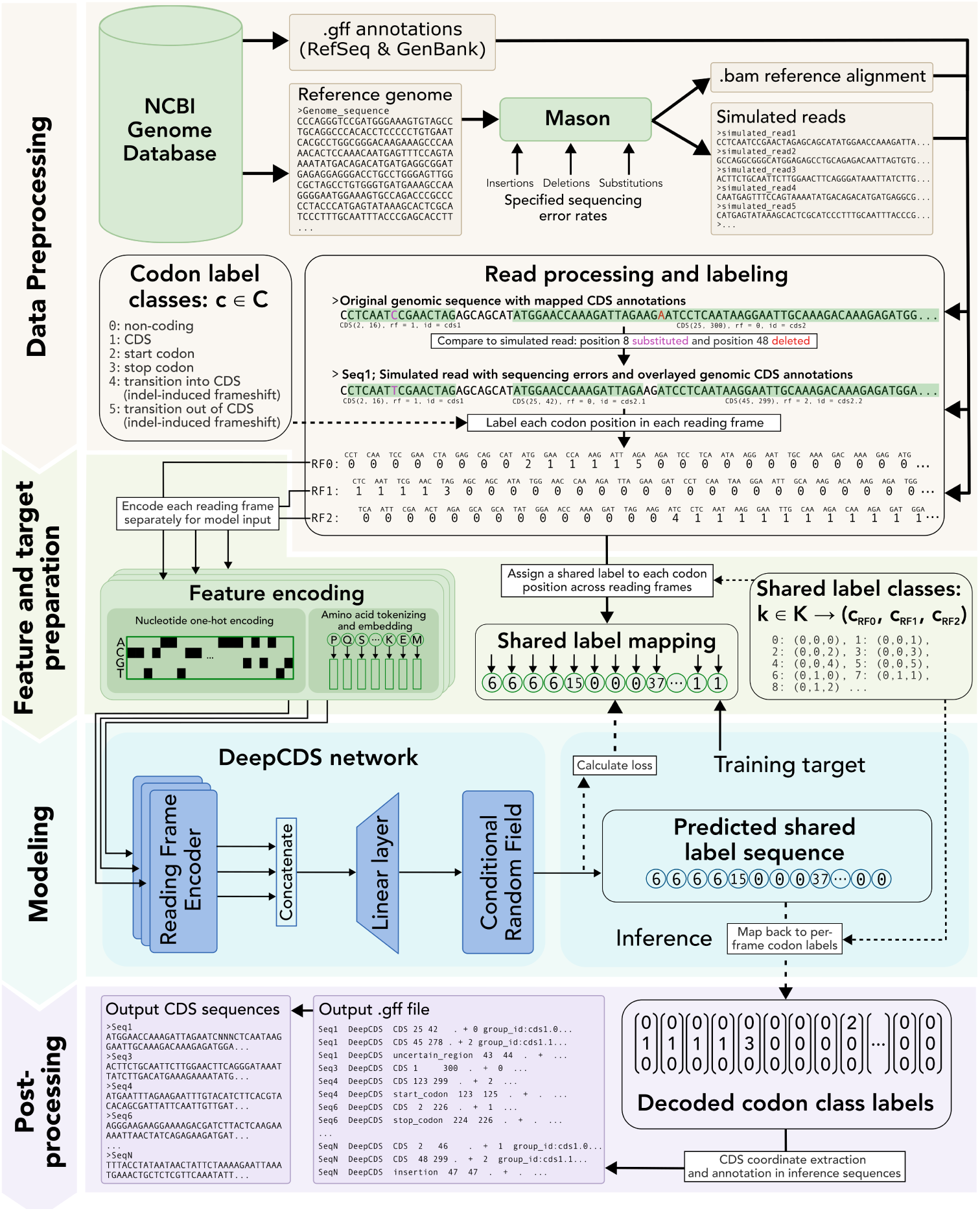
Overview of the DeepCDS development pipeline. Overall workflow for the development of DeepCDS, from data collection to model training and user inference output. **Data preprocessing.** Reference genomes and their corresponding RefSeq and GenBank annotations were collected from the NCBI Genome database [33]. Mason [35] was used to simulate short reads from the genome sequence with Illumina-like, substitution-only, and error-free profiles. The simulated reads were labeled as follows: Each codon position in each RF (RF_0_,RF_1_,RF_2_) was assigned a label from the codon label classes *c ∈ C*, where *C* = *{*0, 1, 2, 3, 4, 5*}*, based on the genomic .gff annotations and the .bam reference alignment produced by Mason. For the datasets simulated without indel errors, the same labeling convention was used but excluding codon class labels 4 and 5. **Feature and target preparation.** The codon class labels across RFs were jointly encoded per position, such that each unique combination of (*c*_RF0_, *c*_RF1_, *c*_RF2_) maps to a single index *k*. For each RF, the codon sequence was encoded individually in two ways: first, the nucleotide sequence was one-hot encoded per codon, and second, the codon sequence was translated to its corresponding amino acid sequence and embedded with ESM-2 [27]. **Modeling.** The DeepCDS model was trained end-to-end, minimizing the negative log-likelihood of the true shared label sequences (Figure 2). **Post-processing.** During inference, the predicted shared label sequence is mapped back to the per-frame codon labels, from which CDS coordinates, start and stop codon positions, and indel errors are extracted and written to .gff and FASTA output files.

For reads containing indel errors, the RF of a coding region is disrupted (Figure 1, “Data preprocessing”). We adjusted the overlapping CDS annotations accordingly during the labeling process. Specifically, an insertion error shifts the annotation one base forward (RF_0_ *→* RF_1_ *→* RF_2_ *→* RF_0_), whereas a deletion error shifts it one base backward (RF_0_ *→* RF_2_ *→* RF_1_ *→* RF_0_).

Ultimately, the performance of a machine learning model is limited by the quality of the data it is trained on. For this reason, we conducted several quality checks and discarded reads containing positions that could not confidently be labeled as coding or non-coding, by checking the following: (1) We kept track of all CDS annotations that were marked as either incomplete, hypothetical (such as being predicted or not yet functionally characterized), or with a frameshift where the position of the frameshift was not noted. Specifically, we searched for the tags “pseudo=true”, “product=hypothetical protein”, “partial=true”, “ab initio prediction” or “note=programmed frameshift”, and checked that a CDS corresponded to a complete number of codons. We then removed all simulated reads overlapping with any of these uncertain CDS ranges from our dataset. (2) We removed reads containing unknown bases (marked by any other letter than A, T, G, or C). (3) We checked that the amino acid sequence of each labeled CDS corresponded to a protein fragment present in the proteome of the corresponding reference genome (simulated substitution errors were “substituted back” to the true bases while conducting this check).

Each codon in each of the three RFs was assigned a label *c ∈ C*, where *C* = *{*0, 1, 2, 3, 4, 5*}* (Figure 1, “Data preprocessing”) using the processed annotations. For the datasets simulated with only substitution errors, and without any sequencing errors, respectively, we used the same labeling convention but excluding labels 4 and 5. Labeling at codon-level (and thereby for each RF) rather than at the nucleotide level allowed us to easily identify the correct RF and better handle overlapping coding regions in different RFs [37].

### 2.3 Dataset Partitioning

Many biological sequences share substantial similarity due to their evolutionary relationships. While deep learning models have proven successful in learning complex patterns from such data, they are highly influenced by the characteristics of the training data. Consequently, careful consideration is necessary when training such models to avoid overfitting to overly specific sequence characteristics and learning spurious correlations that do not reflect the true biological patterns. To accurately assess a model’s ability to generalize to sequences outside of its training scope, it is critical to evaluate it on sequences not too similar to those in the training data [38, 39]. We performed a taxonomy-based partitioning at the whole-genome level to split data into training, validation, and test sets using family-level taxonomy as the basis for partitioning, such that no pair of organisms belonging to the same family ends up in different partitions. This was done with the following procedure:

1. A selection of archaeal and bacterial organisms was chosen for the test set (see Supplementary Table A3). These were selected to span a broad range in genomic GC-contents and because they have been widely used in previous gene prediction benchmark studies [2, 5, 6, 11, 24].
2. All organisms belonging to the same family as one of the organisms selected for the test set were added to the test partition.
3. All remaining organisms were grouped by family, and the average GC-content for each family was calculated.
4. Archaeal and bacterial families were processed separately. Within each domain, families were sorted by mean GC-content and sampled systematically to select families for the validation set, ensuring the validation set to contain *≈* 10% of the remaining organisms per domain and good coverage across the GC-content range.
5. All organisms from the remaining families (*≈* 90%) were assigned to the training set.

Following this procedure, the training set consisted of the sequences from 813 genomes, the validation set of sequences from 97 genomes, and the test set of sequences from 215 genomes (see Supplementary Table A4 for summary statistics). Family-level partitioning does, however, not rule out similarity between training and test sequences in conserved regions shared across broader taxonomic groups, which we therefore assessed separately (Sections 2.5.2 and 3.7). The final datasets used for model development consist of approximately 20M sequences in the training partition and 2.3M sequences in the validation partition. The test datasets vary in the number of sequences due to the simulation of sequence lengths (see Section 2.5.1). Supplementary Tables A5-A12 report the number of reads for each genome in each dataset, and hierarchical taxonomic trees showing the distribution of partitions across the archaeal and bacterial domain are shown in Supplementary Figures A2 and A3.

### 2.4 DeepCDS Modeling

#### 2.4.1 Model architecture overview

DeepCDS is an end-to-end model that takes an arbitrary DNA sequence as input and outputs both complete and partial CDS coordinates. The model runs independently on the input sequence and its reverse complement to detect CDSs on both strands (Figure 2). DeepCDS was developed for and evaluated on short sequences (60–1,000 bp), either without sequencing errors or with simulated Illumina-like error profiles (the latter evaluated at up to 300 bp). Sequences longer than the 300 bp training window are processed using a sliding window approach (Supplementary Note A1.1). DeepCDS can be divided into three overall stages: *(i)* individual encoding of each RF into a contextual representation (Figure 2B), *(ii)* joint decoding of a biologically constrained label sequence across all three RFs using a linear-chain conditional random field (CRF), and *(iii)* converting decoded labels into CDS coordinates, start codon positions, and stop codon positions.

**Fig. 2:**
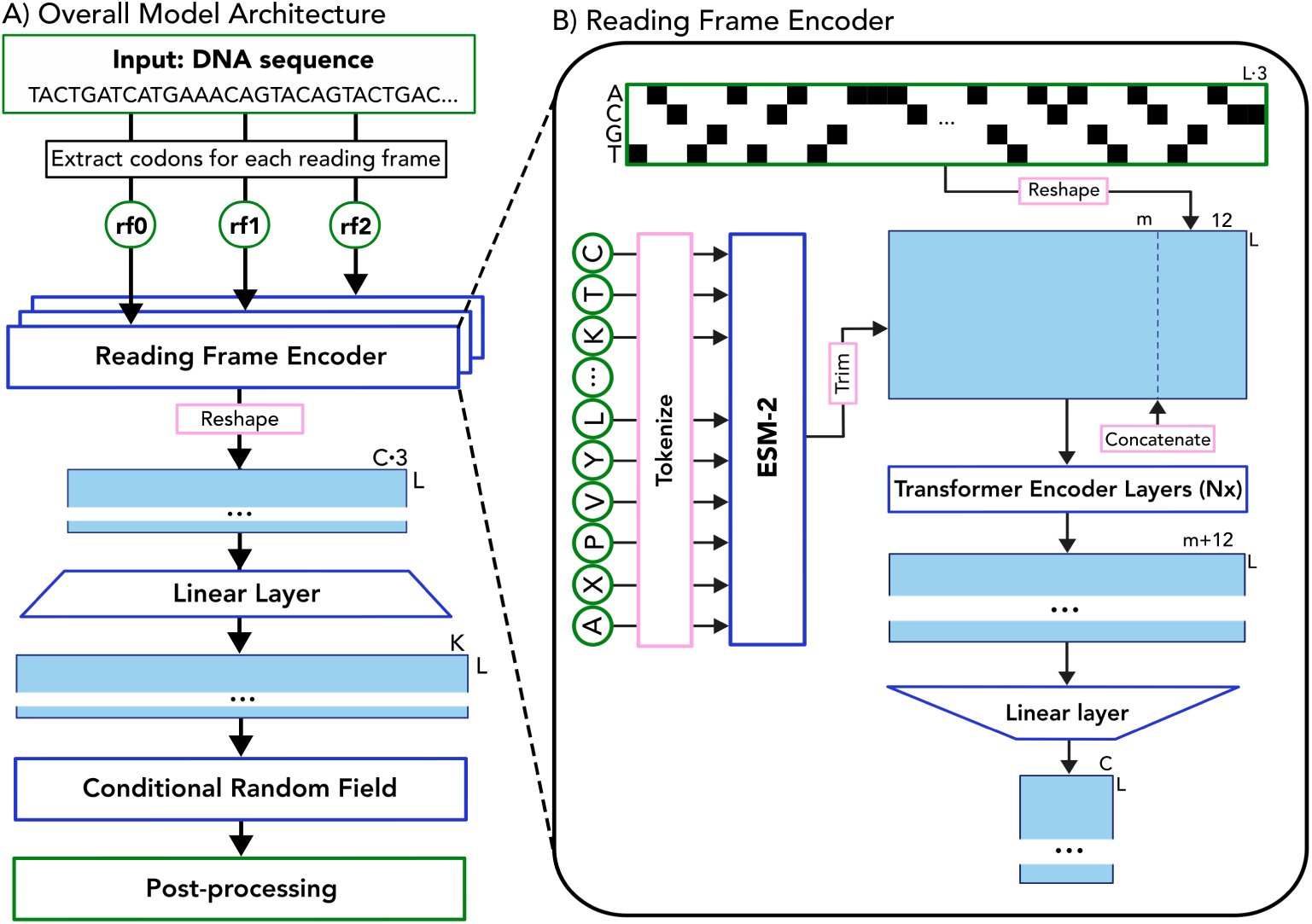
DeepCDS model architecture. A) End-to-end model architecture. Given an input DNA sequence, DeepCDS is run independently on the input sequence and its reverse complement to detect CDSs on both strands. For each sequence, the codon sequence for each of the three RFs is processed independently through a shared Reading Frame Encoder (weights shared across frames). The Reading Frame Encoder produces per-codon logits over label classes *C* at each codon position *l ∈ L*. Log- its from all three frames are concatenated and projected onto a shared label space of dimension *K*, corresponding to biologically valid joint label combinations across frames. The projected logits are decoded by a linear-chain CRF, yielding the most probable shared label sequence. During postprocessing, shared labels are mapped back to per-frame label sequences and converted into CDS coordinates, including start codons, stop codons, and indel error positions. B) The Reading Frame Encoder. Each RF is processed through two parallel branches: (*i*) one-hot encoding of nucleotides, reshaped into 12-dimensional codon representations *∈* R*^L,^*^12^, and (*ii*) translation of the codon sequence into amino acids, which are embedded and contextualized through the pretrained pLM ESM-2 [27]. The classification (CLS) and end-of-sequence (EOS) tokens are removed, and the resulting amino acid embeddings are concatenated with the codon representations to combine nucleotide-level and amino acid-level context. The concatenated embeddings are passed through stacked transformer encoder layers (optimized as a hyperparameter, see Supplementary Table A13) and projected to percodon logits over *C* codon label classes.

##### Encoding reading frames into contextual representations

Given an input DNA sequence, the model encodes each RF independently using the Reading Frame Encoder (Figure 2B). The weights are shared across all three RFs, enforcing the assumption that the same biological features are informative for each frame, while frame-specific representations emerge from the distinct input codon sequences. As RFs are defined relative to the start of the input sequence rather than to the CDS, a CDS in a randomly sampled read is equally likely to occur in any RF, leaving no frame-specific signal for separate encoders to specialize on. RF identity is retained in the downstream joint label space, allowing the CRF to model frame-specific transitions. Each RF is encoded through two parallel branches that together capture both nucleotide-level and amino acid-level context:

*(i) a codon representation.* Nucleotides are one-hot encoded (A = [1, 0, 0, 0], C = [0, 1, 0, 0], G = [0, 0, 1, 0], T = [0, 0, 0, 1], and N = [0, 0, 0, 0]), and three consecutive nucleotides forming a codon were concatenated to create 12-dimensional codon representations *∈* R*^L,^*^12^ to capture the nucleotide-level context of each codon, where *L* is the number of codons in a RF.
*(ii) a contextualized amino acid embedding.* The RF is translated to amino acids using the standard prokaryotic genetic code (NCBI genetic code 11). The amino acid sequence is tokenized and embedded into vectors *∈* R*^L^*^+2*,m*^ (where the +2 accounts for ESM-2’s special tokens), which are processed through the pretrained pLM ESM-2 [27] with 8 million parameters. Stop codons (TAA, TAG, and TGA) are encoded as X. Codons containing an unknown nucleotide are encoded as unknown tokens (
*<*unk*>*). We extract the final hidden layer representation for each amino acid token (excluding special tokens) to obtain a contextualized representation of the sequence *∈* R*^L,m^*, where *m* = 320 is ESM-2’s hidden dimension.

For each codon position, the nucleotide-level codon representation vector and the learned amino acid embedding are concatenated to combine nucleotide-level codon information with amino acid-level context, yielding vector representations *∈* R*^L,m^*^+12^.

The concatenated embeddings are passed through stacked transformer encoder layers, each consisting of standard multi-head self-attention followed by a position-wise feedforward network. The feedforward sublayer within each transformer encoder layer uses a hidden dimension of 4 *×* (m + 12), following the standard expansion factor from the original transformer architecture [40]. The number of layers, attention heads per layer, and the activation function within each feedforward network were optimized as hyperparameters for each model (Supplementary Table A13). No additional positional encoding was added to the transformer encoder layers, as positional information is implicitly carried by the pLM embeddings. The transformer layers are followed by a linear layer which projects each codon position to class logits *∈* R*^L,|C|^*, where *|C|* = 6 for the model trained on the dataset with indel and substitution errors, and *|C|* = 4 for the models trained with substitution errors only and without errors, respectively (Figure 1, “Data preprocessing” and Supplementary Figure A4).

The two input representations are intended to be complementary: the one-hot codon encoding retains nucleotide-level information such as codon usage and synonymous codon choice, while the ESM-2 embeddings provide a learned prior over protein-like sequence features from pretraining on millions of protein sequences, which may be particularly informative when only limited sequence context is available. The apparent information loss in translating nucleotide sequence to amino acid sequence may furthermore be beneficial for generalization to remotely related organisms, since codon biases tend to be species-specific [41]. Since all three RFs are translated and encoded, the correct translation of each frame remains available to the model even when an indel error shifts the true RF.

##### Joint structured decoding across reading frames

To model inter-frame dependencies, logits from all three RFs were concatenated to form a joint representation *∈* R*^L,^*^3*·|C|*^ (Figure 2A). A linear layer projects this to a shared label space *∈* R*^L,K^*, where *K* is the number of biologically valid label state combinations across RFs, which was determined empirically from the training datasets. Specifically, for the *|C|* = 4 model variants, *K* = 33, and for the *|C|* = 6 model variants, *K* = 69. This joint label space encourages consistency across RFs, particularly at label transitions between non-coding and coding boundaries. This is especially critical in the presence of indel errors, where a frameshift in one RF is expected to be reflected by a corresponding transition in the affected frame.

In early stages of this work, we experimented with model versions operating independently on each RF. These experiments showed poor agreement in predicted transition positions across RFs. Each shared label, *k_n_* = [*c_n,_*_rf0_ *, c_n,_*_rf1_ *, c_n,_*_rf2_], represents the joint state at codon position *l* across all three RFs, where rf_0_, rf_1_, and rf_2_ are offset by 0, 1, and 2 nucleotides relative to the sequence start, respectively.

We employ a linear-chain CRF to decode the label sequence [42], implemented using the Python package pytorch-crf. Given per-position logits **H** = (**h**_1_*, . . .,* **h***_L_*) where **h***_l_ ∈* R*^K^*, the CRF models the conditional probability of a label sequence **y** = (*y*_1_*, . . ., y_L_*) as:

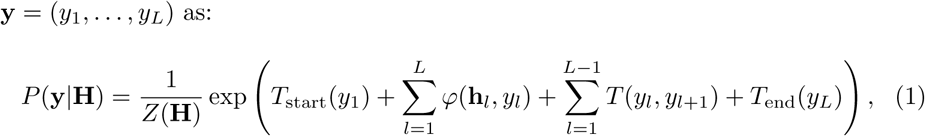

where *φ*(**h***_l_, y_l_*) is the emission score for assigning label *y_l_* at codon position *l*, *T* (*y_l_, y_l_*_+1_) is the learned transition score between consecutive labels, and *T*_start_(*y*_1_) and *T*_end_(*y_L_*) capture the prior likelihood of a sequence beginning and ending with labels *y*_1_ and *y_L_*, respectively. *Z*(**H**) is the partition function that sums over all possible label sequences to ensure proper normalization.

By learning transition scores, the CRF strongly discourages biologically invalid transitions (in practice, at most 6*·*10*^−^*^5^% of test sequences contained such a transition, see Supplementary Note A1.2). In early model versions without the CRF, we observed noisy regions where stretches of a few codons frequently flipped between non-coding (*c* = 0) and CDS (*c* = 1) labels without passing through transition labels (start codon, stop codon, or indel error-induced transition labels, *c ∈ {*2, 3, 4, 5*}*). With the CRF, a coding region is, for example, strongly discouraged from transitioning directly to a non-coding region without passing through a stop codon or indel error event. The CRF thus constitutes an integral part of the model output rather than an auxiliary module: by producing a single, biologically consistent label sequence, it allows CDS coordinates, start and stop codon positions, and indel error positions to be derived directly from the decoded labels, without extensive rule-based post-processing to resolve invalid label transitions or noisy regions with frequent label flips. During inference, the most probable label sequence **ŷ** = arg max**_y_** *P* (**y***|***H**) is decoded using the Viterbi algorithm.

##### Mapping decoded labels to CDS coordinates

The decoded shared labels are mapped back to their corresponding per-RF label sequences, providing the per-codon class labels for each of the three RFs (Figure 1, “Post-processing”). The label sequence for each RF is then converted into corresponding CDS coordinates, including start codon, stop codon, and indel position information.

#### 2.4.2 Training procedure

We trained three variants of the model: one trained on the sequences with both substitution and indel errors (**DeepCDS S+I**), one with only substitution errors (**DeepCDS S**), and one without sequencing errors (**DeepCDS N**). For each model, we used a fixed batch size of 32 and optimized a range of hyperparameters using Optuna [43] (see Supplementary Tables A13 for optimized hyperparameters and A14 for training settings). In the last stage of hyperparameter selection, we filtered away configurations where training did not converge properly.

All the models were trained to minimize the negative log-likelihood of the true label sequences [42] (See Supplementary Note A1.3). We employed the AdamW optimizer [44] with weight decay of 0.01, implemented early stopping [45] by monitoring the negative log-likelihood loss on the validation set, and applied gradient clipping with a maximum norm of 1.0 to prevent exploding gradients.

During training, we kept all parameters of ESM-2 frozen for the first 3 million sequences to allow the parameters learned from scratch to stabilize before beginning to fine-tune ESM-2. Afterwards, the last half of the ESM-2 layers were fine-tuned at a lower learning rate than that used for the weights trained from scratch. To prevent destabilization of the pretrained weights, we employed a linear warmup over 32,000 training sequences, starting at 10% of the base ESM-2 learning rate. We employed the PyTorch ReduceLROnPlateau scheduler to monitor validation loss and reduced both learning rates by 30% when no improvement was observed for 6 consecutive validations.

The transition matrix in the CRF was initialized by assigning all biologically invalid transitions a transition score of *−*10, which multiplies the unnormalized probability of any label sequence containing such a transition by a factor exp(*−*10) *≈* 4.5 *·* 10*^−^*^5^, making label sequences containing such a transition highly unlikely. Specifically, transitions between non-coding and coding states are invalid without passing through a start codon, stop codon, or indel error-induced frameshift state (Supplementary Figure A4). In contrast, the frequent transitions were initialized with a neutral transition score of 0 (exp(0) = 1), which included all shared label states where each RF had a non-coding*→*non-coding transition, or where one RF had a coding*→*coding transition and the remaining two RFs had a non-coding*→*non-coding transition. We define the remaining valid transitions as “infrequent valid” transitions, and the initialization value of these was optimized as a hyperparameter (Supplementary Table A13). From this initialization scheme, the transition scores were learned during training (Supplementary Figure A5).

All sequences were translated using the standard prokaryotic genetic code, even those from the minority of genomes using an alternative code (NCBI genetic code 4, where TGA is translated as W instead of a stop codon). Since DeepCDS targets sequences of unknown origin, we opted for a single universal encoding rather than requiring translation table specification as input. We tested the robustness to this on 3 genomes from the family *Mycoplasmataceae* which uses an alternative genetic code (Section 3.4).

For validation, we sampled 25% of sequences in the full validation set (*≈* 575,000 sequences), following the original distribution of sequences per genome and labeled sequence types. The DeepCDS models were trained on an NVIDIA H100 GPU, where DeepCDS N, DeepCDS S, and DeepCDS S+I required 91, 90, and 152 GPU hours to train, respectively. DeepCDS N and DeepCDS S each contain *≈* 12.8M parameters, and DeepCDS S+I contains *≈* 15.5M parameters. In each case, *≈* 8 M originates from ESM-2.

To examine the effect of genomic diversity on model performance, we trained the DeepCDS S+I models on progressively smaller subsets of the full training data, following an identical training procedure as described above. Starting from the full set of 813 genomes, we constructed subsets of 400, 200, and 100 genomes, each sampled to preserve the taxonomic family distribution of the full training set. In cases where a family was represented by a single genome, that genome was retained regardless of subset size. The results are shown in Supplementary Table A15.

#### 2.4.3 Input feature ablations

To assess the relative contributions of the two input features (pLM embeddings and one-hot codon encoding) used in the full DeepCDS model, we trained ablation versions processing only one feature type as input. The models trained solely on the one-hot codon encodings are referred to as **DeepCDS (Codon)**, and the models trained solely on the pLM embeddings are referred to as **DeepCDS (pLM)**. For the (Codon) ablation, each 12-dimensional vector encoding a codon is projected to a 332-dimensional hidden representation, and a standard sinusoidal positional encoding is added before being fed to the transformer encoder layers in the Reading Frame Encoder (Figure 2B) to match the dimension of the full DeepCDS model. For the (pLM) ablation, each amino acid is represented as a 320-dimensional vector determined by the ESM-2 8M embedding dimensionality, which was also used as the transformer encoder hidden size. All models were trained with identical hyperparameters and procedures as described in Section 2.4.2.

### 2.5 Evaluation

#### 2.5.1 Test sets

Since the sequencing error rate of a given dataset may not be known a priori, we evaluated the robustness of the trained models across test sets spanning a range of error rates, from error-free sequences to a stress-test condition exceeding typical Illumina error rates. Additionally, we produced test sets of varying sequence lengths. For the test sets simulated with sequencing errors, we used Mason’s default Illumina sequencing error profile. We constructed test sets across a range of read lengths, including the standard Illumina read lengths of 100, 150, and 300 bp, as well as shorter lengths of 60 and 75 bp, to assess robustness to reduced sequence context. Longer reads with sequencing errors were not simulated, as these would not reflect Illumina sequencing output. For each read length, we simulated three error rate conditions, keeping the ratio between substitution and indel errors constant:

- Typical Illumina error rates (**Typical-error**): a substitution error rate of 4 *·* 10*^−^*^3^ (0.4%) per base and indel error rate of 1 *·* 10*^−^*^5^ (0.001%) per base.
- High-range Illumina error rates (**High-error**): a substitution error rate of 1 *·* 10*^−^*^2^ (1%) per base and indel error rate of 2.5 *·* 10*^−^*^5^ (0.0025%) per base.
- Stress testing error rates (**Stress-error**): a substitution error rate of 3 *·* 10*^−^*^2^ (3%) per base and indel error rate of 7.5 *·* 10*^−^*^5^ (0.0075%) per base.

Read lengths and error rates vary across Illumina platforms, and while simulated error profiles cannot replicate empirical sequencing errors, evaluating a range of error rates provides a reasonable approximation of realistic sequencing scenarios. The empirical distributions of error rates obtained for each test set are shown in Supplementary Figure A6.

We also generated test sets without sequencing errors at the same lengths with Mason, as well as at 700 and 1,000 bp, referred to as the **error-free** test sets. The 700 and 1,000 bp test sets were included to assess performance on sequences exceeding the 300bp training window, which are processed using a sliding window approach (Supplementary Note A1.1), and to evaluate the potential applicability of DeepCDS to longer error-free sequences, such as short contigs from fragmented metagenomic assemblies.

The test sets were simulated based on the 215 genomes in the test partition, of which 212 follow the standard prokaryotic genetic code. Furthermore, we included *M. capricolum* and 2 further genomes belonging to the family *Mycoplasmataceae* to investigate model behavior on genome representatives using an alternative genetic code. Family-level statistics are shown in Supplementary Table A16.

##### Test sets with alternative short read simulator

To assess whether model performance was robust to simulator-specific biases, we performed an additional benchmark on reads simulated with art modern (v1.4.0) [46] on the 215 test genomes. art modern is an accelerated re-implementation of the well-established short read simulator ART [47]. We simulated three test sets using the built-in HiSeq2500 (150bp), NextSeq500 (150bp), and MiSeq v3 (300bp) quality profiles, with the remaining settings left at their defaults (See Supplementary Figure A7).

##### *Realistic simulated short-read human gut microbiome metagenomic test* set

A challenge arising in the development of a CDS predictor aimed at metagenomic short read contexts is that validation on an actual metagenomic dataset is difficult, due to the lack of such benchmark datasets with pre-defined ground truth CDS annotations not originating from another ab initio gene prediction program like Prodigal [5, 48]. However, to assess model performance in a more realistic metagenomic setting, compared to measuring performance on the whole test genomes as described above, we additionally downloaded sample 0 of the simulated short-read shotgun metagenomic human gut microbiome from the Critical Assessment of Metagenome Interpretation (CAMI) III challenge “Toy Longitudinal Human Gut” [49–51]. In the ”Longitudinal Human Gut” version, the genomes that sequences are simulated from are not publicly available, and so we cannot perform proper validation, as no ground truth can be defined. In the “Toy” version of the dataset, reads are simulated from publicly available genomes, allowing us to map reads accurately to their respective genomes of origin while retaining the characteristics of a true metagenomics dataset. The full dataset consists of 16,633,513 paired-end reads of length 150 bp from 309 distinct genomes.

To avoid overestimating performance due to overlap with the data used for model development, we removed all genomes from the CAMI III sample that were identical to, or had an average nucleotide identity (ANI) of at least 95%, to any genome in the training or validation sets, corresponding to the commonly used species boundary for prokaryotic genomes [52]. The ANI was computed with skani dist [53] (v0.4.0) using default parameters. This removed 43 genomes (2 identical and 41 belonging to a species represented in the training or validation sets), corresponding to 17.37% of the reads.

We separated the paired-end reads to map each individually to the genomic coordinates of the genome the read originated from and kept track of stored indel positions, which were labeled as described in Section 2.2. For each contig within each genome assembly (NCBI accession), we collected both RefSeq and GenBank annotations from NCBI’s Nucleotide (nuccore) database, with GFF3 annotation files retrieved via NCBI’s Sequence Viewer. We discarded reads overlapping with CDS annotations including a pseudogene (“pseudo=true” or “pseudogene”), partial (“partial=true”), or programmed frameshift (“note=programmed frameshift”) tag. Reads overlapping with CDS annotations tagged as “product=hypothetical protein” or “ab initio prediction” were marked. We constructed three versions of this test set with varying degrees of annotation certainty, in each case retaining the exclusion of reads overlapping pseudogene-, partial-, or frameshift-tagged CDS annotations described above: (i) a baseline test set, where reads overlapping with a hypothetical-protein or ab initiotagged CDS annotation were retained and treated as correctly labeled; (ii) a test set excluding all reads overlapping with either tag; and (iii) the baseline test set, additionally excluding reads originating from contigs with no annotations at all. Detailed summary statistics of the processed test sets are presented in Supplementary Table A17.

As mentioned in Section 2.2, we removed reads overlapping with CDSs annotated as hypothetical proteins as a conservative measure to maximize label reliability during training. However, hypothetical proteins frequently represent genuine protein-coding genes not yet functionally characterized. For this reason, we measured the sensitivity on the genome overlap-reduced prokaryotic CDS fragments annotated as a hypothetical protein that were not explicitly tagged as ab initio predictions (452,202 reads, corresponding to 4.65% of all reads in this set with an annotated prokaryotic CDS). As a reference, we also extracted reads overlapping CDS annotations explicitly annotated with a molecular function (“go function” attribute). As false positives (FPs) cannot be meaningfully assigned to a subset of the ground truth CDSs, performance on these subsets was evaluated in terms of sensitivity only, and is presented in Supplementary Figure A8.

##### *Real Illumina-sequenced gut metagenome from the Danish MetaHIT* cohort

To provide a proof of concept on real, non-simulated short read data, which, unlike simulated reads, can contain artifacts such as chimeric reads, cross-contamination and PCR duplicates, we used a human gut metagenome sample from the Danish cohort of the MetaHIT (Metagenomics of the Human Intestinal Tract) project [54], obtained from the European Nucleotide Archive (project PRJEB4336, sample MH0161, run ERR321493). The sample was sequenced on an Illumina HiSeq 2000 platform as paired-end 2*×*90bp reads (33,282,106 read pairs). The reads were quality- and adapter-trimmed with fastp [55] (v1.4.0, default settings), leaving 29,201,473 pairs.

We removed potential host contamination by mapping reads to the human reference genome (GRCh38) with Bowtie 2 [56] (v2.5.5, default settings) and keeping only pairs in which neither mate mapped. This removed 1,435 additional read pairs. The mates were then treated as separate reads, resulting in *≈*58.4M reads longer than 60 bp, 99.6% of which were full length (90 bp; Supplementary Figure A9).

#### 2.5.2 Homology-aware assessment at the nucleotide- and protein-level

As the data partitioning is taxonomy-based (Section 2.3), test genomes may still share homologous genes with the model development (training and validation) genomes. This could cause the performance of DeepCDS to partly reflect shared homologous information rather than genuine cross-species generalization. To assess this, we stratified the error-free 300bp test set, based on sequence identity to the error-free training and validation datasets, and measured performance within each identity bin. We chose the 300bp error-free test set for this analysis because it is the setting without sequencing noise and with the same sequence length as the training data, and thus the most direct test of the influence of sequence similarity.

We conducted the homology-stratification at both the protein-level and the nucleotide-level. At the protein-level, all labeled CDS fragments longer than 60bp from the training, validation and test sequences were translated into their corresponding amino acid sequences. Each test fragment was then searched against all training and validation fragments with MMseqs2 [57] (v18.8cc5c, mmseqs search -s 7.5 -e 10 --min-aln-len 20). For the nucleotide-level test, the complete test sequences, including non-coding regions, were searched against all training and validation sequences (mmseqs search --search-type 3 --strand 1 -s 7.5 -e 10 --min-aln-len 60). Only the given strand was searched, as both strands are simulated separately and each orientation thus enters the training and test data as a sequence of its own (Section 2.2). Each test fragment or sequence was assigned the sequence identity of its best hit by bitscore.

Because the sequences are short and the model development data comprise a large number of sequences, similarity may arise by chance rather than from shared ancestry or conservation. However, the model could still benefit from such similarity during training. We therefore used a permissive E-value threshold of 10, such that similarities that are likely to have arisen by chance rather than by shared homology are also kept. Consequently, a test sequence is only assigned to the “no hit” category if MMseqs2 finds no alignment of at least 60 nucleotides (or 20 amino acids in the protein-level test) with *E*-value *≤* 10 compared to any training or validation sequence, making this category a stringent set of sequences novel relative to the training data. Complementary results for the standard E-value threshold of 10*^−^*^3^ are reported in Supplementary Figure A10.

Protein-level fragments were grouped into identity bins (no hit, *<* 30%, 30–50%, 50–70%, 70–90%, *≥* 90%), and CDS-level sensitivity was computed per bin (see Section 2.5.4). This is the primary analysis, as protein-level search detects more distant homology due to the redundancy of the genetic code [58], and matches DeepCDS’ use of pLM representations. Because the nucleotide-level test also covers non-coding regions, it provides a complementary analysis measuring sequence similarity across entire sequences from the model development and test set. CDS-level precision, sensitivity and F1 score were computed per bin (no hit, *<* 80%, 80–90%, 90–95%, 95–99%, *≥* 99%).

#### 2.5.3 Models in benchmark

We compared the performance of the three DeepCDS model variants (N, S, and S+I) to that of MetaProdigal (running Prodigal v2.6.3 in the mode -p meta) [13] and FGS [6] using the reimplemented version from 2022, FragGeneScanRs (v1.1.0) [12], which has been shown to produce identical results but at a substantially shorter computational time. MetaProdigal does not take into account sequencing errors, and for this reason we ran the same model on all test sets. In contrast, FGS provides both a model variant for predicting on complete genomic sequences, as well as variants that integrate sequencing error models. For the test sets without sequencing errors, we used the variant for predicting complete genomic sequences, and for the test sets with sequencing errors, we ran both the variant for complete genomic sequences, and variants for Illumina sequencing reads of error rate 0.5% and 1.0%, respectively. These are referred to as **FGS (Complete)**, **FGS (0**.**5**% **error rate)**, and **FGS (1**.**0**% **error rate)**.

We additionally compared performance to MetaGeneMark-2 [26] (GeneMark.hmm-2 v1.23). The model parameters of MetaGeneMark-2 fix its minimum gene length, so that it cannot predict CDSs shorter than 90 bp. Because DeepCDS is aimed at detecting short CDS fragments, our main benchmark retains all CDSs longer than 60 bp (Section 2.5.4). This would penalize MetaGeneMark-2 for CDSs it cannot predict by design. For this reason, we evaluated MetaGeneMark-2 in a separate benchmark, in which both the test set CDSs and the predictions of all models were restricted to CDSs of at least 90 bp. This was based on the 300bp versions of all four test sets (error-free, typical-error, high-error and stress-error), using both exact coordinate matching and the less strict overlap criteria described in Section 2.5.4. Because MetaGeneMark-2 cannot predict CDSs shorter than 90 bp, and because it was outperformed by MetaProdigal and DeepCDS throughout, it is not included in the main benchmark (see Section 3.1).

#### 2.5.4 Evaluation Metrics

##### CDS-level validation

For overall assessment, we evaluated the models in their ability to predict the correct CDS regions. We defined a True Positive (TP) as a predicted CDS with exact coordinate match (identical start and stop positions) to the test set CDS. False Positives (FP) were defined as predicted CDS coordinate ranges not present in the test set, and False Negatives (FN) were defined as CDS coordinate ranges in the test set that were not predicted. This strict criterion means that predictions with shifted boundaries, whether by as little as a single codon or due to RF disagreement, contribute to both FP and FN counts. For the benchmark, we discarded all CDS ranges in the test set shorter than 61 bps, which is the minimum length threshold of FGS, and removed predicted CDS ranges from MetaProdigal and the DeepCDS models shorter than this threshold to ensure a fair comparison across all models (MetaProdigal predicts down to 60 bps). Although we consider exact coordinate matching the most biologically meaningful criterion for proper evaluation, we also tested less strict overlap criteria. This was measured as the Intersection over Union (IoU), defined as the fraction of nucleotide positions that are shared between an actual CDS in the test set and a predicted CDS, while requiring agreement in RF. These results are reported in Supplementary Tables A18–A25.

Based on the above-defined TP, FP, and FN counts, we calculated the precision (fraction of correctly predicted CDS ranges out of all predicted CDS ranges), the sensitivity (fraction of correctly predicted CDS ranges out of all actual CDS ranges), and the F1 score (harmonic mean between precision and sensitivity):

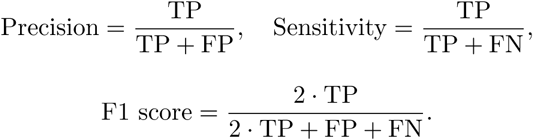

##### Codon-level validation

We calculated codon-level performance using the Matthews Correlation Coefficient (MCC), defined as:

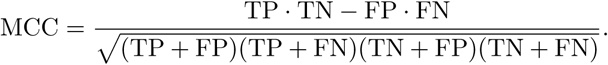

TPs are codons belonging to a CDS and predicted as CDS, FPs are non-coding codons predicted as CDS, FNs are CDS codons predicted as non-coding, and True Negatives (TN) are non-coding codons predicted as non-coding. Codons from CDS fragments shorter than 61 bps were masked to ensure a fair comparison to FGS and MetaProdigal. The MCC was chosen because it is easily interpretable, with a well-defined value of 0 corresponding to random classification.

##### Start and stop codon identification

To assess each model’s ability to correctly identify start and stop codons, we defined predicted start codon positions as the 5’ boundaries of predicted CDS regions, and stop codon positions as the 3’ boundaries. We refer to start codon identification as predicting the annotated start codon of a CDS, which approximates, but may not always correspond to, the true translation initiation site (see Section 3.3, “Model behavior on CDSs with conflicting annotated start positions”). We assessed predictions separately for each RF. From the test sets, we identified all sequences containing annotated start or stop codons, excluding CDS boundaries caused by frameshift-inducing indel errors. We further excluded cases where a start or stop codon occurred at the first or last possible position in a sequence for a given RF (positions 1-3 at the 5’ end or the final 3 positions at the 3’ end), as FGS does not explicitly annotate start or stop codons, making it ambiguous whether such positions represent true start or stop codon predictions or simply CDS regions extending to the sequence boundary (see Supplementary Table A26 for DeepCDS and MetaProdigal’s performance on canonical- and non-canonical start codon identification, including counts at sequence borders). We calculated the F1 score based on defining TPs as start/stop codon positions present in both the test set and the model’s predictions, FPs as start/stop codon positions predicted by the model but absent from the test set, and FNs as start/stop codon positions present in the test set but not predicted by the model.

##### Indel detection and localization

For the models trained on indel error-containing reads (i.e., FGS 0.5% error rate, FGS 1.0% error rate, and DeepCDS S+I), we evaluated indel detection performance across all test sets with sequencing errors (typical-error, high-error, and stress-error). TPs, FPs, and FNs were defined at the indel level; within each read, predicted indels were matched one-to-one to true indels in order of increasing distance between their positions. A matched predicted indel located within a positional tolerance of *±k* bp of its true indel error site was counted as a TP. Predicted indels without a TP match, including those on reads without an actual indel error, were counted as FPs, and true indels without a TP match were counted as FNs. Precision, sensitivity, and F1 score were derived from these counts at the genome level using *k* = 10 (Supplementary Figure A11), and from counts aggregated across the test genomes for *k ∈ {*0, 1, 3, 5, 10, 25*}* (Supplementary Figure A12). To assess positional accuracy, we computed the mean absolute error (MAE) per genome as the mean absolute distance between the positions of all matched true and predicted indels, independent of the positional tolerance (Supplementary Figure A11). Note that precision, sensitivity, and F1 score reflect positional accuracy only up to the chosen tolerance; finer positional accuracy is captured by the MAE.

##### Uncertainty of aggregated test performance: genome-level cluster bootstrap

For performance metrics calculated as aggregated values across several genomes in the test set (i.e., not boxplots), TP, FP, FN, and TN counts are summed across sequences from each genome, so that each predicted or annotated feature contributes equally irrespective of its genome of origin. Because sequences from the same genome are not independent observations, a sequence-level interval would understate the true uncertainty. We therefore estimated uncertainty using a genome-level cluster bootstrap [59, 60]. In each of 1,000 replicates, we resampled the full set of test genomes with replacement, summed the resampled genomes’ raw TP, FP, FN, and TN counts, and recomputed each metric from those totals. We report the standard deviation across the 1,000 replicate scores as the standard error (*±*); this is a bootstrap standard error rather than a percentile confidence interval.

##### *Protein-level support of predicted CDSs from a real gut metagenome* dataset

Obtaining ground-truth CDS annotations for real metagenomic samples is generally difficult, as the genome of origin of each read is unknown, many community members lack reference genomes, and available annotations may be based on computational gene prediction. For the MetaHIT sample (Section 2.5.1, “Real Illumina-sequenced gut metagenome from the Danish MetaHIT cohort”), we therefore used similarity to known proteins as a proxy for the support of each predicted CDS. Each tool predicted CDSs on both strands of each read (DeepCDS was applied to each read and its reverse complement), and only predicted CDSs longer than 60 bp were retained, so that all tools were compared on CDSs of the same minimum length. Each CDS was translated in its predicted frame and strand using the standard prokaryotic genetic code, except for CDSs predicted by MetaProdigal, which were translated with the genetic code of the MetaProdigal model chosen for the read. DeepCDS S+I CDSs containing a predicted indel were translated segment by segment, and the segments were joined into one protein. For FGS (0.5% error rate) and FGS (1.0% error rate), each CDS was reconstructed from its reported coordinates and FGS’s predicted indel positions (removing inserted bases and adding an unknown base at each deletion) before translation. All translated CDSs were searched against UniRef90 [61] (release 2026 03, 121.5M sequences) with DIAMOND [62] (v2.2.8, blastp --sensitive --max-target-seqs 1). A CDS was considered supported if it had a UniRef90 hit with an E-value *≤* 10*^−^*^3^ (default threshold). As a negative control, about 20,000 randomly selected predicted CDSs per tool (140,672 in total) were searched in the same way after shuffling the order of their amino acids, which keeps their length and composition but removes any real homology. The results are reported in Section 3.6.

## 3 Results

### 3.1 DeepCDS consistently outperforms well-established short read coding sequence predictors

We evaluated DeepCDS and the benchmark models on test sets of sequences from the 215 test genomes across a range of sequence lengths and sequencing error rates (Methods Section 2.5). Unless otherwise stated, the results below reflect performance on the 212 genomes that follow the standard prokaryotic genetic code (see Results Section 3.4 for an assessment on the remaining three genomes with an alternative genetic code).

Across all tested conditions, DeepCDS consistently outperformed the FGS models and MetaProdigal at both the CDS- and codon-level (Figure 3, Table 1 and Supplementary Table A27). The performance advantage increased with sequencing error rate and sequence length, with all pairwise comparisons remaining statistically significant after multiple testing correction (adjusted p-value *<* 0.05 for all tested conditions, see Supplementary Note A1.4). The CDS-level results presented in the sections below are based exclusively on exact coordinate matching (defined as an IoU of 1, see Section 2.5.4). While we consider exact coordinate matching the most suitable approach to assess performance, less strict overlap criteria for defining a predicted CDS as a TP provide an important complementary view on the ability to identify approximate CDS regions and may be more robust in the presence of uncertain annotated CDS positions. Using IoU thresholds of 0.8 and 0.5 yields the same model ranking as exact coordinate matching across all error rate conditions, with the DeepCDS models achieving the highest performance. However, the differences in performance between models decrease as the overlap criterion becomes less strict (Supplementary Tables A18–A25).

**Fig. 3:**
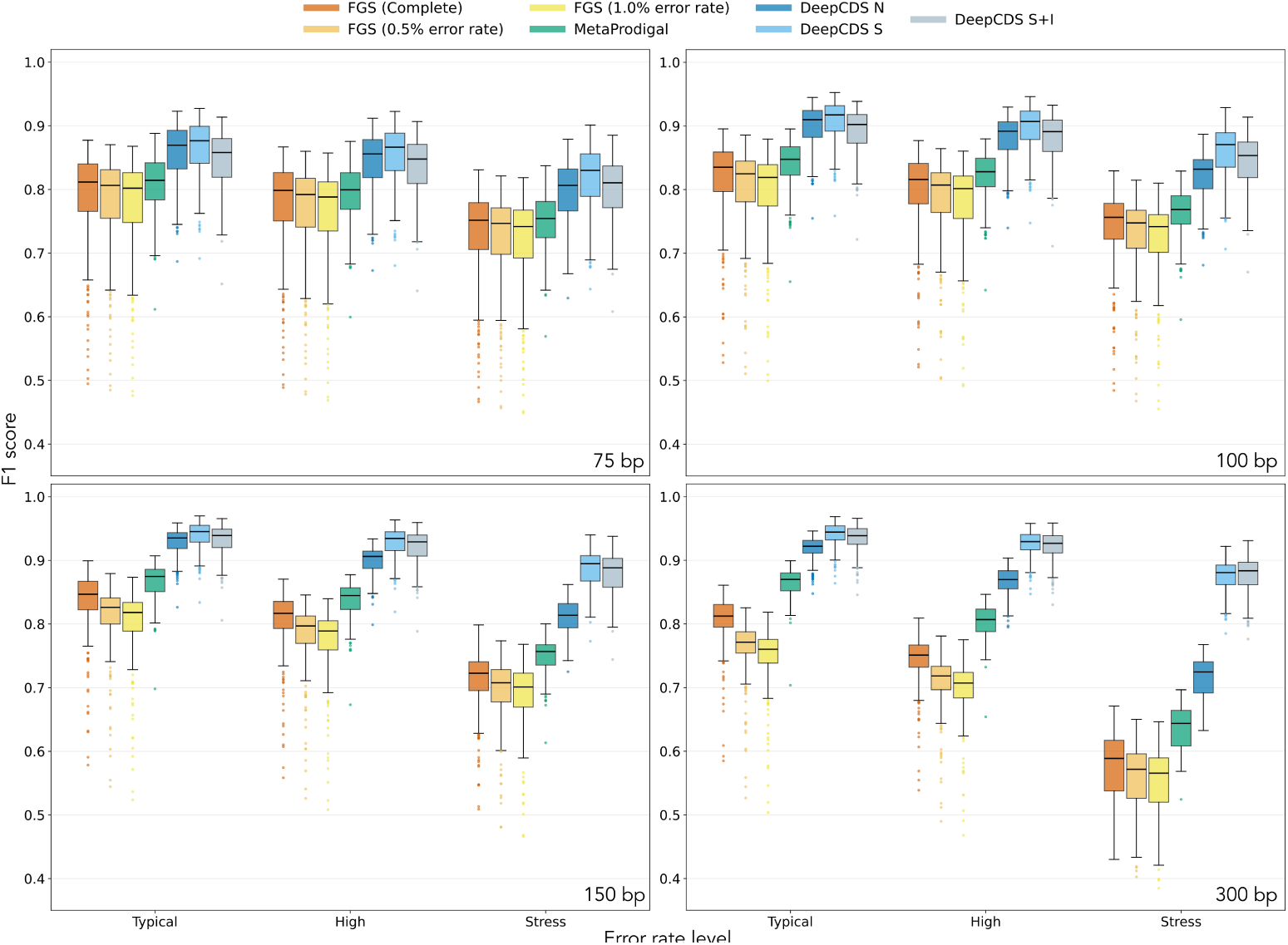
CDS-level performance across error rate and sequence length conditions at the genome-level. Boxplots visualizing the CDS-level F1 score for the 212 genomes that follow the standard prokaryotic genetic code across test sets of varying sequence length and error rate conditions. The performance is calculated at the genome level, such that counts of TPs, FPs, and FNs are summed per genome and then used for calculating the F1 score for each genome. Each point in a boxplot thus corresponds to the performance on one genome in the test set. The results are based on all CDS fragments longer than 60bp. Note that the y-axes are shown in the range [0.35, 1.0].

**Table 1:** Aggregated test set performance across error rate and sequence length conditions. Aggregated CDS-level (F1 score, sensitivity, and precision) and codon-level (MCC) performance on the test sets of different error rates and sequence lengths, based on the 212 test genomes that follow the standard prokaryotic genetic code. Aggregated means that counts of TPs, FNs, FPs and TNs are summed across all 212 genomes, from which the presented scores are calculated. The numbers marked in bold highlight the highest performance metric obtained for the specific error rate and sequence length condition. The results are based on all CDS fragments longer than 60bp. Standard errors are omitted from the cells for readability; across all metrics, read lengths, and error rates, the genome-level cluster bootstrap standard error (see Methods Section 2.5.4) spans, per model: FGS (Complete) 0.001–0.004; FGS (0.5% error rate) 0.001–0.004; FGS (1.0% error rate) 0.001–0.005; MetaProdigal 0.001–0.003; DeepCDS N 0.000–0.003; DeepCDS S 0.000–0.004; DeepCDS S+I 0.000–0.004.

|  | **Typical-error** |  |  |  | **High-error** |  |  |  | **Stress-error** |  |  |  |
|---|---|---|---|---|---|---|---|---|---|---|---|---|
|  | 75bp | 100bp | 150bp | 300bp | 75bp | 100bp | 150bp | 300bp | 75bp | 100bp | 150bp | 300bp |
| **CDS-level F1 Score** |  |  |  |  |  |  |  |  |  |  |  |  |
| FGS (Complete) | 0.803 | 0.828 | 0.842 | 0.810 | 0.790 | 0.809 | 0.812 | 0.751 | 0.745 | 0.751 | 0.723 | 0.591 |
| FGS (0.5% error rate) | 0.796 | 0.815 | 0.818 | 0.765 | 0.782 | 0.797 | 0.791 | 0.714 | 0.738 | 0.740 | 0.707 | 0.571 |
| FGS (1.0% error rate) | 0.791 | 0.809 | 0.809 | 0.752 | 0.778 | 0.790 | 0.782 | 0.702 | 0.733 | 0.734 | 0.699 | 0.563 |
| MetaProdigal | 0.810 | 0.842 | 0.868 | 0.867 | 0.796 | 0.823 | 0.839 | 0.808 | 0.752 | 0.766 | 0.753 | 0.646 |
| DeepCDS N | 0.866 | 0.906 | 0.932 | 0.923 | 0.852 | 0.888 | 0.904 | 0.872 | 0.805 | 0.829 | 0.819 | 0.725 |
| DeepCDS S | **0.873** | **0.913** | **0.942** | **0.944** | **0.863** | **0.903** | **0.931** | **0.930** | **0.827** | **0.867** | **0.892** | 0.881 |
| DeepCDS S+I | 0.854 | 0.898 | 0.935 | 0.939 | 0.844 | 0.888 | 0.924 | 0.927 | 0.809 | 0.851 | 0.885 | **0.884** |
| **CDS-level Sensitivity** |  |  |  |  |  |  |  |  |  |  |  |  |
| FGS (Complete) | 0.855 | 0.873 | 0.879 | 0.849 | 0.838 | 0.852 | 0.850 | 0.797 | 0.786 | 0.788 | 0.759 | 0.648 |
| FGS (0.5% error rate) | 0.848 | 0.863 | 0.856 | 0.791 | 0.832 | 0.843 | 0.828 | 0.744 | 0.781 | 0.779 | 0.743 | 0.611 |
| FGS (1.0% error rate) | 0.844 | 0.856 | 0.846 | 0.775 | 0.828 | 0.836 | 0.819 | 0.729 | 0.776 | 0.774 | 0.735 | 0.598 |
| MetaProdigal | 0.849 | 0.876 | 0.890 | 0.881 | 0.833 | 0.856 | 0.859 | 0.827 | 0.780 | 0.791 | 0.769 | 0.673 |
| DeepCDS N | 0.880 | 0.909 | 0.925 | 0.922 | 0.859 | 0.885 | 0.893 | 0.873 | 0.795 | 0.808 | 0.792 | 0.725 |
| DeepCDS S | 0.886 | 0.904 | 0.921 | 0.933 | 0.876 | 0.904 | 0.923 | **0.927** | 0.827 | 0.856 | 0.872 | 0.874 |
| DeepCDS S+I | **0.890** | **0.918** | **0.937** | **0.942** | **0.898** | **0.913** | **0.924** | 0.924 | **0.853** | **0.867** | **0.875** | **0.876** |
| **CDS-level Precision** |  |  |  |  |  |  |  |  |  |  |  |  |
| FGS (Complete) | 0.758 | 0.787 | 0.807 | 0.774 | 0.746 | 0.771 | 0.778 | 0.710 | 0.708 | 0.718 | 0.691 | 0.543 |
| FGS (0.5% error rate) | 0.750 | 0.773 | 0.784 | 0.741 | 0.738 | 0.756 | 0.756 | 0.686 | 0.700 | 0.705 | 0.674 | 0.536 |
| FGS (1.0% error rate) | 0.745 | 0.766 | 0.775 | 0.730 | 0.733 | 0.749 | 0.748 | 0.677 | 0.695 | 0.698 | 0.667 | 0.532 |
| MetaProdigal | 0.774 | 0.809 | 0.848 | 0.854 | 0.763 | 0.793 | 0.820 | 0.791 | 0.725 | 0.743 | 0.738 | 0.621 |
| DeepCDS N | 0.853 | 0.902 | 0.939 | 0.923 | 0.845 | 0.890 | 0.916 | 0.871 | 0.816 | 0.851 | 0.847 | 0.724 |
| DeepCDS S | **0.857** | **0.908** | **0.948** | **0.945** | **0.850** | **0.902** | **0.940** | **0.932** | **0.828** | **0.880** | **0.913** | 0.888 |
| DeepCDS S+I | 0.804 | 0.872 | 0.933 | 0.940 | 0.796 | 0.864 | 0.924 | 0.930 | 0.770 | 0.836 | 0.894 | **0.892** |
| **Codon-level MCC** |  |  |  |  |  |  |  |  |  |  |  |  |
| FGS (Complete) | 0.790 | 0.840 | 0.888 | 0.933 | 0.775 | 0.823 | 0.871 | 0.917 | 0.726 | 0.771 | 0.817 | 0.866 |
| FGS (0.5% error rate) | 0.787 | 0.834 | 0.881 | 0.926 | 0.772 | 0.818 | 0.865 | 0.914 | 0.723 | 0.768 | 0.815 | 0.875 |
| FGS (1.0% error rate) | 0.784 | 0.831 | 0.877 | 0.922 | 0.769 | 0.816 | 0.863 | 0.912 | 0.721 | 0.766 | 0.814 | 0.874 |
| MetaProdigal | 0.795 | 0.848 | 0.904 | 0.955 | 0.780 | 0.831 | 0.885 | 0.937 | 0.730 | 0.777 | 0.825 | 0.877 |
| DeepCDS N | 0.857 | 0.914 | 0.960 | 0.985 | 0.842 | 0.899 | 0.947 | 0.974 | 0.793 | 0.851 | 0.903 | 0.936 |
| DeepCDS S | **0.866** | **0.920** | **0.965** | **0.988** | **0.854** | **0.911** | **0.958** | **0.986** | **0.816** | **0.878** | **0.934** | 0.973 |
| DeepCDS S+I | 0.846 | 0.906 | 0.960 | 0.987 | 0.835 | 0.897 | 0.954 | 0.985 | 0.798 | 0.865 | 0.930 | **0.974** |

All models were also compared to MetaGeneMark-2 on the 300bp test sets in a separate benchmark restricted to CDSs of at least 90 bp (Section 2.5.3). The DeepCDS models achieved the highest F1 score across all error conditions and IoU thresholds, followed by MetaProdigal (Supplementary Figure A13). MetaGeneMark-2 performed comparably to FGS (Complete) under less strict overlap criteria, but below it with exact coordinate matching. Although MetaGeneMark-2 can automatically detect genetic code 4 [26], it also performed similarly to FGS (Complete), and well below MetaProdigal and DeepCDS, on the three *Mycoplasmataceae* test genomes using genetic code 4 (Supplementary Figure A14). This is likely because MetaGeneMark-2 only considers genetic code 4 for sequences of at least 5,000 bp and therefore applied the standard prokaryotic genetic code to all test sequences.

DeepCDS also substantially improves start and stop codon identification across tested conditions (Figure 4 and Supplementary Figure A15). For example, on the 75bp error-free test set, DeepCDS N increases the median start codon F1 score from 0.152 (MetaProdigal) to 0.438, and on the 300bp stress-error test set, DeepCDS S achieves a start codon F1 score of 0.771 compared to 0.469 for MetaProdigal, which is an improvement indicative of practically useful annotation performance at challenging conditions.

**Fig. 4:**
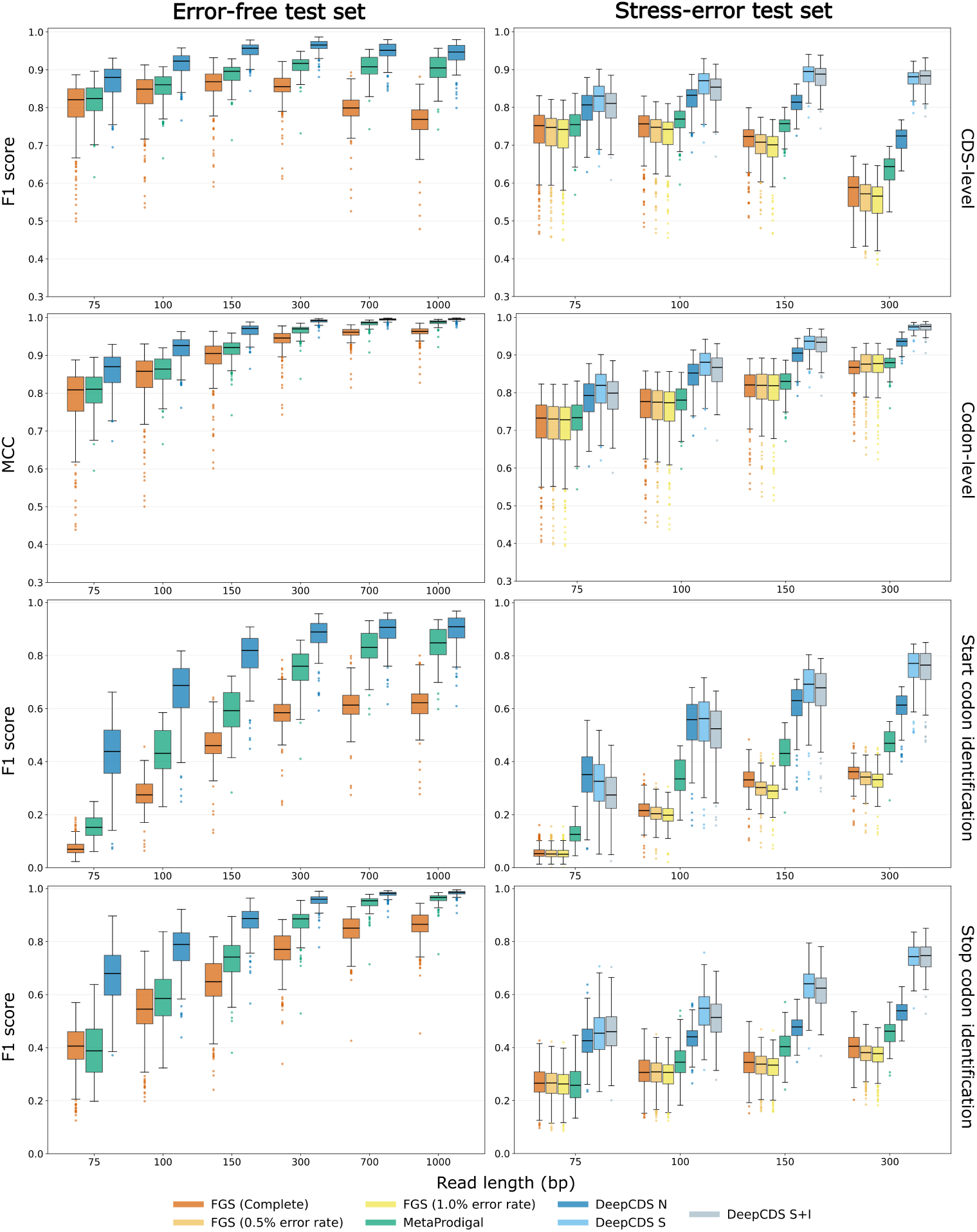
Multi-metric genome-level performance on error-free and stress-error test sets. Boxplots visualizing a range of performance metrics for the error-free and stress-error test sets simulated at different sequence lengths and measured at the genome level for the 212 test genomes that follow the standard prokaryotic genetic code. The results are based on all CDS fragments longer than 60 bps. Each row shows a different performance metric: (1) the CDS-level F1 score. (2) the codon-level MCC, where a codon is a TP if it is both coding and predicted as coding, etc. (3) start codon identification F1 score. (4) stop codon identification F1 score. Note that the y-axes for the CDS-level and codon-level performance are shown in the range [0.3, 1.0]. The corresponding results for the typical-error and high-error test sets are shown in Supplementary Figure A15.

Across all error conditions and models, both the overall CDS detection and the start and stop codon prediction performance improve with sequence length and decline with error rate. While these results provide a brief overview of model performance, the following sections examine the most important findings and contributions of DeepCDS in greater detail.

### 3.2 Robustness to sequencing errors

Sequencing errors disrupt the biologically meaningful signal embedded within original sequences, as demonstrated by the progressive performance degradation across all models as error rates increase (Figure 3 and Table 1). However, the degree to which this noise affects each model’s performance differs notably. The DeepCDS models trained on error-containing data (S and S+I variants) are notably less sensitive to error-prone sequences, which becomes particularly evident on the stress-error test sets and at longer sequence lengths. While the performance of DeepCDS N degrades notably compared to the S and S+I variants, it still outperforms MetaProdigal and the FGS models, suggesting that the deep learning framework employed contributes meaningfully to CDS detection independent of the error-robustness training.

Across the three error rate conditions, DeepCDS S improves the aggregated CDS-level F1 score over MetaProdigal by 0.077–0.235 and over FGS (Complete), which was the best performing FGS variant, by 0.134–0.290 at 300bp, with the largest discrepancies observed under the stress-error condition (Table 1). The trend is consistent at the genome level across all tested conditions, where each DeepCDS variant achieves higher median scores across the distribution of test genomes, indicating robust performance elevation rather than aggregate gains driven by a subset of genomes (Figures 3 and 4, Supplementary Figure A16, and Supplementary Table A28).

### 3.3 Improved detection of start and stop codons

Start and stop codon positions define the exact boundaries of a prokaryotic CDS, making their accurate identification essential for reliable gene annotation. On the error-free test sets, the improvement with DeepCDS N is most substantial at shorter sequence lengths where limited context makes precise boundary detection inherently challenging (Figure 4). Notably, even at the longest sequence length tested (1,000 bp), DeepCDS N achieves the best performance for both start and stop codon prediction, increasing the genome-level median start codon F1 score from 0.848 (MetaProdigal) to 0.909, and the stop codon F1 score from 0.966 (MetaProdigal) to 0.986, despite being trained on shorter sequences of 300bp.

Under increasing error rate conditions, the DeepCDS models trained on error-containing data (S and S+I variants) are most robust in start and stop codon identification, with substantial performance gains at longer sequence lengths compared to the other models (Figure 4 and Supplementary Figure A15). This suggests that training on error-prone sequences has enabled DeepCDS to rely on broader sequence context patterns, learning to tolerate error-induced sequence variation at the boundaries of coding and non-coding regions; for example, recognizing stop codons despite one or more substitution errors in the coding region. Taken together, these findings suggest that DeepCDS is robust at predicting exact start and stop codon positions, rather than merely identifying approximate CDS boundaries.

For the error-free test sets, we assessed performance on canonical (ATG) and groups of non-canonical (GTG, TTG, and others) separately. DeepCDS N achieves the best F1 score, precision, and sensitivity across all test sets and start codon categories, whether start codons at sequence borders are excluded (Table 2) or included (Supplementary Table A26). All three tools identify canonical start codons more accurately than non-canonical ones, likely reflecting the greater frequency of canonical starts in the training data. Performance improves with increasing sequence length across all models. At 75 bp, sensitivity on all non-canonical start codons is very low for all tools, showing that nearly all true start codons are missed at this length. In contrast, DeepCDS N obtains a notably higher precision for GTG and TTG start codons (0.748 and 0.695, respectively) at 75bp, compared to MetaProdigal, which achieves the next best precisions of 0.292 at start GTG codons and 0.044 at TTG start codons. Notably, none of the models are capable of making reliable predictions on other non-canonical start codons than GTG and TTG, with all FGS (Complete) and MetaProdigal scores being near-zero. As sequence length increases, FGS (Complete)’s performance on non-canonical start codons plateaus, peaking at an F1 score of 0.385 for GTG start codons at 700bp, while both MetaProdigal and DeepCDS N continue to improve, reaching F1 scores of 0.729 and 0.819, respectively, at 1,000 bp (Table 2). Including start codons positioned at the sequence border (assessed only for DeepCDS N and MetaProdigal) slightly reduces F1 scores for both models across all sequence lengths, though DeepCDS N retains the better performance (Supplementary Table A26). Notably, MetaProdigal’s precision remains unchanged in this comparison, indicating that it never predicts a start codon at a sequence border. This suggests that, despite making explicit start codon calls throughout, MetaProdigal is not designed to call start codons at sequence edges.

**Table 2:**
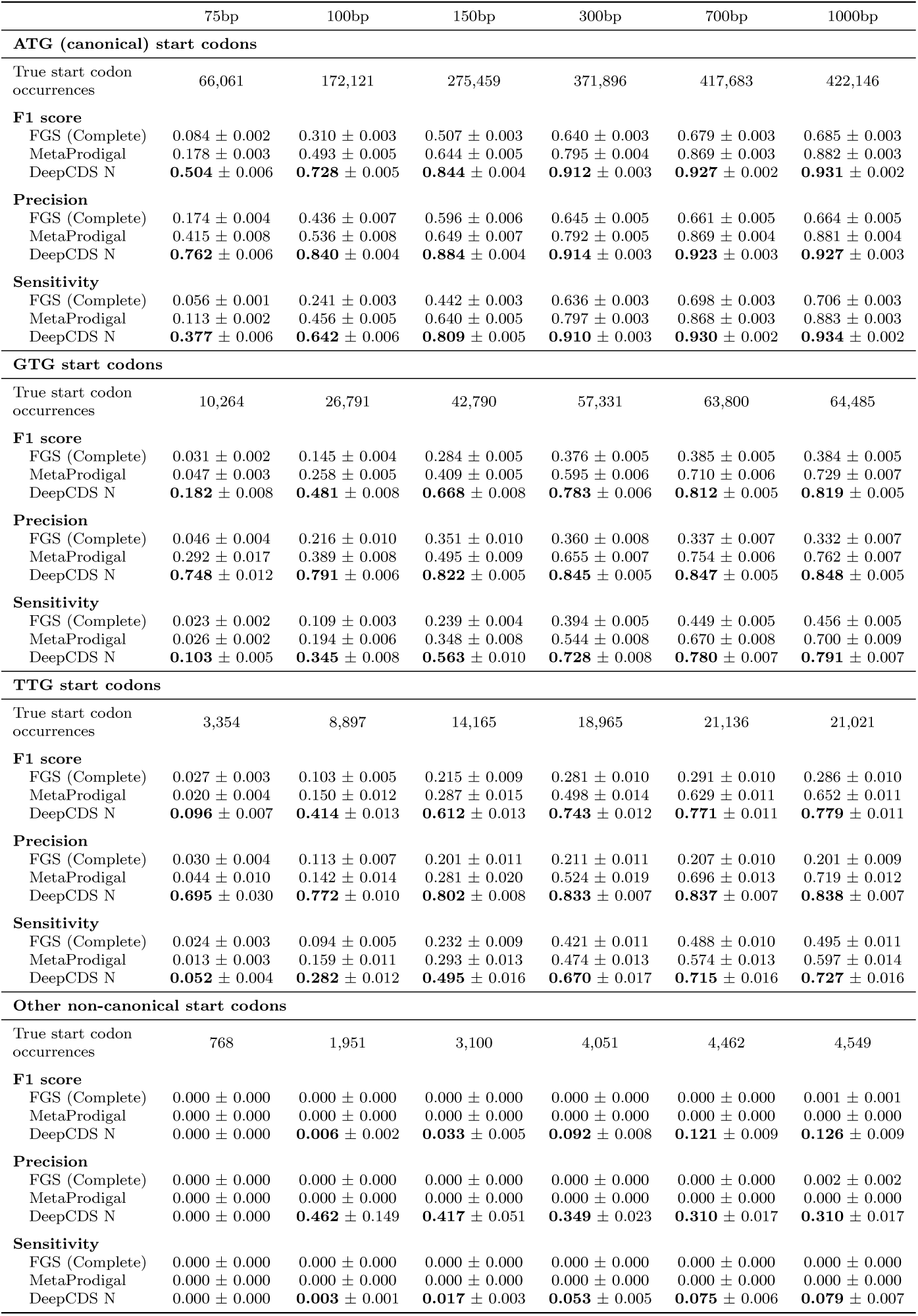
Identification of canonical and non-canonical start codons in error-free test sequences. Start codon identification performance, calculated separately for canonical (ATG) and groups of non-canonical (GTG, TTG, and others) start codons on the error-free test sets simulated at each sequence length. Scores are based on summed TP, FP, and FN counts aggregated across the 212 test genomes using the standard prokaryotic genetic code. The standard error from a genome-level cluster bootstrap is reported (*±*; see Methods Section 2.5.4). The best performance for each test set is shown in bold. Start codons at sequence borders are excluded from both 31 true occurrences and predictions, since FGS does not explicitly annotate start codon positions and border cases cannot be reliably assessed as true starts versus internal CDS regions; for a comparison of MetaProdigal and DeepCDS N including border start codons, see Supplementary Table A26. It should be noted that the number of occurrences of true start codons increases with test set sequence length; this is due to the fact that many CDS parts in the test sets become shorter than 61 bp and for that reason are excluded from the results, as FGS and MetaProdigal do not predict shorter stretches of CDSs.

#### Model behavior on CDSs with conflicting annotated start positions

For training and for the main assessment of start codon identification, the start codon was defined as the first codon of the annotated CDS. Where the RefSeq and GenBank annotations shared a stop codon but differed in their start positions, we retained the longer CDS (Section 2.2). As we cannot know which version reflects the true translation initiation site, we examined whether this choice biased DeepCDS towards the longer version. In the annotations of the 212 test genomes using the standard prokaryotic genetic code, 6.6% of all CDSs had two annotated start positions for the same stop codon. In each 300bp test set, we selected all reads containing both start positions, and where both resulting CDS fragments would be longer than 60 bp within the read. For each of these reads, we recorded which of the two start positions each model predicted. The DeepCDS models predicted neither start position less often than FGS and MetaProdigal in all test sets, and predicted both the longer and the shorter version more often (Supplementary Table A29). All models predicted the longer version more frequently than the shorter one, with ratios between 1.33 and 1.67. The ratios of the DeepCDS models (1.40–1.51) were in the middle of this range. During training, DeepCDS saw only the longer version, with the shorter version’s start position labeled as coding. Nevertheless, its preference for the longer version was comparable to that of MetaProdigal and FGS, which were not trained on these annotations. This suggests that retaining the longer CDS did not cause a systematic bias in DeepCDS.

### 3.4 Accurate coding sequence prediction across phylogenetic diversity and GC-content ranges

Robust performance across diverse phylogenetic groups, genomic GC-content ranges, and genetic codes is essential for CDS predictors targeting metagenomic sequences. We evaluated DeepCDS separately on 16 representative organisms from the test set (Methods Section 2.3 and Supplementary Table A3), and aggregated all 215 test genomes to family-level (Supplementary Table A16), on the 300bp typical-error test set. DeepCDS S achieves the highest CDS-level F1 score both across all 16 representative genomes and their family-level aggregations (Supplementary Table A28 and Supplementary Figure A16), suggesting that DeepCDS generalizes well across phylogenetically diverse prokaryotic organisms.

The assessment of *M. capricolum* and its family *Mycoplasmataceae* (Supplementary Table A28 and Supplementary Figure A16) provides insight into model behavior on organisms employing an alternative genetic code (genetic code 4). On *M. capricolum*, MetaProdigal achieves a CDS-level F1 score of 0.873 while the DeepCDS models range from 0.913 to 0.933, with DeepCDS S yielding the best performance (Supplementary Table A28). These results suggest that DeepCDS can handle sequences from genomes using this alternative genetic code, although all sequences are translated using the standard prokaryotic genetic code for the pLM input (Methods Section 2.4). In contrast, the FGS models exhibit substantially reduced performance across *Mycoplasmataceae* (CDS-level F1 scores *<* 0.45), indicating limited robustness toward non-standard translation tables. As this evaluation is limited to three genomes from a single family, a broader evaluation is needed to establish general robustness to alternative genetic codes.

We calculated the Spearman correlation (*ρ*) between the genomic GC-content and CDS-level F1 score, observing increasing correlation with error rate, most notably for bacterial genomes (Supplementary Figure A17 and Supplementary Table A30). On the stress-error test set, FGS shows the strongest correlation on bacterial genomes (*ρ ∈* [0.79; 0.92]), comparable to MetaProdigal (*ρ* = 0.8). While DeepCDS N shows a high correlation as well (*ρ* = 0.78), the DeepCDS variants trained on error-containing data (S and S+I) exhibit lower correlation (*ρ* = 0.34 and *ρ* = 0.33, respectively). We hypothesize this reflects the AT-rich nature of start and stop codons: in AT-rich genomes, substitution errors are more likely to spuriously introduce or abolish such codons, degrading prediction accuracy. The comparatively weak correlation for DeepCDS S and S+I across all error rate conditions demonstrates that training on noisy data increases the robustness to variations in genomic GC-content, a property particularly relevant for metagenomic applications where sequence input spans a broad range of GC compositions.

### 3.5 Performance evaluation on a simulated metagenomic dataset

The CDS-level evaluation on the “Toy Longitudinal Human Gut” test set was carried out on the full cleaned test set, the cleaned test set excluding uncertain annotations, and the cleaned test set excluding contigs completely lacking annotations, for as complete an assessment as possible (see Section 2.5.1 and Supplementary Table A17). The F1 score, precision, and sensitivity were calculated based on summing TP, FP, and FN counts across the whole dataset. We chose this approach to characterize performance across the metagenomic sample as a whole, rather than being disproportionately shaped by individual genomes that may have incomplete annotations in this dataset. The DeepCDS models achieve the best performance across all three versions of the cleaned test set, considering both F1 score, sensitivity and precision (Table 3), with DeepCDS S reaching the highest F1 score but DeepCDS S+I achieving the highest sensitivity. The genomes from which sequences were simulated in the CAMI III “Toy Longitudinal Human Gut” datasets are, as mentioned, publicly available. However, the degree to which they are correctly and completely annotated is difficult to quantify. Some genomes may be considerably under-annotated, which could, hypothetically speaking, result in higher FP rates. Because we are uncertain about the degree to which under-annotation is a problem, the precision metric and ultimately F1 score should be viewed and interpreted with caution. While we evaluate on a test set where reads from contigs completely lacking annotations are excluded, the remaining contigs might still be under-annotated (Table 3, “Excluding contigs lacking annotations”). For this reason, the more appropriate metric to evaluate performance is the sensitivity, measuring the degree of actual CDSs that are also predicted to be CDSs. We find this assessment relatively trustworthy given our initial test set cleaning and processing, and show the results for the test set where reads overlapping with uncertain annotations, defined by being hypothetical or ab initio predicted, are excluded (Table 3, “Excluding uncertain annotations”).

**Table 3:** CDS-level performance on a CAMI III simulated metagenomic dataset. CDS-level predictive performance based on the cleaned CAMI III “Toy Longitudinal Human Gut” test set in three versions: (i) the base, (ii) excluding reads overlapping with uncertain annotations (defined by being hypothetical or ab initio predicted), and (iii) excluding contigs without any annotations (Section 2.5.1 and Supplementary Table A17). In the base test set, uncertain annotations are treated as being correct. The standard error from a genome-level cluster bootstrap is reported (*±*) (see Methods Section 2.5.4).

|  | F1 Score | Sensitivity | Precision |
|---|---|---|---|
| **Base** |  |  |  |
| FGS (Complete) | 0.766 ± 0.060 | 0.878 ± 0.003 | 0.679 ± 0.087 |
| FGS (0.5% error rate) | 0.748 ± 0.056 | 0.858 ± 0.002 | 0.662 ± 0.081 |
| FGS (1.0% error rate) | 0.740 ± 0.055 | 0.849 ± 0.003 | 0.656 ± 0.080 |
| MetaProdigal | 0.795 ± 0.045 | 0.872 ± 0.011 | 0.731 ± 0.072 |
| DeepCDS N | 0.890 ± 0.033 | 0.924 ± 0.007 | 0.858 ± 0.060 |
| DeepCDS S | **0.899 ± 0.034** | 0.936 ± 0.006 | **0.865 ± 0.060** |
| DeepCDS S+I | 0.887 ± 0.040 | **0.938 ± 0.006** | 0.841 ± 0.069 |
| **Excluding uncertain annotations** |  |  |  |
| FGS (Complete) | 0.760 ± 0.060 | 0.886 ± 0.003 | 0.665 ± 0.086 |
| FGS (0.5% error rate) | 0.744 ± 0.058 | 0.869 ± 0.003 | 0.651 ± 0.082 |
| FGS (1.0% error rate) | 0.737 ± 0.059 | 0.861 ± 0.003 | 0.645 ± 0.084 |
| MetaProdigal | 0.791 ± 0.046 | 0.880 ± 0.012 | 0.719 ± 0.073 |
| DeepCDS N | 0.889 ± 0.035 | 0.931 ± 0.007 | 0.850 ± 0.063 |
| DeepCDS S | **0.899 ± 0.038** | 0.944 ± 0.006 | **0.858 ± 0.067** |
| DeepCDS S+I | 0.886 ± 0.044 | **0.946 ± 0.006** | 0.834 ± 0.073 |
| **Excluding contigs lacking annotations** |  |  |  |
| FGS (Complete) | 0.841 ± 0.010 | 0.878 ± 0.003 | 0.807 ± 0.019 |
| FGS (0.5% error rate) | 0.821 ± 0.012 | 0.858 ± 0.002 | 0.788 ± 0.021 |
| FGS (1.0% error rate) | 0.813 ± 0.012 | 0.849 ± 0.003 | 0.780 ± 0.021 |
| MetaProdigal | 0.857 ± 0.010 | 0.872 ± 0.011 | 0.842 ± 0.014 |
| DeepCDS N | 0.933 ± 0.004 | 0.924 ± 0.007 | 0.943 ± 0.005 |
| DeepCDS S | **0.943 ± 0.004** | 0.936 ± 0.006 | **0.950 ± 0.005** |
| DeepCDS S+I | 0.936 ± 0.004 | **0.938 ± 0.006** | 0.935 ± 0.008 |

**Table 4:** Protein-level support of predicted CDSs on a real metagenome short-read dataset. Predicted CDSs (*>*60bp on either strand) on the 58.4M reads (*>*60bp) of the real gut metagenome sample MH0161 and their support by UniRef90. A CDS has a hit if DIAMOND [62] finds a match with E-value *≤* 10*^−^*^3^; none of 140,672 shuffled control sequences had a hit. Bold marks the most CDSs with a hit, the fewest CDSs without a hit, and the highest share with a hit. As CDSs without a hit may represent genes without known homologs, the share with a hit is a lower bound on the fraction of genuine CDSs.

| Model | Reads with<br>a CDS (%) | CDSs (Million) |  |  | CDSs with<br>a hit (%) |
|---|---|---|---|---|---|
|  |  | Predicted | With hit | Without hit |  |
| FGS (Complete) | 92.5 | 54.04 | 39.76 | 14.28 | 73.6 |
| FGS (0.5% error rate) | 93.3 | 54.51 | 39.60 | 14.91 | 72.6 |
| FGS (1.0% error rate) | 93.5 | 54.62 | 39.46 | 15.16 | 72.2 |
| MetaProdigal | 88.5 | 51.68 | 38.79 | 12.89 | 75.1 |
| DeepCDS N | 80.1 | 47.54 | 39.80 | 7.74 | 83.7 |
| DeepCDS S | 81.1 | 48.10 | 40.39 | **7.71** | **84.0** |
| DeepCDS S+I | 85.4 | 51.58 | **41.53** | 10.05 | 80.5 |

We also observe error bars of greater magnitude for the precision metric in the test set versions where we include reads from contigs lacking annotations, compared to the sensitivity. Furthermore, the magnitude of the error bars decline on the test set where we exclude reads from contigs lacking annotations. This suggests that the assumption regarding under-annotation and varying degrees of FPs, some which are actual FPs, and perhaps others which might be actual CDSs not annotated, could be true.

#### Assessment of runtime

The full metagenomic dataset consisted of 33,267,026 individual 150bp reads (from 16,633,513 read pairs; before removing genomes that share a species with the model development sets). Inference on the full dataset, including predictions made on both the input sequences and their reverse complement sequences for analysis completeness, took on average 29 minutes for a single FGS model (re-implemented version [12]) on CPU (19,119 reads/second), 8 hours and 20 minutes for a single DeepCDS model on an L40S GPU (1,109 reads/second), and 3 hours and 11 minutes for MetaProdigal on CPU (2,903 reads/second). Throughput and peak memory usage for both GPU and CPU runs across sequence lengths and inference batch sizes are reported in Supplementary Table A31.

### 3.6 DeepCDS predictions on a real gut metagenome are more frequently supported by known proteins

To assess DeepCDS on real sequencing data, we applied all models to the human gut metagenome sample MH0161, using the count of predicted CDSs with a UniRef90 hit as a proxy for their support (see Sections 2.5.1, “Real Illumina-sequenced gut metagenome from the Danish MetaHIT cohort” and 2.5.4, “Protein-level support of predicted CDSs from a real gut metagenome dataset”). As a CDS without a hit may still belong to a genuine gene without known homologs, the share of supported CDSs is a lower bound for each tool, and most informative for comparing tools with each other. The predicted CDSs had nearly identical length distributions across tools (Supplementary Figure A9), so differences in support cannot be attributed to differences in prediction length.

While MetaProdigal and FGS predicted CDSs on more reads than DeepCDS (88.5– 93.5% vs. 80.1–85.4%), both exhibited fewer absolute counts and lower share of their predicted CDSs with a UniRef90 hit (38.79M–39.76M vs. 39.80M–41.53M; 72.2–75.1% vs. 80.5–84.0%, see Table 4). Of all models, DeepCDS S+I recovered the most CDSs supported by UniRef90 (41.53M), whereas DeepCDS S predicted the fewest CDSs without this support (7.71M).

We grouped the reads by which models predicted a CDS in them (Supplementary Table A32). On the 74.9% of reads where all models predicted a CDS, 84.7–85.7% of the DeepCDS predictions had a UniRef90 hit, compared to 82.1% for MetaProdigal and 82.9% for FGS (Complete), indicating fewer unsupported predictions on the same sequences. In all 21 groups where at least one DeepCDS model and MetaProdigal or FGS (Complete) predicted a CDS, a DeepCDS model had the highest share of supported CDSs.

Predictions made by a single tool were rarely supported, but more often for Deep-CDS: 20.0–32.3% of the predictions made by a single DeepCDS model had a hit, compared to 11.9% for MetaProdigal and 13.3% for FGS (Complete), and the Deep-CDS models made such predictions on far fewer reads (0.03–0.33% vs. 0.91% and 3.15%). Similarly, on the 0.29% of reads where only the three DeepCDS models predicted a CDS, 59.2–60.6% of their predictions had a hit, whereas on the 4.3% of reads where only MetaProdigal and FGS (Complete) predicted a CDS, 23.6–23.7% of their predictions had a hit. Thus, while DeepCDS misses some genuine CDSs on the latter reads, the majority of the additional predictions by MetaProdigal and FGS (Complete) lacked support from known proteins.

### 3.7 Generalization beyond homology, similarity, and well-characterized coding sequences

DeepCDS was trained on genomes partitioned by family-level taxonomy, using only labels from well-characterized CDS annotations (i.e., excluding hypothetical proteins and explicitly tagged ab initio predictions). To assess whether its performance relied on similarity between training and test sequences, we stratified the 300bp error-free test set by sequence identity to the training and validation sequences at both the protein level (translated CDS fragments) and the nucleotide level (complete sequences) (Figure 5).

**Fig. 5:**
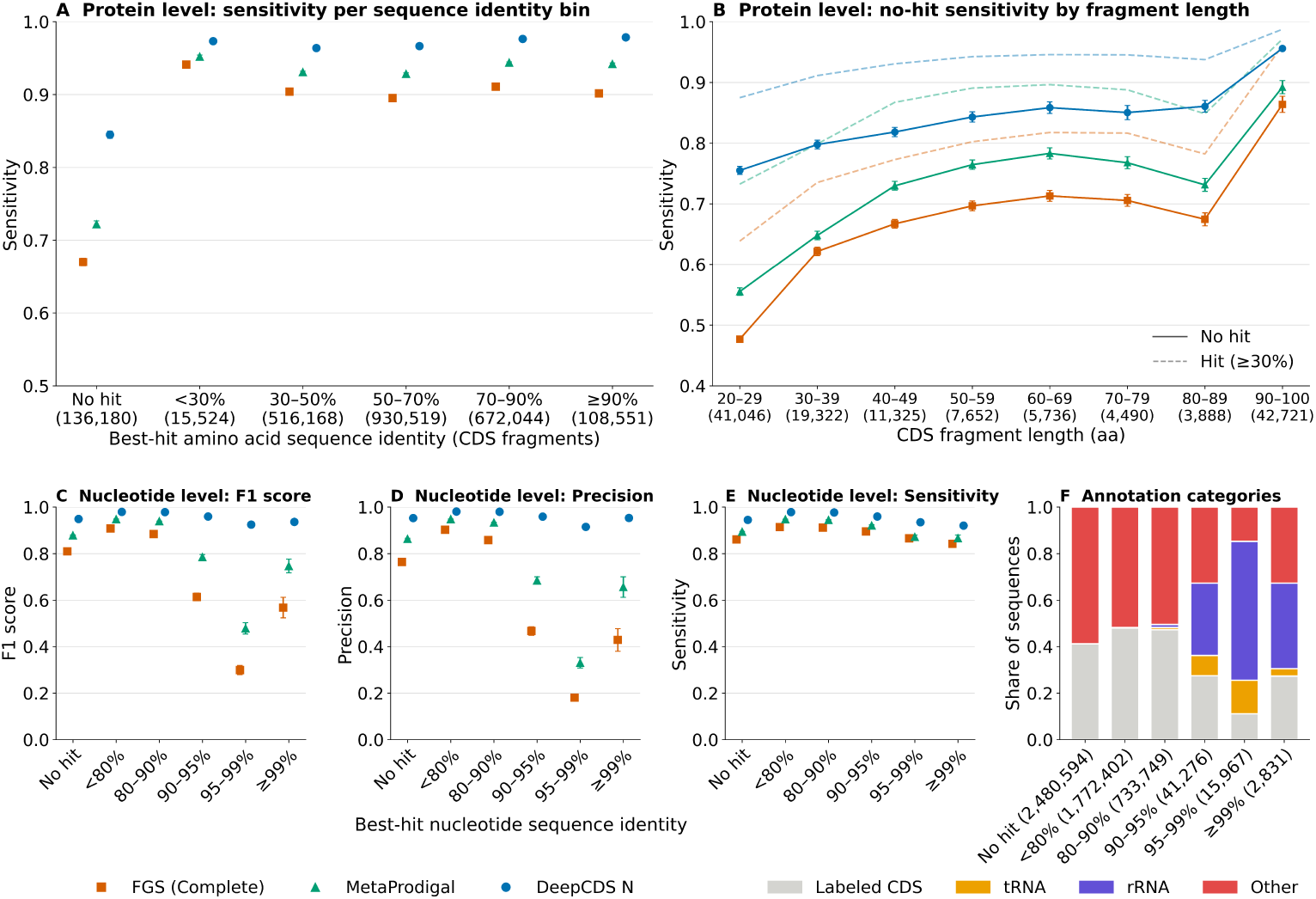
Performance stratified by similarity to the training data. Test set performance stratified by the sequence identity of each test sequence to the training and validation sequences, based on the best hit by bitscore using MMseqs2 [57] (*E ≤* 10). The results are based on the 300 bp error-free test set of the 212 genomes using the standard prokaryotic genetic code. “No hit” corresponds to no detected alignment of at least 20 amino acids (protein level) or 60 nucleotides (nucleotide level). Performance is CDS-level with exact matching of predicted and labeled CDSs. **(A)** Protein-level sensitivity of labeled CDS fragments (*>*60 bp), binned by the amino acid sequence identity of the best hit among the training and validation CDS fragments. **(B)** Sensitivity of “No hit” fragments by fragment length (solid lines). Fragments with a hit of *≥*30% identity are shown for reference (dashed lines). **(C–E)** Nucleotide-level F1 score, precision, and sensitivity, with test sequences binned by the nucleotide sequence identity of their best hit among the training and validation sequences; all test sequences are included, also those without a labeled CDS. **(F)** Share of test sequences per nucleotide identity bin that contain a labeled CDS fragment, or otherwise lie within an annotated rRNA or tRNA gene (overlap *>*60 bp, either strand). Sequences were simulated from both strands and are labeled only with CDSs on their own strand; “Other” therefore also comprises sequences antisense to a CDS. Numbers in parentheses give the number of CDS fragments (A, B) or sequences (F; the same counts apply to C–E) per bin. Error bars mark one standard error from a genome-level cluster bootstrap (1,000 replicates). Results for *E ≤* 10*^−^*^3^ are shown in Supplementary Figure A10.

At the protein level, 5.7% of the CDS fragments had no hit. The sensitivity of DeepCDS N was 0.96–0.98 in all identity bins with a hit, without any monotonic or notable increase towards high identity (Figure 5A). Sensitivity was lower on CDS fragments without any hit (0.85), but the drop was larger for MetaProdigal (0.72) and FGS (Complete) (0.67). As neither MetaProdigal nor FGS was trained on our training data, this could suggest that these fragments are inherently harder to recover, rather than the drop in sensitivity for DeepCDS being caused by memorization. Had DeepCDS relied on similarity to its training data, we would instead expect it to be more affected than the other tools. Fragments without a hit were shorter than fragments with a hit (median 46 vs. 99 amino acids), and sensitivity increased with fragment length for all tools. The difference between fragments with and without a hit persisted within each length range, although it decreased with fragment length, and DeepCDS achieved the highest sensitivity at all lengths (Figure 5B).

At the nucleotide level, 49% of the test sequences had no hit and only 1.2% had a hit with *≥* 90% identity, as nucleotide-level search mainly detects close similarity rather than distant homology. The F1 score of DeepCDS was 0.95 on sequences without a hit and 0.93–0.96 on sequences with a *≥* 90% identical hit, and thus did not increase monotonically with similarity to the training data (Figure 5C). For both MetaProdigal and FGS (Complete), precision dropped markedly at *≥* 90% identity, all the way down to 0.33 and 0.18, respectively, at 95–99% identity (Figure 5D). These sequences were enriched for annotated rRNA and tRNA genes (Figure 5F), which are highly conserved across prokaryotes [63, 64]. In the sequences with a *≥* 90% identity hit, 81.4% and 81.9% of the FP CDSs predicted by MetaProdigal and FGS (Complete) overlapped an rRNA or tRNA gene (Supplementary Table A33). DeepCDS predicted far fewer FPs on these sequences (722 vs. 9,186 and 21,447), and its precision remained *≥* 0.91, indicating that it has learned to distinguish conserved non-coding RNA genes from CDSs.

We also assessed performance on hypothetical proteins from the CAMI III sample (Methods Section 2.5.1), representing genes that are likely genuine but not yet functionally characterized. While all models recovered a larger fraction of CDS fragments from functionally annotated genes than from hypothetical proteins, the DeepCDS models achieved the highest sensitivity on hypothetical proteins across all tested IoU thresholds (Supplementary Figure A8). At exact coordinate matching, DeepCDS S+I recovered 90.0% of the CDS fragments, compared to 82.2% for MetaProdigal and 82.9% for FGS (Complete). As some hypothetical protein annotations may themselves be incorrect, lower sensitivity does not necessarily indicate missed CDSs. However, the relative sensitivity across models indicates their ability to recover CDSs in less-characterized genomic regions, supporting the applicability of DeepCDS to such sequences.

### 3.8 Conservative indel error predictions with higher positional accuracy

Because DeepCDS S+I is explicitly trained to predict indel positions, we assessed its ability to detect indel errors using the evaluation setup described in Methods Section 2.5.4 (“Indel detection and localization”). We compared it to the benchmark models also trained to explicitly predict indels, i.e., FGS (0.5% error rate) and FGS (1.0% error rate). DeepCDS S+I was trained on sequences simulated with 0.5% substitution error rate and 0.05% indel error rate (Methods Section 2.2).

DeepCDS S+I is more conservative than the FGS models in calling indel errors, offering a more balanced trade-off between precision and sensitivity on the simulated test sets (Supplementary Figures A11 and A12). The FGS models predict up to 94 times more indels than are present (41–94-fold under typical-error conditions and 6– 18-fold under stress-error conditions), resulting in near-zero precision across all error conditions and sequence lengths. This suggests a systematic overestimation of indel frequency in sequences with simulated modern Illumina-like sequencing error profiles.

In contrast, DeepCDS S+I achieves substantially higher precision and F1 scores, with the performance gap increasing with sequence length. This comes at the cost of lower sensitivity than FGS on the shorter sequences (75–150 bp), whereas sensitivity is comparable at 300 bp.

For all models, the F1 score is near-zero when enforcing an exact match between the true indel error site and the predicted indel error site (Supplementary Figure A12). For the FGS models, precision remains low across all tested positional tolerances, error rates, and sequence lengths, whereas sensitivity increases with positional tolerance and sequence length. For DeepCDS S+I, both precision and sensitivity increases with positional tolerance and sequence length across all error conditions. These results suggest that while the exact indel position is challenging to recover, DeepCDS S+I can, to a limited extent, recover approximate indel positions.

The MAE offers further insight into positional accuracy: when DeepCDS S+I predicts an indel in a read containing a true indel, it localizes the indel with high accuracy, with more than 85% of genomes exhibiting an MAE below 10 bp across all conditions (Supplementary Figure A11). The FGS models produce only slightly higher MAEs on short sequences (75–100 bp; by 1–2.4 bp in genome-level median MAE), but their MAEs increase with sequence length and error rate, with the genome-level median MAE exceeding that of DeepCDS S+I by a factor of approximately two on 300 bp sequences under stress-error conditions. Note that the large number of indels predicted by FGS likely favors its MAE, as more predictions increase the chance of one being located close to a true indel by chance. DeepCDS S+I has a higher number of genomes without any matched indel pairs than the FGS models (*NA in the MAE plots of Supplementary Figure A11), especially at short sequence lengths (75–100 bp) under typical-error conditions. The more conservative indel calling of DeepCDS S+I, combined with the inherently low indel rates of typical Illumina sequencing, makes indel-containing reads in some genomes too sparse for DeepCDS S+I to reliably detect them.

### 3.9 Coding sequence prediction from ultra-short fragments beyond existing tool limits

A key advantage of DeepCDS is its ability to predict CDS fragments shorter than the minimum lengths supported by MetaProdigal and FGS (hardcoded to CDS *≥* 60bp (MetaProdigal) [13] and CDS *≥* 61bp (FGS) [6] respectively). We tested this ability using the same test sets as above but retaining all CDS fragments longer than 30bp, and included test sets simulated at 60bp sequence length (Methods Section 2.5.1). While performance degrades as sequences become shorter, a meaningful predictive signal is retained: on the 60bp error-free test set, DeepCDS N achieves a CDS-level F1 score of 0.815, rising to 0.955 at 300bp, and DeepCDS S achieves 0.765 on the 60bp stress-error test set rising to 0.871 at 300bp (Supplementary Figure A18 and Supplementary Tables A34-A37). These results demonstrate that DeepCDS can predict CDS fragments well below the minimum lengths supported by existing tools, extending its applicability to highly fragmented or low-quality metagenomic data.

### 3.10 Effect of sequence simulator

DeepCDS is trained on sequences simulated with Mason [35] (Methods Section 2.2). To assess the impact of simulator-specific biases on the performance of DeepCDS, we tested its robustness on test sets simulated with an alternative read simulator, art modern [46], using three built-in quality profiles (see Methods Section 2.5.1). The rankings across all four measured metrics are consistent with the Mason results across each of the three tested quality profiles (Supplementary Figure A19). This suggests that DeepCDS has not learned simulator-specific artifacts from its Mason-based training data to a degree that meaningfully affects performance, indicating robustness across sequencing platforms and simulation frameworks.

### 3.11 Nucleotide- and amino acid-level sequence context provide complementary information

To assess the contribution of each input feature, we trained ablation models using only the nucleotide-level codon encoding (**DeepCDS (Codon)**) or the pLM embeddings (**DeepCDS (pLM)**) as input (see Methods Section 2.4.3). Integrating both input types in the full DeepCDS model results in increased CDS-level performance compared to either ablation alone, most notably on shorter sequences, with both ablations approaching that of the full model at longer sequence lengths; a trend observed across all error conditions (Supplementary Figure A20; adjusted p-value *<* 0.05 for all pairwise comparisons between the full model and each ablation across all tested conditions, see Supplementary Note A1.4). On the test sets of shorter sequences, the (Codon) ablation achieves a slightly higher CDS-level and stop codon identification performance than the (pLM) ablation. In contrast, the (pLM) ablations achieve markedly better start codon identification performance compared to the (Codon) ablation, with the difference being most pronounced at shorter sequence lengths (Supplementary Figure A21). Notably, none of the DeepCDS N variants are robust to high error rates, especially at longer sequence lengths.

## 4 Discussion

DeepCDS improves prokaryotic *ab initio* CDS prediction from short, error-prone reads by integrating pLM embeddings with nucleotide-level codon encodings in an end- to-end framework. The results demonstrate robust performance across a wide range of sequencing error rates and sequence lengths, accurate start and stop codon identification, generalization across diverse genomic GC-content ranges, and promising performance on three genomes using an alternative genetic code, all without requiring any prior phylogenetic information. The performance of DeepCDS did not depend on similarity to its training data, and on a real gut metagenome, its predictions were more frequently supported by known proteins than those of the benchmarked tools.

The three model versions (i.e., DeepCDS N, DeepCDS S, and DeepCDS S+I) were trained on datasets with different levels of sequencing error noise, and their hyperparameters were optimized independently. As expected, the optimal settings differed between them (Supplementary Table A13). Forcing all three models to share one set of hyperparameters would therefore introduce its own bias rather than remove one. Comparing each model at its best configuration across the different test sets instead allows us to quantify how well each model performs in the setting it was designed for. We recommend DeepCDS S as the default model for modern short-read sequencing data, as it demonstrated the overall strongest performance across tested conditions with simulated Illumina-like sequencing errors. For error-free or very low error rate sequences, DeepCDS N is the more suitable choice. On the real gut metagenome, DeepCDS S+I recovered the largest absolute number of CDSs supported by known proteins, at the cost of a slightly lower share of supported predictions, and may therefore be preferred when maximizing the number of recovered CDSs is prioritized over precision (Table 4). DeepCDS S+I additionally provides explicit indel error position predictions at a generally modest cost to the overall CDS-level performance and is therefore recommended for highly noisy data where explicit knowledge of frameshifts caused by indel errors is beneficial. DeepCDS S+I was trained exclusively on indels arising as sequencing errors, and its behavior on genuine biological frameshifts has not been evaluated. Predicted indels should therefore be interpreted as putative sequencing errors. Extending DeepCDS to explicitly identify biologically genuine frameshifts is beyond the scope of this work, but represents a potential direction for future development.

Among the benchmark models, MetaProdigal performs surprisingly well on error-containing test sets despite explicitly not accounting for sequencing errors in its development [13]. Contrary to our expectations, the FGS variants developed specifically for error-prone sequences perform worse at the CDS level than both MetaProdigal and FGS (Complete), even on test sets with high error rates. This likely reflects a mismatch between the error profiles used during FGS development and the characteristics of modern Illumina sequencing platforms, particularly in the substitution-to-indel error ratio. The near-zero indel precision of both FGS error models, combined with FGS (1.0% error rate) achieving slightly higher sensitivity than FGS (0.5% error rate), supports this interpretation, and is consistent with higher error rate versions being optimized to predict a progressively larger fraction of positions as indels, including indels in many indel-free reads (Supplementary Figures A11 and A12, “Precision”). This is further consistent with an earlier benchmark of short read CDS predictors [19], which found FGS to be the tool detecting the most prokaryotic CDSs, but at the cost of substantial overprediction of spurious CDSs in non-coding regions. Taken together, these findings support the usage of DeepCDS S+I for indel detection under most practical conditions: the higher precision, better overall F1 score, and more accurate positional localization (lower median genome-level MAE) make its indel calls both more trustworthy and, when identified, more precisely placed in Illumina-like sequences, compared to those of the FGS models. Nonetheless, exact indel localization remains limited for all models (Section 3.8), leaving room for improvement.

On the real gut metagenome, both MetaProdigal and FGS predicted more CDSs without a hit to UniRef90 than the DeepCDS models did (Table 4). Several lines of evidence suggest that a large fraction of these are FPs. On simulated data with known ground truth labels, both tools exhibited both lower precision and sensitivity than DeepCDS across error rates and sequence lengths (Table 1). The test sequences most similar to the training data were enriched for rRNA and tRNA genes, which are highly conserved across taxa [63, 64]. Of the test sequences with *≥*90% nucleotide identity to the training and validation sequences (*E ≤* 10), 49% overlapped an rRNA or tRNA gene by more than 60 bp (Figure 5F). On these sequences, MetaProdigal and FGS predicted 7,478 and 17,567 FP CDSs, respectively (81.4% and 81.9% of their FPs at *≥*90% identity), mostly on sequences overlapping rRNA genes (Supplementary Table A33), whereas DeepCDS predicted only 240 (33.2% of its FPs) (Figure 5C–F). Neither MetaProdigal nor FGS is designed to recognize non-coding RNA genes, whereas DeepCDS was trained with these regions labeled explicitly as non-coding and appears to have learned to distinguish them from protein-coding sequences. The integration of pLM embeddings in the DeepCDS framework may contribute to this, which could be explored in future work.

In general, MetaProdigal and the FGS models predicted CDSs on more reads than DeepCDS, but a smaller share of their CDSs were supported by a hit to a known protein in UniRef90 (72.2–75.1% vs. 84.0% for DeepCDS S), and they predicted 5.2–7.5 million more CDSs without such support (Table 4). This difference was most pronounced on the 4.27% of reads where only MetaProdigal and FGS predicted a CDS: here, 23.6–23.7% of their predictions had a hit, compared to 59.2–60.6% on reads where only the DeepCDS models predicted a CDS, and 82.1–82.9% on reads where all tools predicted a CDS (only FGS (Complete) is included here for clarity; see Supplementary Table A32). These reads thus contain genuine CDSs that DeepCDS misses, alongside a large number of predictions without support from known proteins, which may include genes without known homologs. As rRNA and tRNA genes make up only a small fraction of prokaryotic genomes (median 0.56% of genome sequence positions annotated as either across the 1,125 genomes in the full dataset, see Supplementary Figure A22), misclassified RNA genes can account for only a fraction of these predictions, although MetaProdigal and FGS (Complete) frequently predicted CDSs in sequences overlapping annotated rRNA genes in our simulated test sets (Section 3.7). Across all reads, however, each DeepCDS model recovered more supported CDSs in total than MetaProdigal and the FGS models (Table 4). The higher share of supported predictions by DeepCDS therefore does not result from simply predicting fewer CDSs. Furthermore, DeepCDS achieved the highest sensitivity on CDS fragments without detectable similarity to its training data and on hypothetical proteins, as discussed below, suggesting that its fewer predictions do not primarily reflect novel CDSs that it fails to detect.

While the family-level taxonomy-based whole-genome partitioning cannot completely rule out sequence-level overlap between partitions stemming from conserved protein families, we consider it the most appropriate approach for the intended use case of DeepCDS. It reflects a realistic application scenario, in which individual sequences may be homologous to sequences seen during training, although the organism of origin is unknown to the model. Partitioning by, e.g., gene family instead could bias the model towards specific gene families at the expense of others, possibly reducing robustness and overall usability.

This choice does, however, require that performance does not rely on overlap with the training data, which our homology-stratified analysis supports (Section 3.7): the performance of DeepCDS did not increase monotonically with similarity to its training sequences, and its lower sensitivity on CDS fragments without any hit was also observed, to a larger extent, for MetaProdigal and FGS. The finding that all tools perform worse on these fragments suggests that they are inherently harder to recover. This is partly explained by CDS fragment length, as the fragments without a hit were considerably shorter (median 46 vs 99 amino acids), and sensitivity increased with fragment length for all tools. A further possible explanation is that some of these CDS fragments may derive from taxonomically restricted genes without close homologs in other taxa, which may deviate from the sequence characteristics of typical genes. Note that “no hit” is defined relative to the sequences DeepCDS was trained on rather than to the complete training genomes. A CDS fragment without a hit may thus still belong to a protein with a homolog in a training genome if the corresponding region is not covered by any training sequence.

Beyond the demonstration that the performance of DeepCDS does not depend on similarity to its training data, these results suggest that DeepCDS can generalize to novel and less-characterized CDSs. On CDS fragments without detectable similarity to its training data, DeepCDS N achieved a higher sensitivity than MetaProdigal and FGS (Complete) (0.85 vs. 0.72 and 0.67), and its advantage over these tools was larger than on fragments with similarity to the training data (Figure 5A). Additionally, the DeepCDS models recovered more CDS fragments from hypothetical proteins in the CAMI III sample than MetaProdigal and FGS (Supplementary Figure A8). Together, these results indicate that DeepCDS generalizes to CDSs that are distant from its training data and functionally uncharacterized, supporting its potential for uncovering CDSs from previously uncharacterized microbial diversity.

ESM-2 was pretrained on large protein sequence databases that likely include proteins homologous to those in our test sets. This pretraining is unsupervised and involves no information on CDS boundaries or RFs, which DeepCDS learns exclusively from its own training data. As our analysis measures similarity to that training data only, it cannot rule out a contribution of the general protein knowledge acquired during pretraining; this is, however, inherent to any method building on a pretrained pLM.

The ablation experiments suggest that the nucleotide-level codon encoding and pLM embeddings contribute complementary information, with the full DeepCDS model outperforming both ablations most notably at shorter sequence lengths (Supplementary Figure A20). At longer sequence lengths, both ablations approach the performance of the full model, consistent with each input alone providing sufficient context to assess coding potential when more of the sequence is available. The stronger start codon identification performance of the (pLM) ablation may reflect the pLM’s pretraining on millions of protein sequences, encoding rich information about protein-like features, including a strong prior for what the beginning of a CDS should look like (Supplementary Figure A21). However, as the ablation experiments do not separate the effect of pretraining from that of model architecture and parameter capacity, this remains a hypothesis and could be explored in future work.

The unconventional usage of a pLM for CDS prediction may work well for several reasons. Translated non-coding sequence inputs will likely produce representations that are unfamiliar to the pLM, providing a strong discriminative signal in itself to be leveraged in the downstream model layers. This is consistent with previous work demonstrating that pLMs can capture abrupt shifts in amino acid composition at non-coding/coding boundaries [31]. The approach may furthermore work well for noisy data, because translation introduces redundancy through the genetic code, causing many nucleotide substitutions to map to identical amino acid sequences and thereby reducing the impact of sequencing errors on the pLM input representation. Cross-species generalization may furthermore benefit from the amino acid-level representations, because they are invariant to the choice of synonymous codons, and thus to codon usage bias, which is largely species-specific [41]. The pLM embeddings may therefore provide a signal of coding potential that is less dependent on the organism of origin, complementing the codon-level encoding, which captures such species-specific codon usage.

A limitation to DeepCDS is that it has been developed on simulated sequences derived from whole genomes with high-quality reference annotations (Methods Section 2.2, “Short read processing and labeling”), rather than from real metagenomic datasets. As part of this, sequences overlapping CDSs annotated as hypothetical proteins were excluded from model development to maximize label reliability, which may bias the training data towards well-characterized genes. However, our benchmark covers a broad range of scenarios relevant for metagenomic applications, including test sequences originating from 215 phylogenetically diverse prokaryotic genomes, ranges in genomic GC-content, test sequences generated with an alternative read simulator, CDS fragments without detectable similarity to the training data, and CDS fragments from functionally uncharacterized genes (hypothetical proteins), providing a comprehensive evaluation of model generalizability. Future work could explore whether including hypothetical protein annotations in the training data improves performance on poorly characterized sequences further.

Additionally, we used the metagenomic dataset “Toy Longitudinal Human Gut” from CAMI III for a separate validation. Because this dataset has been simulated based on public microbial genomes, it lacks some of the characteristics of a true metagenomic dataset, including chimeric reads, cross-contamination, adapter carryover, and PCR duplicates. While we acknowledge this shortcoming, we deem it the most appropriate way to approximate the performance on a metagenomic short-read dataset while still being able to define ground truths to compare against. The genomes in the sample may, however, lack complete annotations, which could inflate FP rates. Consistent with this, precision showed substantially larger bootstrap error bars than sensitivity when including reads from contigs lacking annotations (Table 3). We therefore consider sensitivity the more reliable metric in this case.

To complement these evaluations, we applied DeepCDS to a real human gut metagenome. This analysis is, however, subject to some limitations. Similarity to a known protein indicates that a predicted CDS is likely genuine, but does not show that its boundaries are exactly correct, and the absence of a hit does not prove that a prediction is false, as genuinely novel proteins may lack homologs in UniRef90. Homology-based support is therefore a conservative proxy that favors well-characterized proteins, and the share of supported predictions is a lower bound on the fraction of genuine CDSs among them. Furthermore, the analysis is based on a single human gut sample, and performance may differ in other environments, such as soil or marine samples, which contain a larger share of uncharacterized organisms. This analysis was intended as a proof of concept of DeepCDS on real sequencing data, and a systematic evaluation across multiple real metagenomes and environments remains an important direction for future work.

As a deep learning model, DeepCDS is more computationally demanding to run than MetaProdigal and FGS, which rely on more lightweight statistical frameworks. For this reason, we opted for the smallest ESM-2 variant (8M parameters), and future work could explore model distillation to reduce inference time without substantially affecting predictive performance. As is the case for most deep learning-based models, we recommend running DeepCDS on GPU. With a batch size of 128, DeepCDS required at most 2.5 GB of GPU memory across all tested sequence lengths (75–1,000 bp; Supplementary Table A31), at close to maximal throughput, and can thus be run on standard GPUs. While DeepCDS can be run on CPU, inference is substantially slower: on an AMD EPYC 9684X CPU (32 threads), DeepCDS processed between 9 and 142 reads per second depending on sequence length (1,000 and 75 bp, respectively), compared to up to 204 and 3,158 reads per second on an NVIDIA H100 GPU (Supplementary Table A31). This makes CPU-based inference impractical for large-scale metagenomic datasets, although feasible for smaller datasets (approximately 3.5 hours per million 150 bp reads on CPU). As each read is processed independently, CPU-based inference can, however, be parallelized across multiple nodes, and the DeepCDS webserver provides an alternative for running small datasets for users without local computational resources. The computational cost is therefore mainly a consideration for very large datasets where GPU resources are not available.

Looking ahead, the results obtained with DeepCDS open several important future lines of work. The finding that DeepCDS N generalizes well beyond its 300 bp training window using a sliding window approach (tested on up to 1,000 bp error-free sequences, see Figure 4, Supplementary Table A27, and Supplementary Note A1.1) encourages future work testing DeepCDS’s capabilities as a whole-genome prokaryotic annotation tool, either in its current form or optimized for longer context windows, potentially complemented by optional information on taxonomic origin where available. A compelling natural application is ancient DNA, where damage such as post-mortem deamination and high fragmentation rates severely limit available sequence context [65]. These are conditions where the ablation results suggest multi-level sequence representations contribute most complementary sequence information. More broadly, the DeepCDS framework could be extended toward both eukaryotic and viral CDS prediction, where fragmented sequences with, e.g., intron-containing genes or ribosomal slippage mechanisms present similar challenges.

## 5 Conclusion

DeepCDS represents a substantial enhancement in coding sequence prediction from short, error-prone, and fragmented sequences compared to well-established methods in the field, by integrating protein-level context with nucleotide-level information in an end-to-end framework. The ability of DeepCDS to accurately identify coding regions from sequences with limited information highlights the value of combining multiple levels of sequence representation in ab initio gene finding, especially in short sequences where the multiple levels provide the most complementary information. These findings open avenues for extending this approach to more challenging domains, such as ancient DNA and eukaryotic gene prediction.

## Supplementary information

## Supporting information

Supplementary Large Tables

## Acknowledgements

We would also like to acknowledge Peter Wad Sackett for his help with setting up the DeepCDS 1.0 webserver.

## Author contribution

H.N., L.S.N., and O.W. designed the research project. L.S.N. collected and processed the data, developed the model, and carried out the performance evaluation. L.S.N. drafted the manuscript with support from H.N. and O.W. H.N. and O.W. supervised the project. All authors read and approved the final manuscript.

## Data Availability

The datasets supporting the conclusions of this article are available at the DeepCDS 1.0 Webserver site, accessible at: https://services.healthtech.dtu.dk/services/DeepCDS-1.0/ (See Section “Data”).

The raw datasets, including reference genomes in fasta format (accession genomic.fna), and RefSeq and GenBank annotations (*.gff) were collected from NCBI’s Genome database: https://www.ncbi.nlm.nih.gov/datasets/genome/?taxon=2&reference_only=true&annotated_only=true&refseq_annotation=true. Supplementary Table A1 lists the metadata for all genomes in the dataset, as retrieved from NCBI Genome Database.

The DeepCDS 1.0 online server is available at https://services.healthtech.dtu.dk/services/DeepCDS-1.0/. For large datasets, the program can be downloaded locally from the GitHub repository at https://github.com/lsandvad/deep-cds.

The CAMI III “Toy Longitudinal Human Gut” dataset was accessed at: https://cami-challenge.org/datasets/toy-human-gut/.

The Illumina HiSeq 2*×*90bp dataset from the Danish cohort of the MetaHIT project (PRJEB4336, sample MH0161, run ERR321493) was accessed at: https://www.ebi.ac.uk/ena/browser/view/PRJEB4336

## Code availability

DeepCDS, along with trained model weights for each variant (N, S, and S+I), is available in the GitHub repository https://github.com/lsandvad/ deep-cds. The source code used to reach the conclusions of this project is also available in the repository in /src dev.

## Supplementary Materials

Available in file deepcds supplementary.pdf.

