## Supplementary Material for "DeepCDS: *Ab initio* coding sequence prediction in prokaryotic metagenomic short reads"

### A1 Supplementary Notes

#### A1.1 Inference on longer sequences

While the DeepCDS models were trained on sequences of 300 bp, the model can also be applied to longer sequences using a sliding window approach. Sequences exceeding 300 bp are split into overlapping windows of 300 bp (100 codons), matching the training length. Consecutive windows are shifted by a user-defined stride (210 bp in the benchmarks, resulting in an overlap of 90 bp between adjacent windows). All window start positions are multiples of three to ensure that codon positions are aligned across windows. If the regular stride does not cover the end of the sequence, an additional window aligned to the sequence end is added, so that every position is covered by at least one window.

Each window is passed through the model up to, but not including, the CRF layer, producing pre-CRF logits for each shared label across reading frames. In regions covered by multiple windows, the logits are averaged position-wise across all contributing windows prior to decoding. The resulting full-length logit sequence is then decoded by the CRF in a single Viterbi pass to produce the final, most probable label sequence. Because predictions from overlapping windows are merged at the logit level prior to decoding, no conflicting label predictions arise between windows, and the CRF produces a single, consistent label sequence for the full input. As is the case for shorter sequences, the input sequence and its reverse complement are processed independently.

#### A1.2 Frequency of biologically invalid transitions in the CRF decoder

While DeepCDS is trained to strongly discourage biologically invalid transitions, it does not strictly prohibit them. To quantify how often they occur, we decoded the 300 bp typical-error test set (5,035,601 sequences from the 212 test genomes using the standard prokaryotic genetic code) with all three versions of DeepCDS and counted transitions between consecutive decoded labels that were initialized as invalid. No sequences contained such a transition with DeepCDS N, 3 sequences ($6 \cdot 10^{-5}\%$) with DeepCDS S, and 1 sequence ($2 \cdot 10^{-5}\%$) with DeepCDS S+I. In all four cases, a coding region switched directly from one reading frame to another without passing through a stop and start codon or, for DeepCDS S+I, an indel error-induced transition label.

#### A1.3 Loss function

During training, all models were trained to minimize the negative log-likelihood of the true label sequences [1]:

$$-\log(P(\mathbf{y}|\mathbf{H})) = \log(Z(\mathbf{H})) - \log\left(\exp\left(T_{\text{start}}(y_1) + \sum_{n=1}^{N} \varphi(\mathbf{h}_n, y_n) + \sum_{n=1}^{N-1} T(y_n, y_{n+1}) + T_{\text{end}}(y_N)\right)\right)$$

#### A1.4 Pairwise permutation tests for statistical comparison of model performance

To statistically assess the difference in performance between each pair of models, we conducted a pairwise permutation test at the read level, based on the F1 score aggregated across all 212 test genomes that use the standard prokaryotic genetic code. Each read contributes a set of TP, FP, and FN counts for each model, based on a 100% overlap between a labeled CDS and a predicted CDS (IoU = 1). Under the null hypothesis that two models perform equivalently, the assignment of prediction counts to each model is exchangeable within each read. For each permutation, the TP, FP, and FN counts were independently swapped between models per read with a probability of 0.5. Counts were then summed across all reads and the F1 score recomputed for each model, yielding a null distribution of F1 score differences across 1,000 permutations. The p-value was defined as the proportion of permutations that yielded an absolute F1 score difference at least as large as the observed difference in the actual F1 scores belonging to the two models. The minimum resolvable p-value is therefore $\frac{1}{1{,}000} = 0.001$. The p-values were corrected for multiple comparisons across all 21 (C(7,2)) model comparisons (benchmark models) and across all 36 (C(9,2)) model comparisons (ablation models) using Holm-Bonferroni correction. This procedure was repeated independently for each test set of combined error rate and sequence length condition.

#### A1.5 Training with unknown nucleotides

In the training set, we randomly "mask" the nucleotides in a sequence by replacing a given base with N at probability 0.05%, reflecting the low frequency of ambiguous base calls in short-read sequencing data. N is one-hot encoded as $[0, 0, 0, 0]$. This encourages the model to be robust towards unknown nucleotides that may occur during inference on real-world data. Codons containing an unknown nucleotide are encoded as unknown tokens (<unk>) with ESM-2 [2].

#### A1.6 Inference on sequence ends

Since DeepCDS encodes codons at the same position across all three reading frames into a single shared label, a challenge can arise at the 3' end of a sequence: a sequence may contain a complete codon in $RF_0$ at the sequence end, while $RF_1$ and $RF_2$ do not have complete codons at the same position due to their 1 and 2 nucleotide offsets, respectively. To handle this, the sequence is padded with unknown nucleotides (N) at the 3' end to ensure that each shared label position captures complete codon information across all three reading frames.

#### A1.7 Inference on sequences with unknown nucleotides

The allowed input vocabulary includes all standard IUPAC nucleotide codes. U is treated as equivalent to T. Characters outside the standard IUPAC vocabulary are converted to N (unknown). Any codon containing a nucleotide other than A, T, G, C or U, is encoded as an `<unk>`-token before being passed through the protein language model encoder component of the model. In the nucleotide-level one hot encoding, any other nucleotide than A, T, G, C and U is represented as an all-zero vector.

#### A1.8 DeepCDS Output Files

The output of DeepCDS is an annotation file (.gff) and corresponding fasta files with the predicted CDSs given as both nucleotide sequences (.fna) and corresponding amino acid sequences (.faa).

The .gff file annotates CDS regions (`type=CDS`), start codons (`type=start_codon`), and stop codons (`type=stop_codon`). If the user runs the variant of DeepCDS trained on sequences with both substitution and indel errors (DeepCDS S+I), sequencing artifacts caused by indel errors are annotated as well. CDS regions interrupted by an indel error are split into two or more `type=CDS` features and share a common `group_id` attribute, with `indel_type=insertion` or `indel_type=deletion` indicating the type of error detected (see Supplementary Note A1.9). A `type=uncertain_region` feature marks the ambiguous nucleotide positions between such fragments if any. In the cases where an inserted position is directly predicted, this position is marked by a `type=insertion` feature instead. CDS fragments belonging to the same indel-interrupted CDS share a common `ID` and are distinguished by their `group_id` attribute. All other predicted CDSs are assigned a unique `ID`. The `ID` is shared across the .gff, .fna, and .faa output files, enabling direct cross-referencing between them.

The predicted annotations are then used to produce the output fasta files holding the predicted CDSs. In the case of a deletion error, the missing region in the merged CDS sequence is represented as "NNN" in the .fna output file and translated as "X" in the .faa output file. Furthermore, all codons with one or more unknown nucleotide positions are translated as "X" in the .faa file. Stop codons are denoted as "*" in the .faa output file.

#### A1.9 Definition of Insertion and Deletion errors

DeepCDS S+I detects indel errors by leveraging reading frame shifts. Because indel errors are rarely introduced by modern short read sequencing platforms, the model assumes that two indel errors will not occur at consecutive positions. If a reading frame shift for a CDS is detected to increase by one (e.g. from $RF_0$ to $RF_1$), DeepCDS S+I predicts that a base has been inserted. In contrast, if the reading frame decreases by one (e.g. from $RF_1$ to $RF_0$), a deletion is predicted. Note that this assumption means the model cannot reliably handle consecutive indel errors.

#### A2 Supplementary Figures

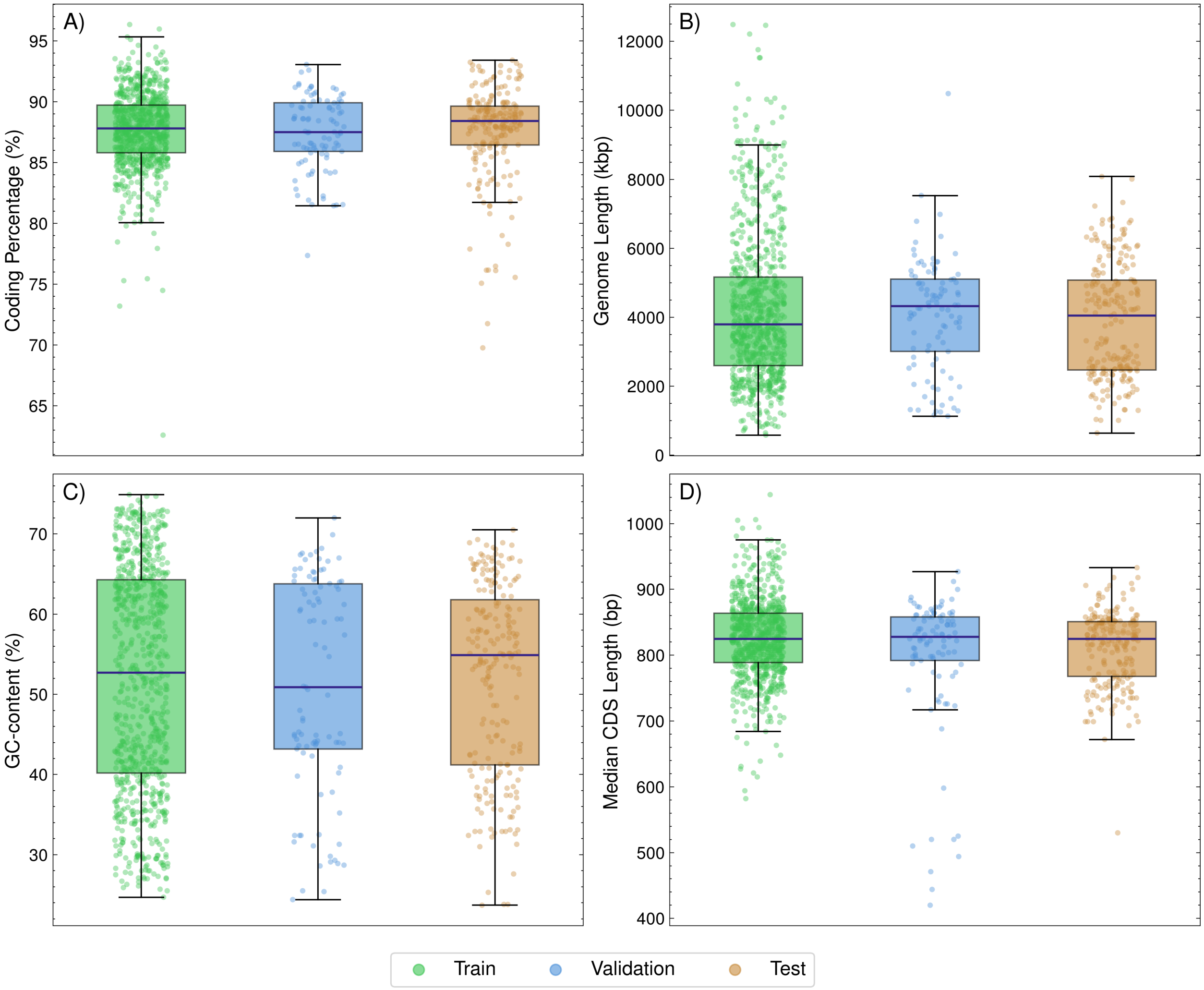


Figure A1: Genome-level statistics based on data from the 1,125 genomes in the full dataset, distributed on the training, validation, and test partition. Each point corresponds to a genome. A) Distribution of protein-coding percentage. The protein-coding percentage is measured as the percentage of positions in the genome, covered by a CDS annotated with RefSeq. B) Distribution of genome size in kilo base pairs based on the RefSeq-assembled genomes. C) Distribution of genomic GC-contents. D) Distribution of median CDS length, based on the RefSeq-annotated CDS lengths.

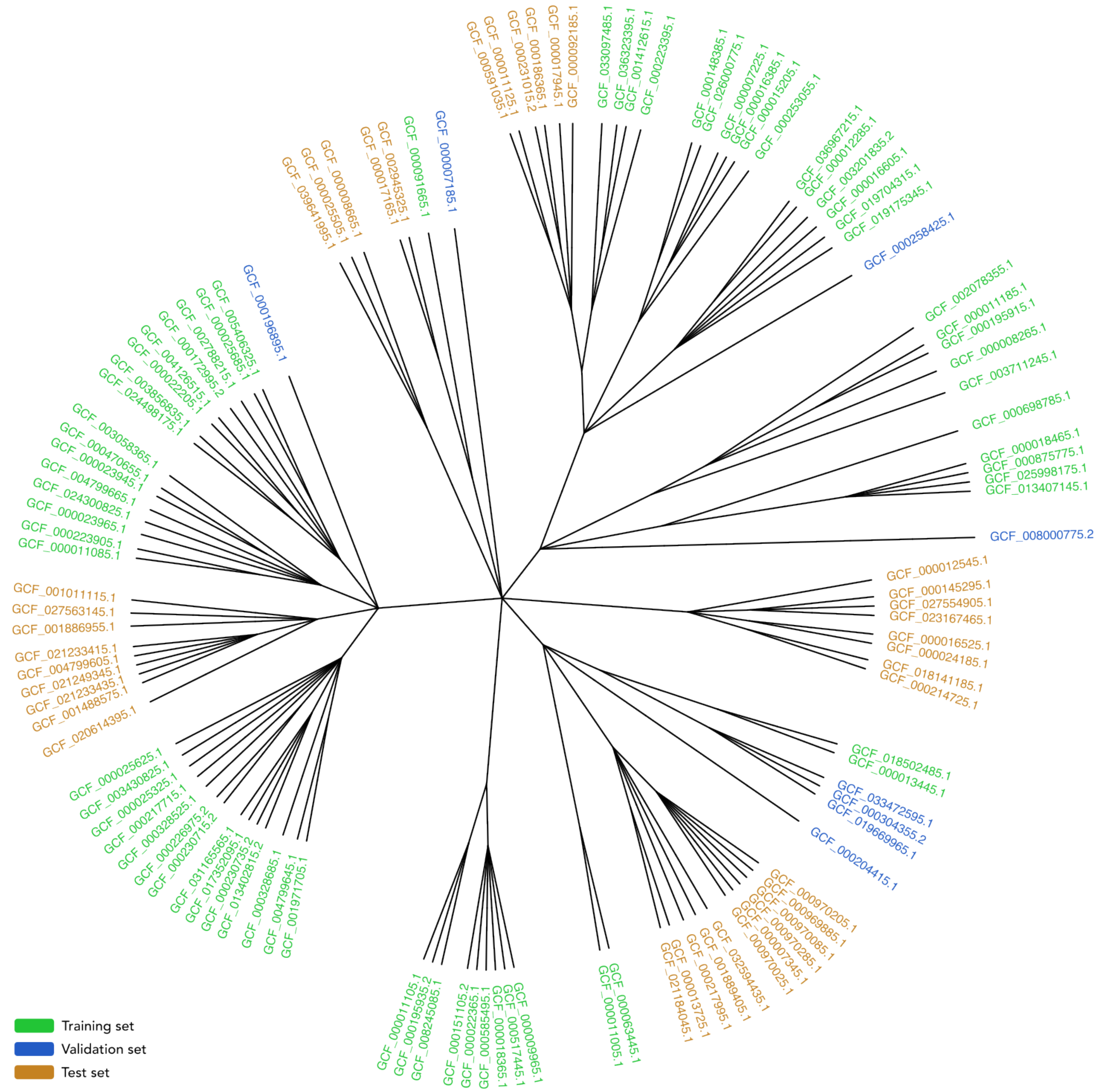

Figure A2: Hierarchical taxonomic tree of the 117 archaeal organisms in the full dataset, colored based on data partition. The training, validation, and test sets include 70, 8, and 39 archaeal genomes, respectively. Genomes were partitioned at the family level, with all members of each family assigned to the same partition to prevent data leakage.

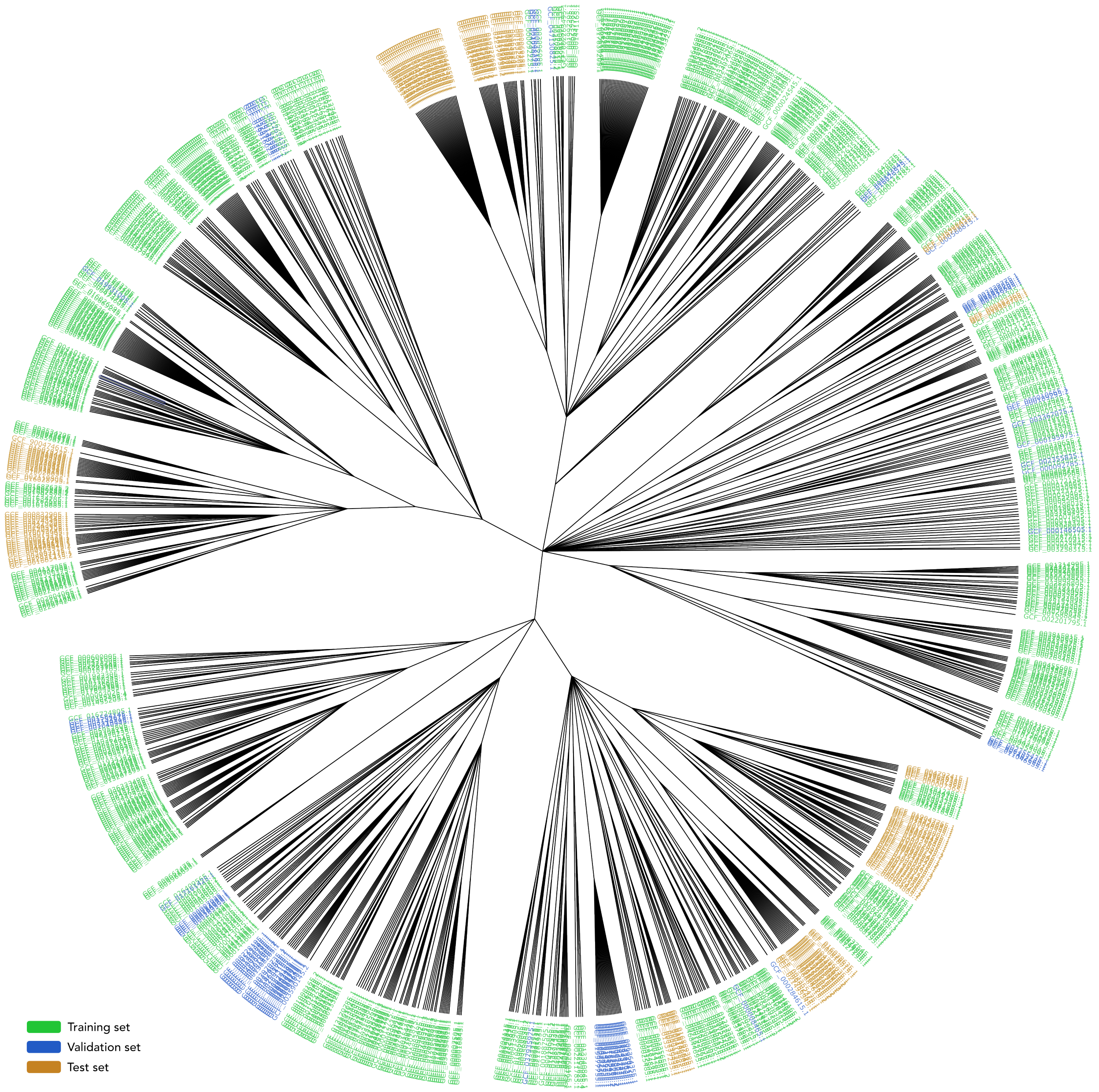


Figure A3: Hierarchical taxonomic tree of the 1,008 bacterial organisms in the full dataset, colored based on data partition. The training, validation, and test sets include 743, 89, and 176 bacterial genomes, respectively. Genomes were partitioned at the family level, with all members of each family assigned to the same partition to prevent data leakage.

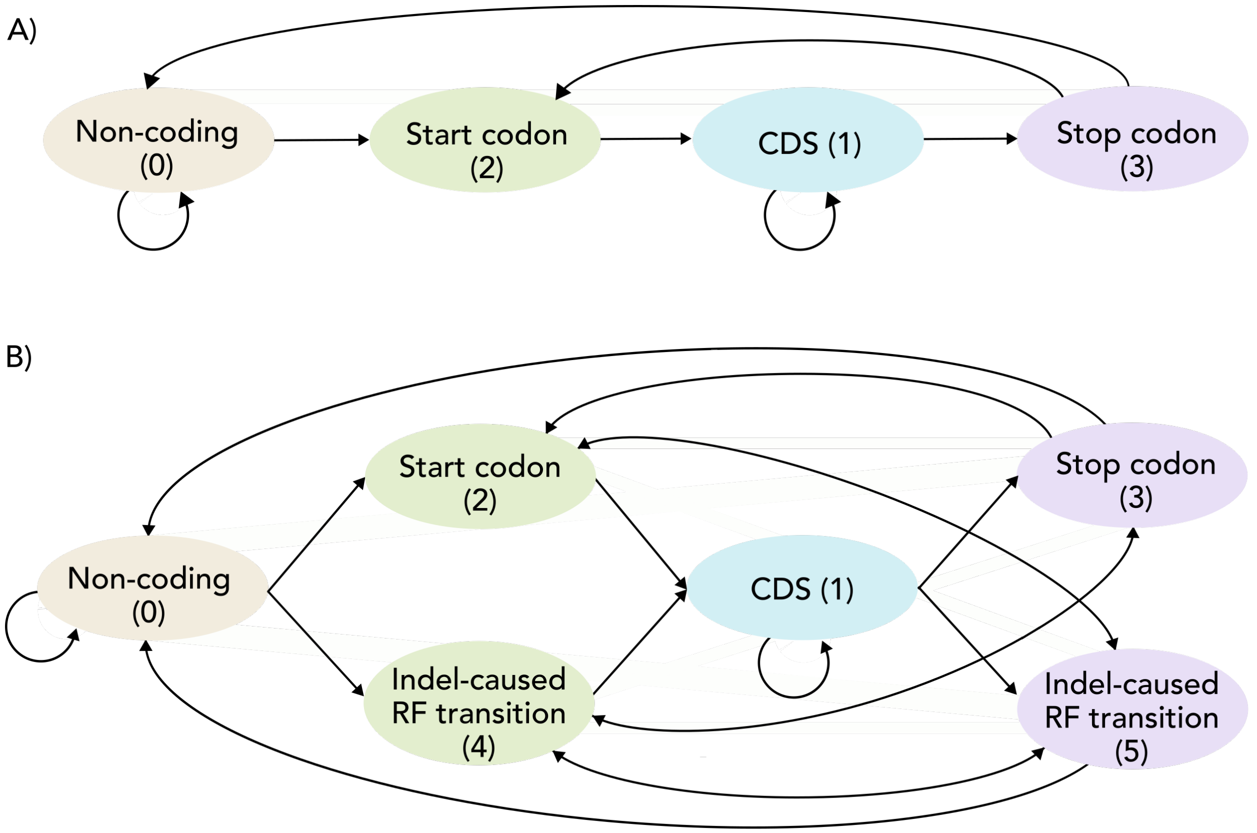


Figure A4: The pre-defined RF-specific codon label state diagrams for all DeepCDS models. All transitions marked by arrows are the biologically valid ones within a single RF. The invalid transitions, not marked by an arrow, were initialized with a large negative value of $-10$ in the shared $K \times K$ label space transition matrix. For example, the $(\mathrm{RF}_0,\, \mathrm{RF}_1,\, \mathrm{RF}_2)_l = (0,1,0) \rightarrow (\mathrm{RF}_0,\, \mathrm{RF}_1,\, \mathrm{RF}_2)_{l+1} = (1,1,0)$ would be an invalid transition and initialized to $-10$, because the label state in $\mathrm{RF}_0$ cannot enter the CDS space (1) without going through a start codon (2) or an indel error-caused RF transition (4). The green states mark entry into a CDS, and the purple states mark exit from a CDS. **A)** Label state transition diagrams for the models trained on data without indel errors ($|C| = 4$), namely DeepCDS N and DeepCDS S. **B)** Label state transition diagrams for the models trained on data with indel errors ($|C| = 6$), namely DeepCDS S+I.

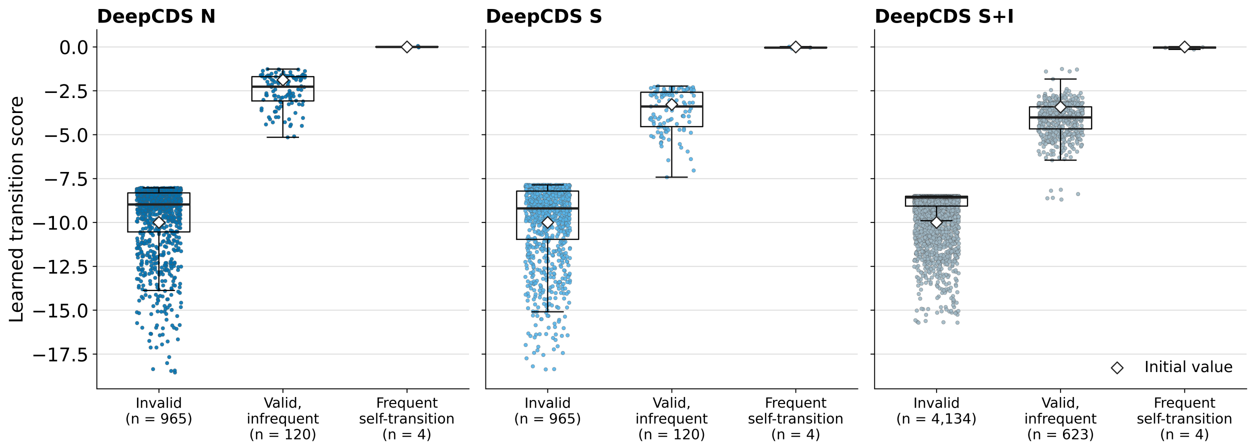


Figure A5: Learned transition scores of the trained CRF for each DeepCDS model, grouped by their initialization. “Invalid” refers to biologically invalid transitions, based on the label state diagrams in Supplementary Figure A4, initialized at $-10$. “Valid, infrequent” refers to all valid transitions not considered frequent, initialized at the tuned transition weight ($-1.88$, $-3.28$, and $-3.42$ for DeepCDS N, S, and S+I, respectively). “Frequent self-transition” refers to transitions where either all RFs go from non-coding to non-coding, or one RF goes from CDS to CDS while the other two go from non-coding to non-coding, initialized at 0. DeepCDS N and S use $|C| = 4$ codon label classes and $K = 33$ shared label states observed in the training data, giving $33 \cdot 33 = 1,089$ transition scores; DeepCDS S+I uses $|C| = 6$ and $K = 69$, giving $69 \cdot 69 = 4,761$. Points show individual transitions. The six lowest-scoring valid transitions of DeepCDS S+I (below $-8$) correspond to a CDS leaving a reading frame due to an indel error and, at the next codon of the same frame, re-entering it due to an indel error or starting at a start codon; these are biologically possible but did not occur in the training data.

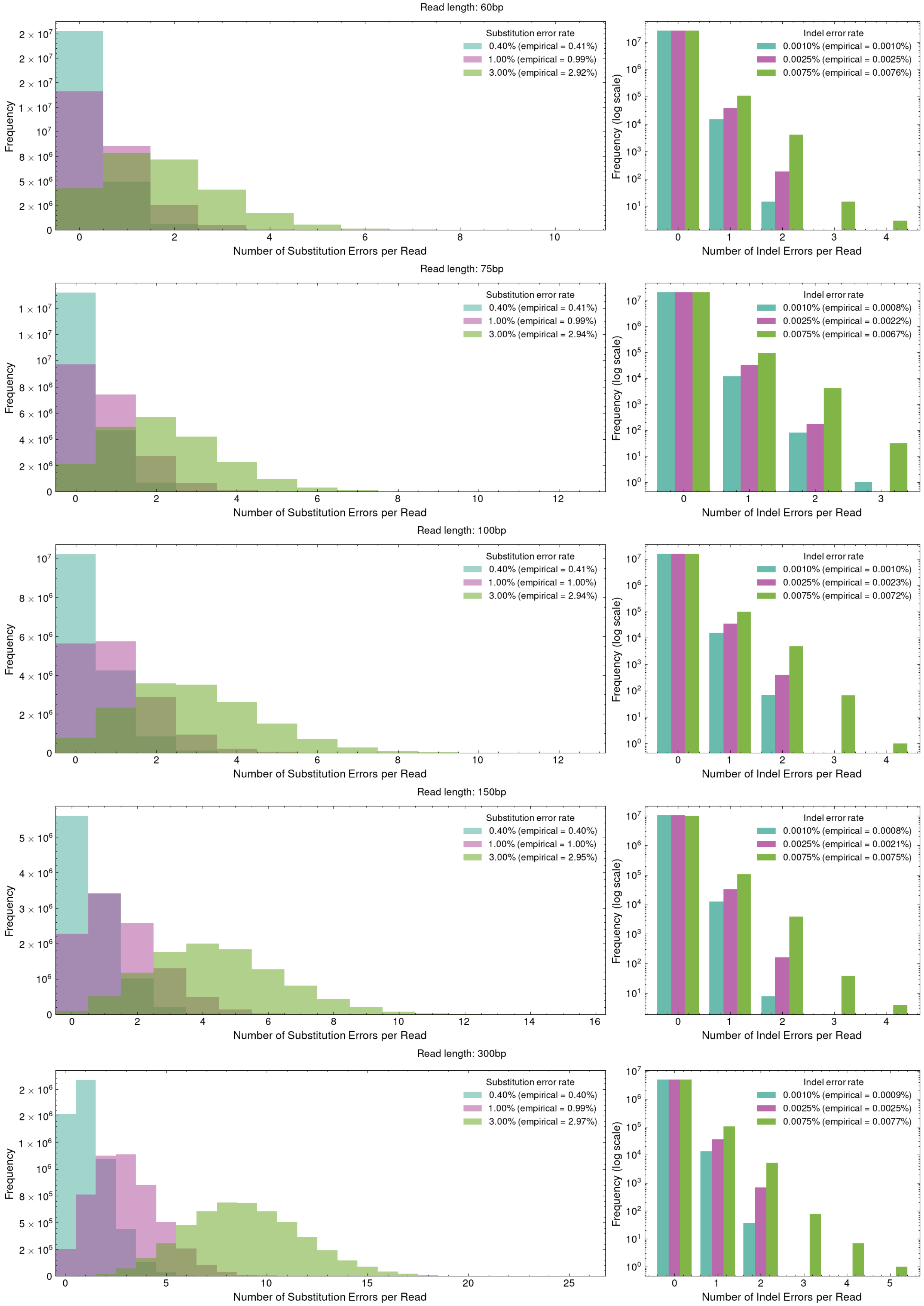


Figure A6: Distributions of sequencing errors per read for each test set of different read length and error rate simulated with Mason [3]. The empirical rates refer to the actual error rates present in the simulated test sets.

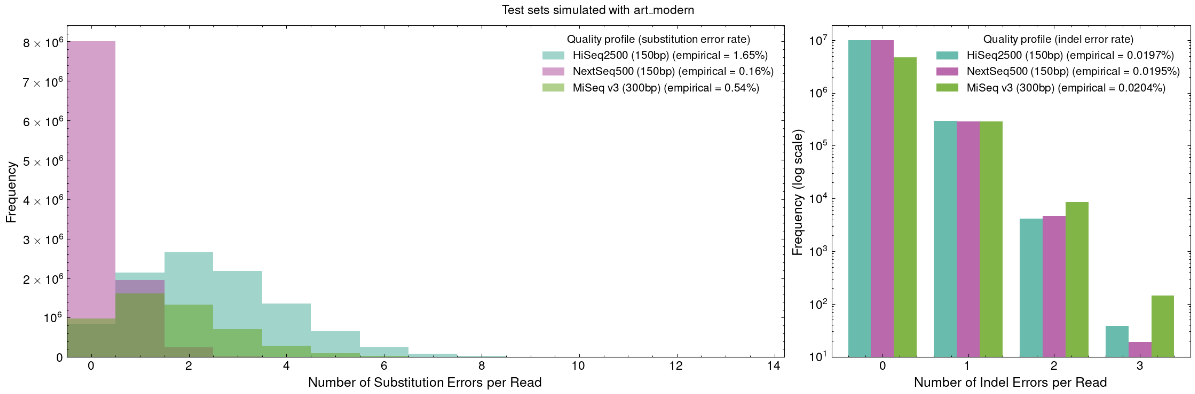


Figure A7: Distributions of sequencing errors per read for the three simulated test sets with different quality profiles from art_modern [4]. The empirical rates refer to the actual error rates present in the simulated test sets.

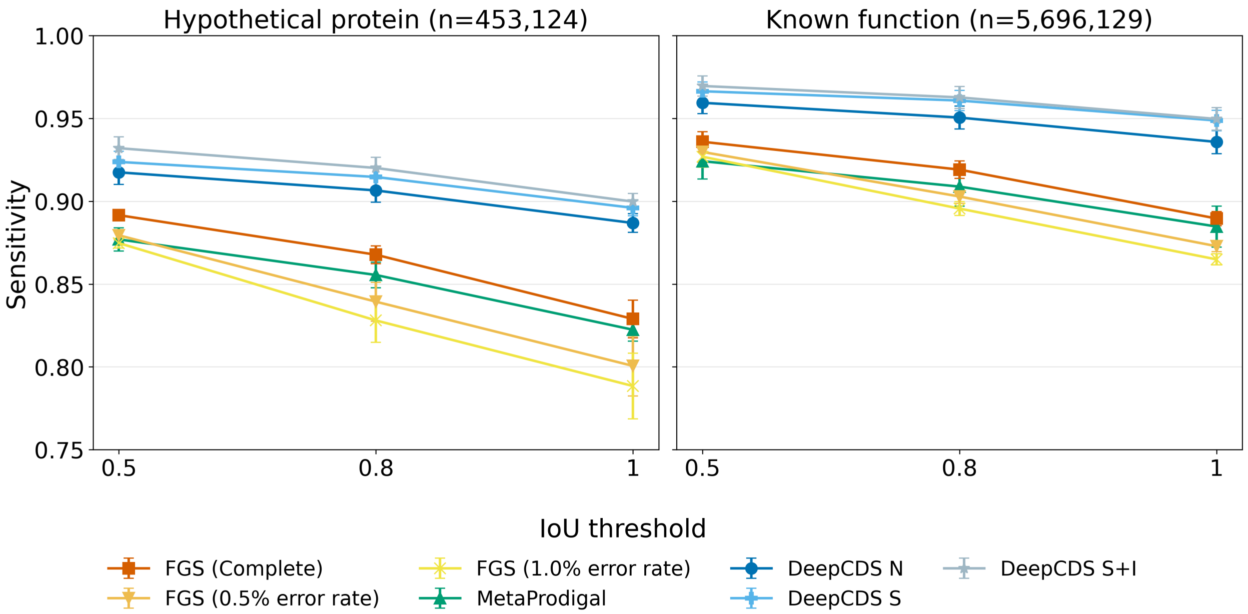


Figure A8: CDS-level sensitivity on CDS fragments from hypothetical proteins compared to CDS fragments with an annotated molecular function in the CAMI III sample ("Toy Longitudinal Human Gut"), after removing genomes overlapping with the training and validation sets. The set of hypothetical proteins shown here comprise CDS annotations with the attribute "product=hypothetical protein" that were not tagged as ab initio predictions (n=453,124 CDS fragments across 452,202 reads). CDSs with an annotated molecular function comprise CDS annotations with a "go_function" attribute (n=5,696,129 CDS fragments across 5,676,352 reads). Performance is shown for exact coordinate matching and for IoU overlap thresholds of 0.8 and 0.5. The error bars report the standard error from a genome-level cluster bootstrap with 1,000 replicates.

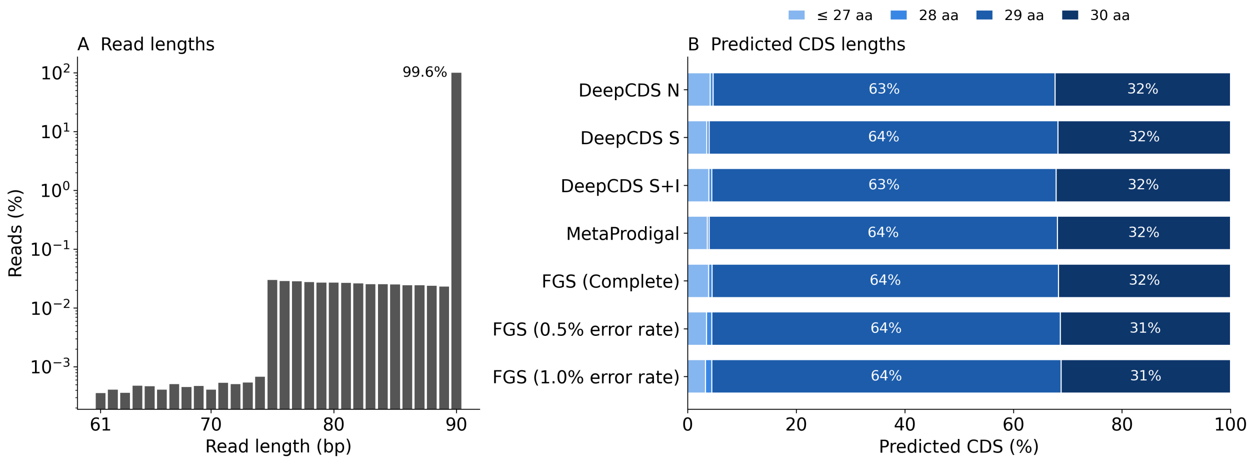


Figure A9: Read and predicted CDS lengths on the real gut metagenome sample MH0161 from the Danish MetaHIT cohort. A) Lengths of the 58.4M reads >60 bp after preprocessing ($\log_{10}$ scale); 99.6% are exactly 90 bp. B) Lengths of the predicted CDSs (>60 bp on either strand) per tool, as the share of each tool's predicted CDSs.

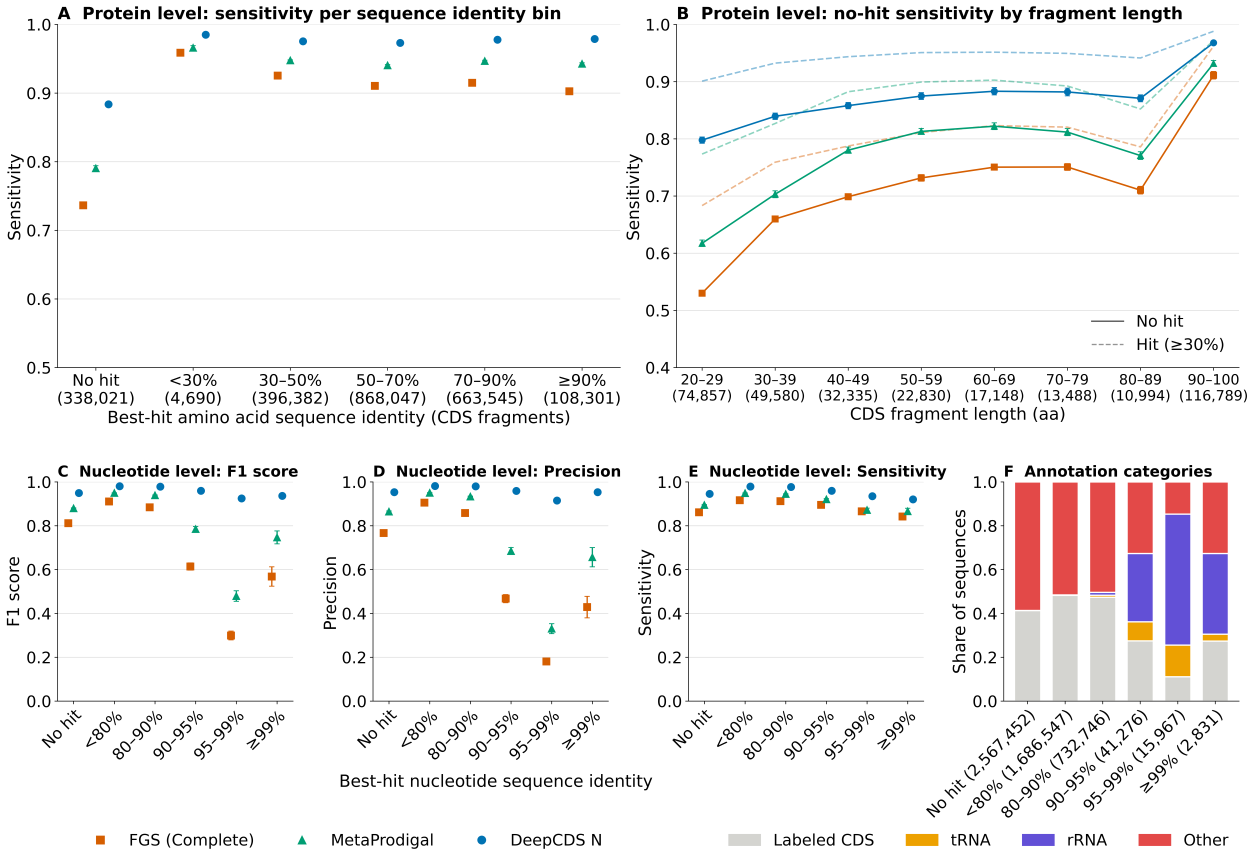


Figure A10: Performance stratified by homology to the training data. Test set performance stratified by the sequence identity of each test sequence to the training and validation sequences, based on the best hit by bitscore using MMseqs2 [5] ($E \leq 10^{-3}$). The results are based on the 300 bp error-free test set of the 212 genomes using the standard prokaryotic genetic code. "No hit" corresponds to no detected alignment of at least 20 amino acids (protein level) or 60 nucleotides (nucleotide level). Performance is CDS-level with exact matching of predicted and labeled CDSs. **(A)** Protein-level sensitivity of labeled CDS fragments (>60 bp), binned by the amino acid sequence identity of the best hit among the training and validation CDS fragments. **(B)** Sensitivity of "No hit" fragments by fragment length (solid lines). Fragments with a hit of $\geq$30% identity are shown for reference (dashed lines). **(C–E)** Nucleotide-level F1 score, precision, and sensitivity, with test sequences binned by the nucleotide sequence identity of their best hit among the training and validation sequences; all test sequences are included, also those without a labeled CDS. **(F)** Share of test sequences per nucleotide identity bin that contain a labeled CDS fragment, or otherwise lie within an annotated rRNA or tRNA gene (overlap >60 bp, either strand). Sequences were simulated from both strands and are labeled only with CDSs on their own strand; "Other" therefore also comprises sequences antisense to a CDS. Numbers in parentheses give the number of CDS fragments (A, B) or sequences (F; the same counts apply to C–E) per bin. Error bars mark one standard error from a genome-level cluster bootstrap (1,000 replicates).

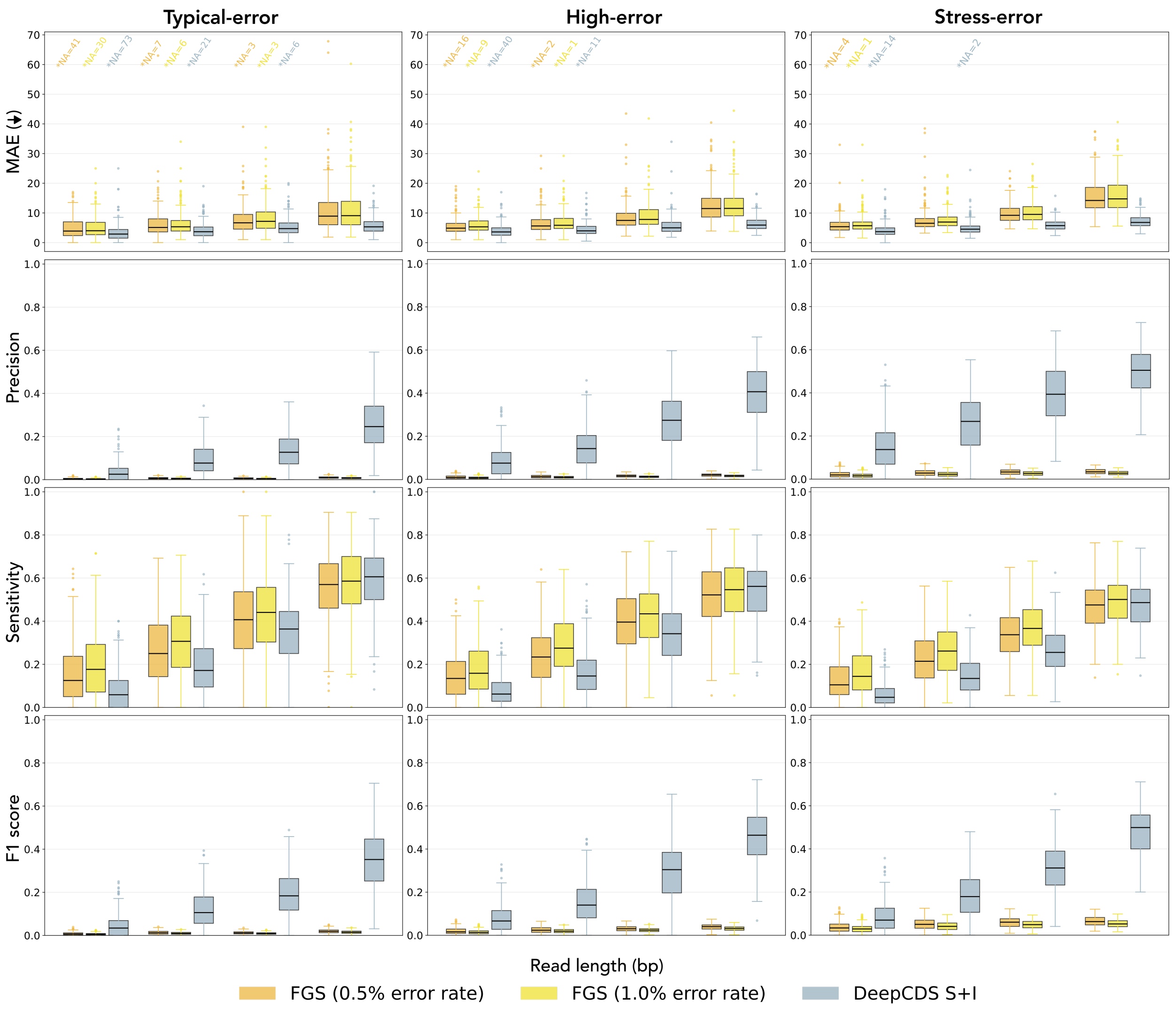


Figure A11: Boxplots of indel error detection performance across the typical-error, high-error, and stress-error test sets for the models trained to explicitly predict indel errors. Each point represents one of the 212 test genomes that follow the standard prokaryotic genetic code. Within each read, predicted indels are matched one-to-one to true indel errors in order of increasing distance. Precision, sensitivity, and F1 score are computed at the indel level, where a TP is a matched predicted indel error located within ±10 bp of the true indel error site. Predicted indel errors without a TP match, including those on reads without an actual indel error, count as FPs, and true indel errors without a TP match count as FNs. The MAE reports the mean absolute distance (in bp) between all matched true and predicted indel positions, independent of the positional tolerance. *NA denotes the number of genomes with no matched indel pairs, for which the MAE is undefined. Note that precision, sensitivity, and F1 score only reflect positional accuracy up to the ±10 bp tolerance; finer positional accuracy is instead reported by the MAE. All metrics are computed on indel-interrupted CDSs longer than 60 bp.

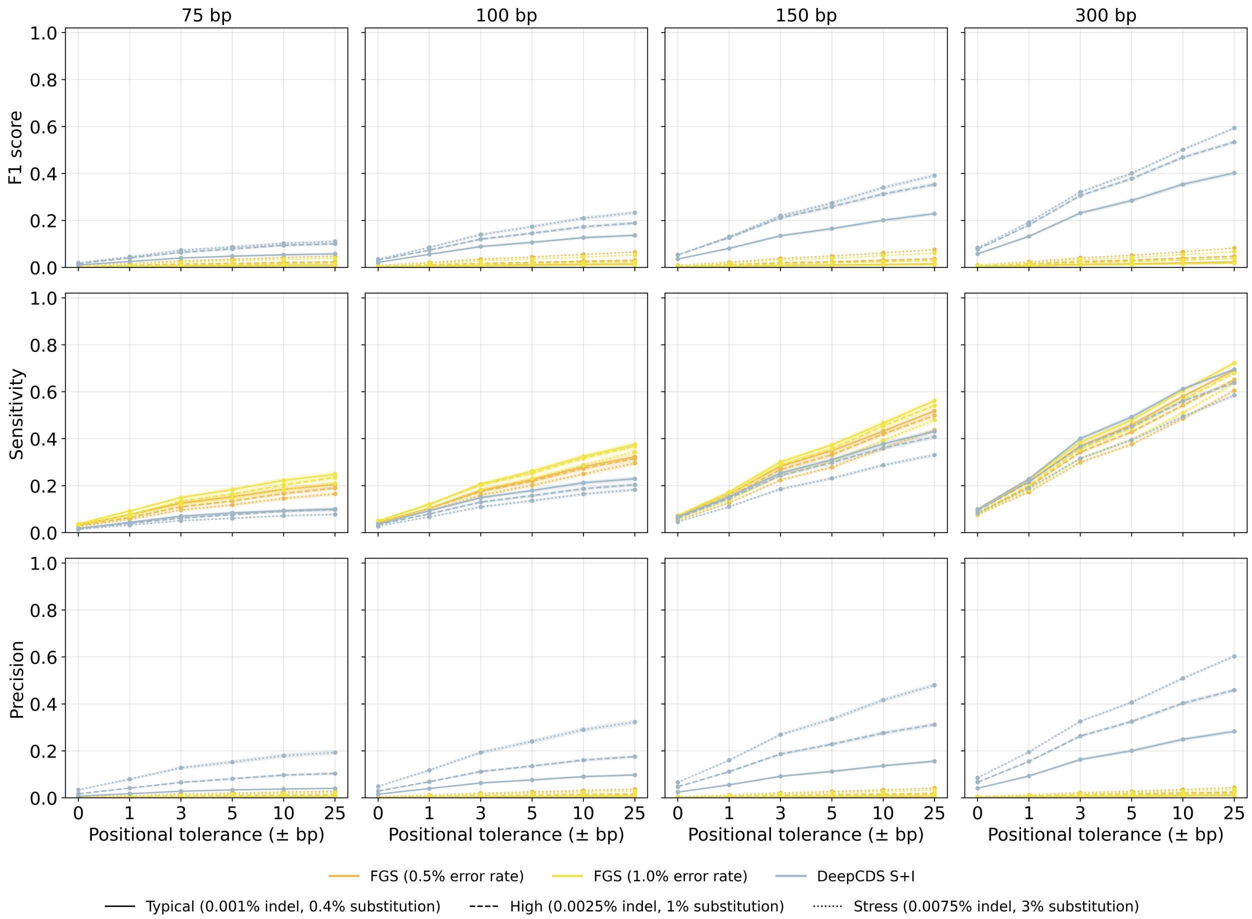


Figure A12: Indel error detection performance (F1 score, precision, and sensitivity) as a function of the positional tolerance used to define a TP, for the models trained to explicitly predict indel errors. Line styles denote the typical-error, high-error, and stress-error test sets. Predicted indels are matched one-to-one to true indels within each read, and a match counts as a TP if it lies within the given tolerance of the true indel error site (0 bp: exact position match; 25 bp: within ±25 bp). Predicted indels without a TP match, including those on reads without an actual indel error, count as FPs, and true indels without a TP match count as FNs. Metrics are computed from TP, FP and FN counts aggregated across the 212 test genomes following the standard prokaryotic genetic code. The shaded bands show the standard error from a genome-level cluster bootstrap with 1,000 replicates.

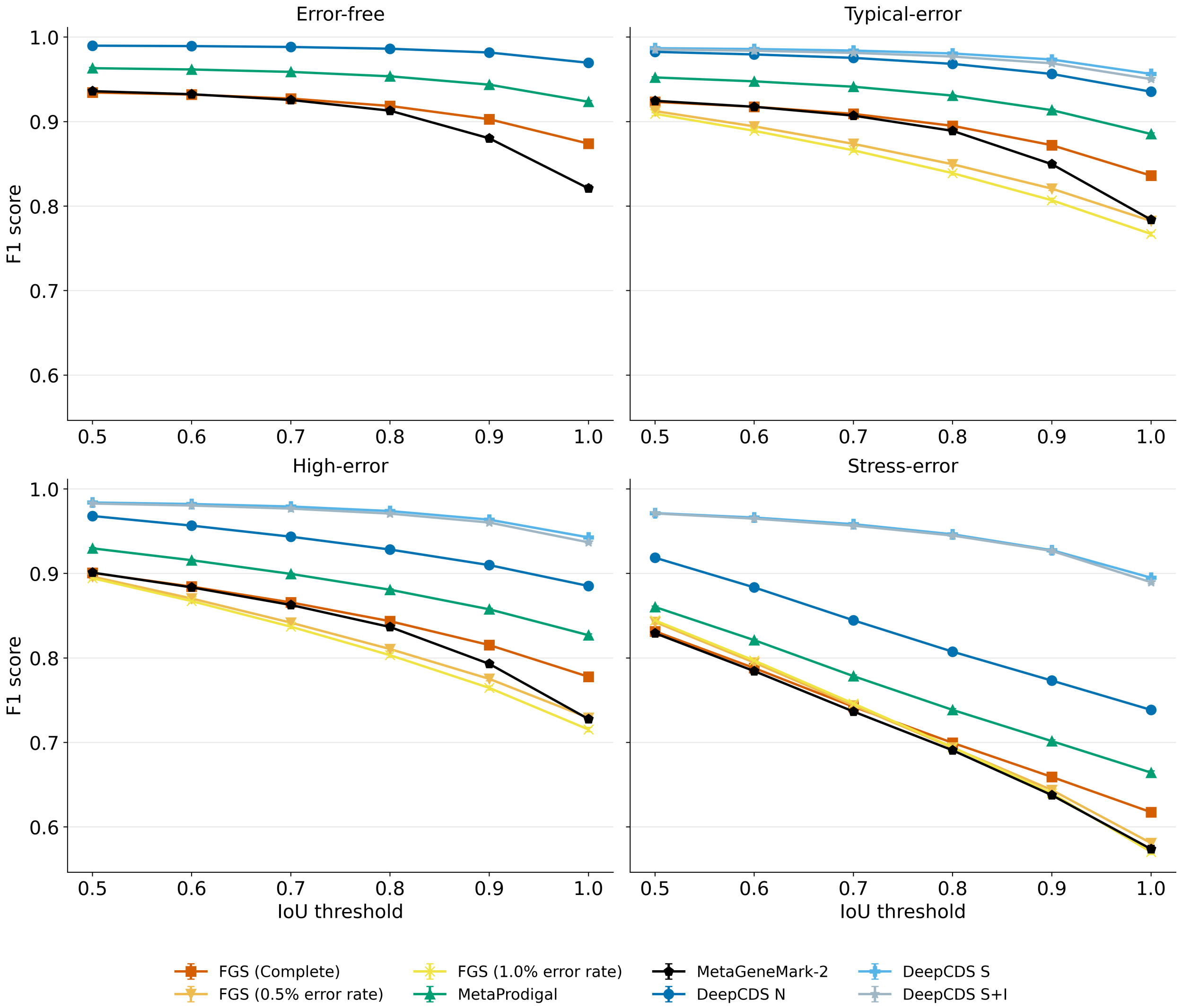


Figure A13: CDS-level F1 score as a function of the IoU threshold for counting a predicted CDS as a TP, on the 300bp error-free, typical-error, high-error and stress-error test sets. An IoU of 1 corresponds to exact matching of the predicted and ground truth CDS coordinates. Results are aggregated across the 212 test genomes using the standard prokaryotic genetic code. For a fair comparison to MetaGeneMark-2, which cannot predict CDSs shorter than 90 bp, only ground truth and predicted CDSs of at least 90 bp are included. Error bars show the genome-level cluster bootstrap standard error (1,000 replicates) and are mostly smaller than the markers.

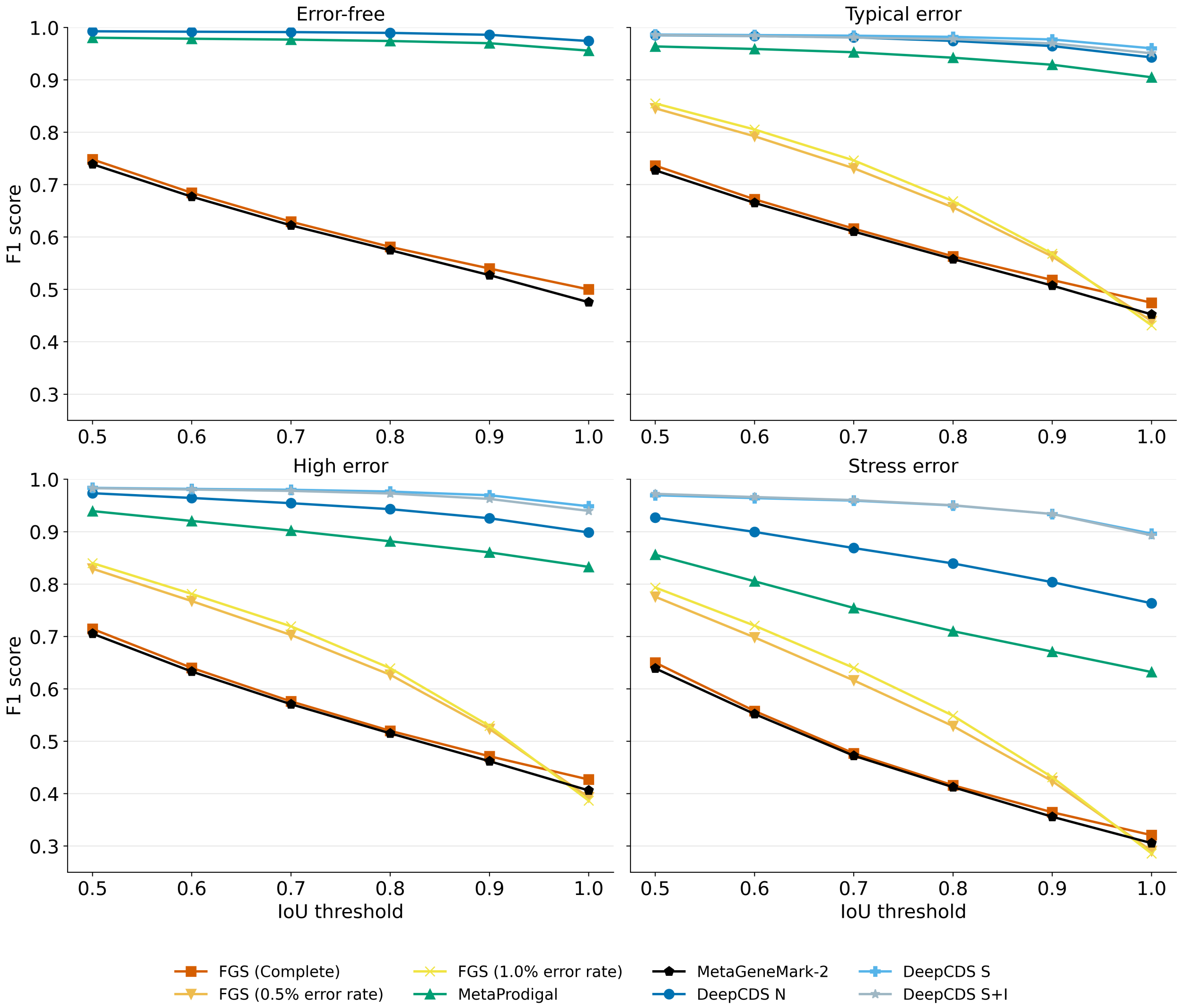


Figure A14: CDS-level F1 score as a function of the IoU threshold for counting a predicted CDS as a TP, on the 300 bp error-free, typical-error, high-error and stress-error test sets. An IoU of 1 corresponds to exact matching of the predicted and ground truth CDS coordinates. Results are aggregated across the 3 *Mycoplasmataceae* test genomes using genetic code 4. For a fair comparison to MetaGeneMark-2, which cannot predict CDSs shorter than 90 bp, only ground truth and predicted CDSs of at least 90 bp are included. No standard errors are shown, as a genome-level bootstrap over three genomes is not meaningful.

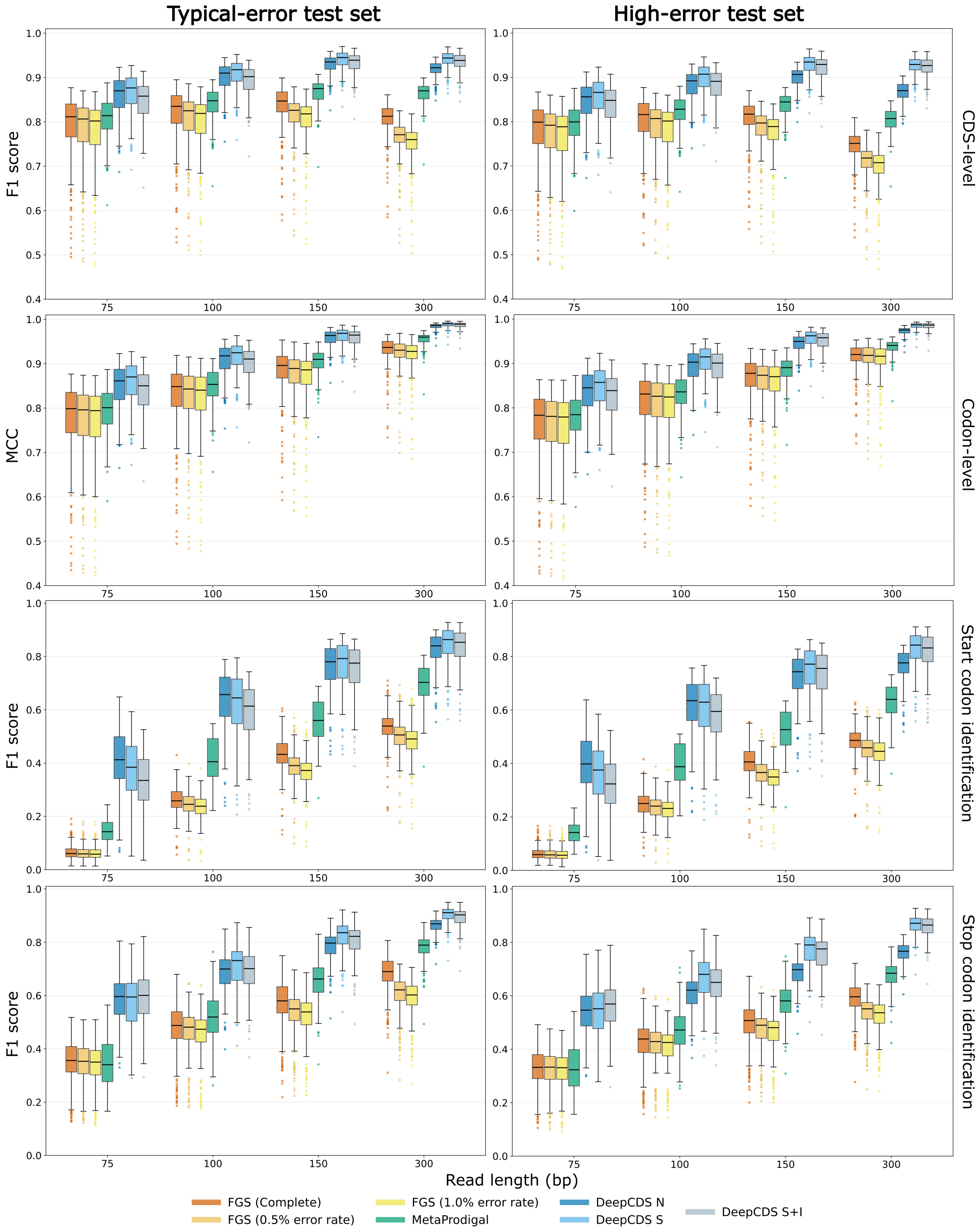


Figure A15: Boxplots visualizing a range of performance metrics for the typical-error and high-error test sets at different sequence lengths and measured at the genome level for the 212 test genomes that follow the standard prokaryotic genetic code. The results are based on all CDS fragments longer than 60 bp. Each row shows a different performance metric: (1) the CDS-level F1 score, based on 100% overlap between predicted and ground truth CDS coordinates. (2) the codon-level MCC, where a codon is a TP if it is both coding and predicted as coding, etc. (3) start codon identification F1 score. (4) stop codon identification F1 score. Note that the y-axes for the CDS-level and codon-level performance are shown in the range $[0.4, 1.0]$.

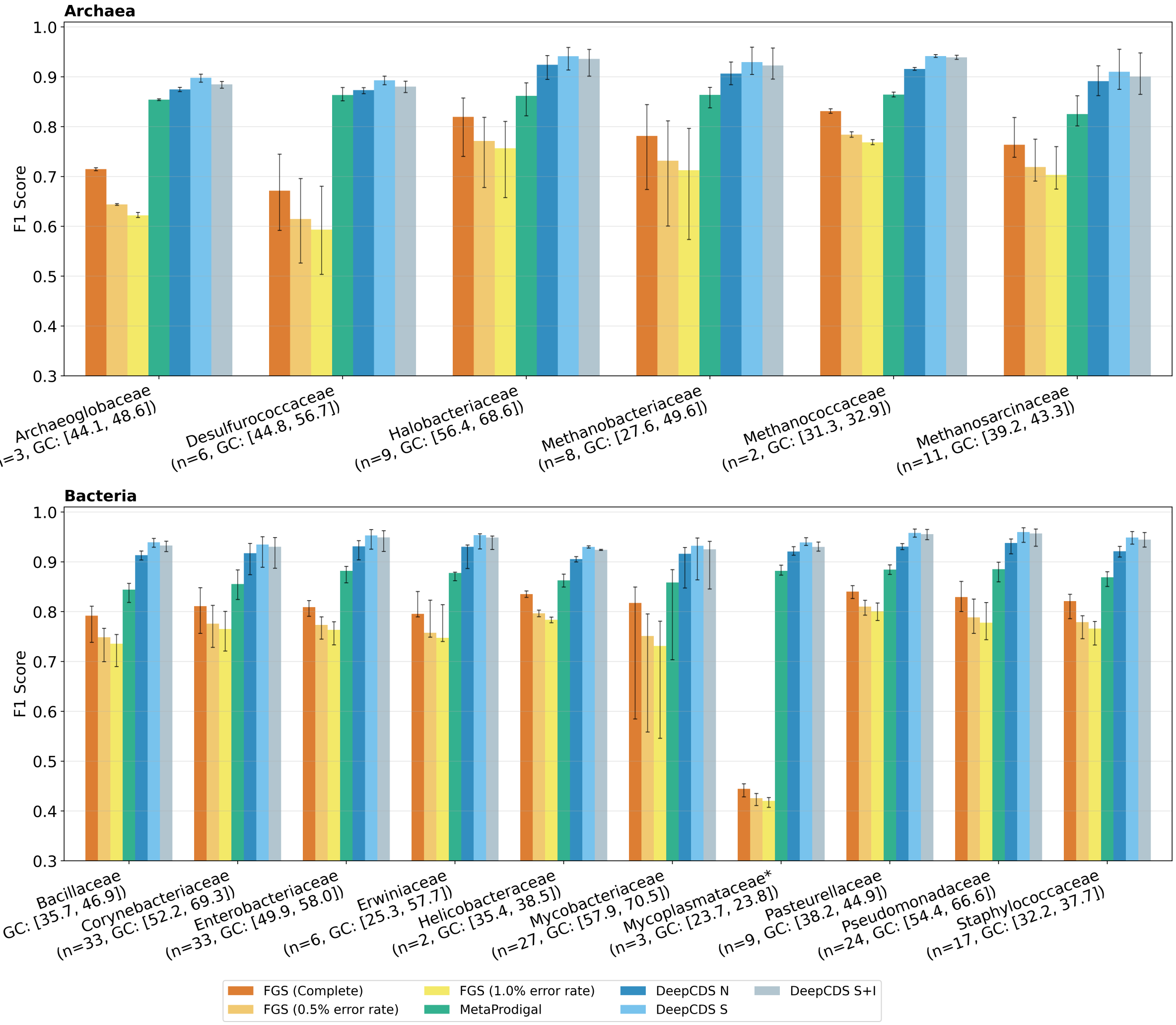


Figure A16: CDS-level F1 score aggregated per family on the 300bp typical-error test set. The error bars demonstrate the per-genome variation in F1 score within each family. n denotes the count of genomes within each test set family. GC denotes the range in GC-content for the genomes within that family. The $*$ represents usage of an alternative genetic code (NCBI genetic code 4). Note that the y-axis is shown in the range $[0.3, 1.0]$.

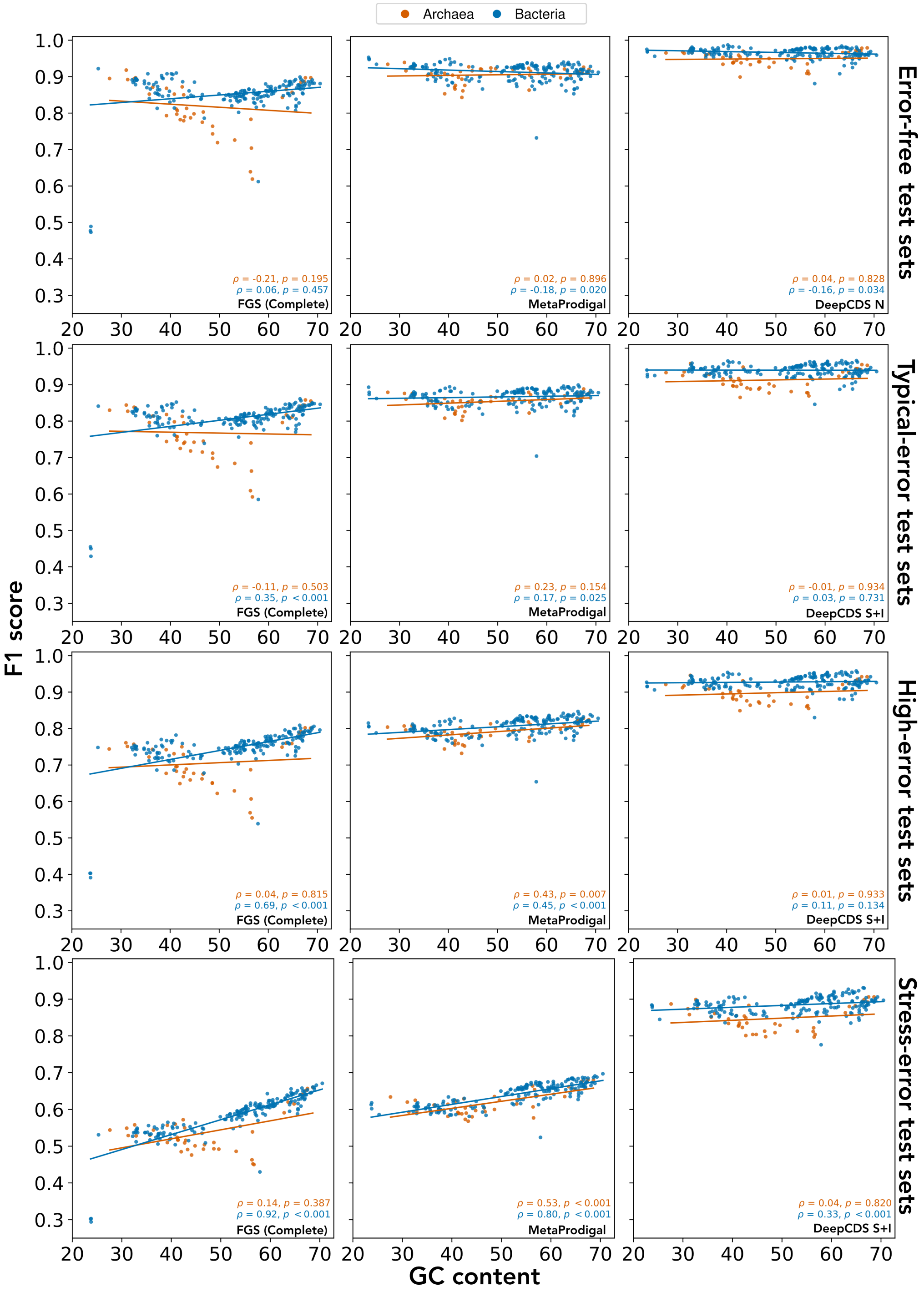


Figure A17: CDS-level F1 scores per genome as a function of genomic GC-content for all four error-rate conditions at 300bp read length. $\rho$ denotes the Spearman correlation for archaeal and bacterial genomes, respectively. Each point represents one of the 215 test genomes, including the 3 organisms that follow a genetic code alternative to the standard prokaryotic one. We show scatterplots for one representative model variant from each of FGS and DeepCDS. The Spearman correlations ($\rho$) for all model variants are shown in Table A30.

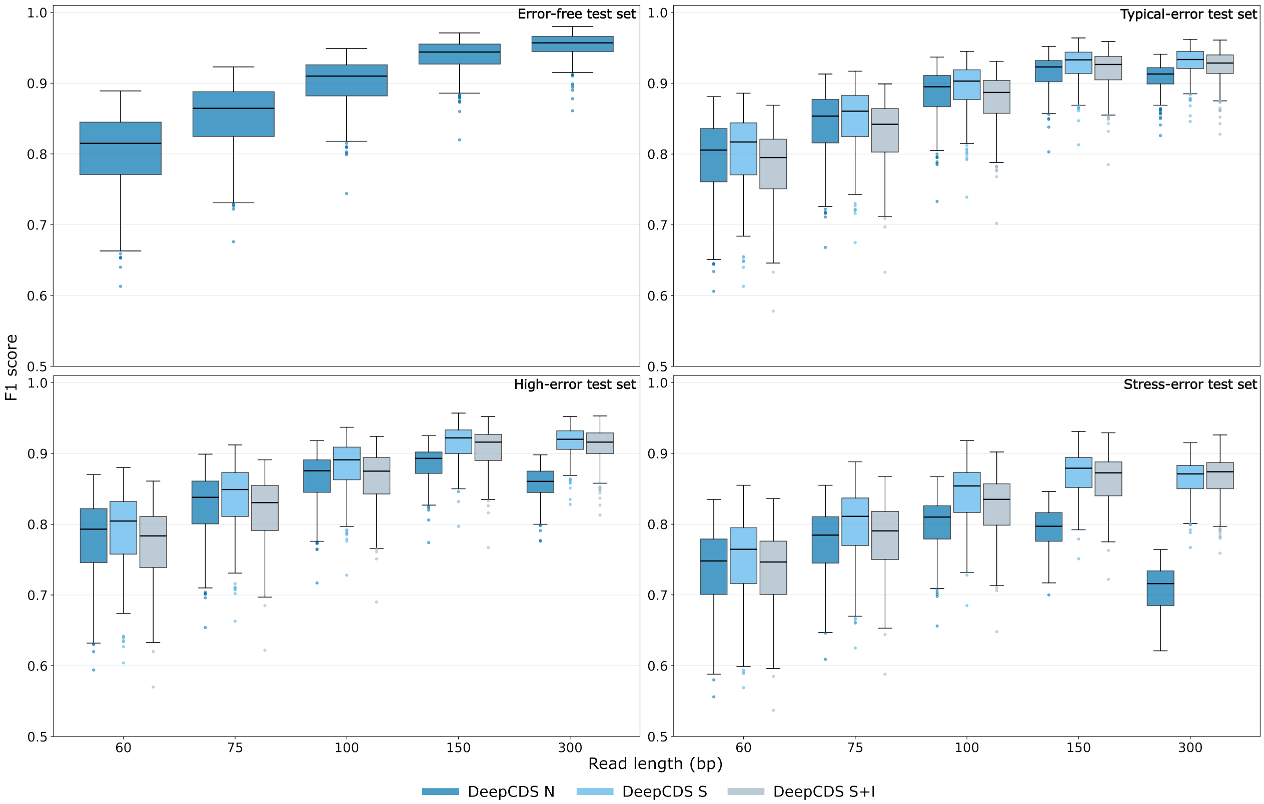


Figure A18: Boxplots visualizing the CDS-level F1 scores measured at the genome level for the test sets of each sequence length and error rate condition, based on CDS fragments longer than 30bp. Each boxplot is based on the F1 scores obtained from each of the 212 test genomes that follow that standard prokaryotic translation table. Note that the y-axes are shown in the range [0.5;1.0]. FragGeneScan and MetaProdigal predict fragments of length 60bp or longer, and are thus not included in this figure.

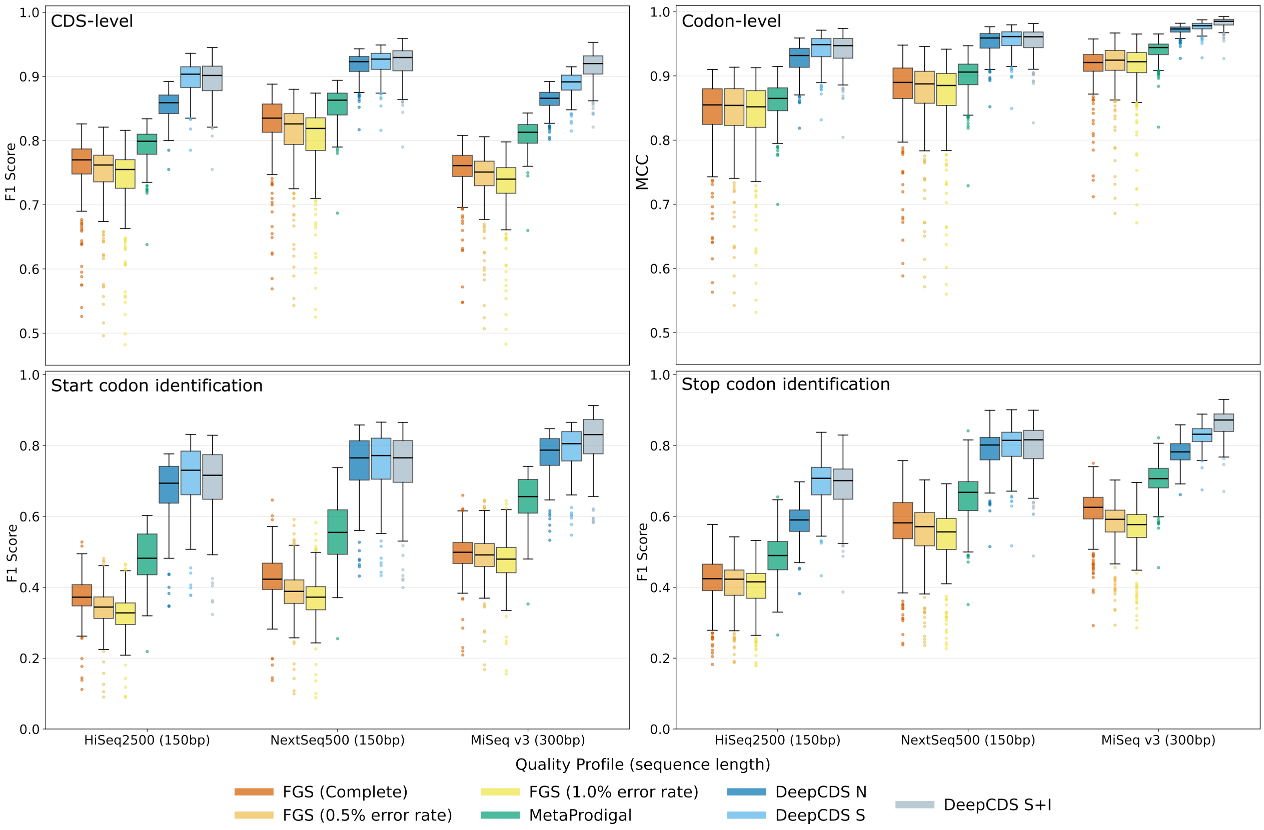


Figure A19: Boxplots visualizing performance metrics measured on the genome level for the test sets simulated with art_modern, testing three built-in quality profiles [4]. The results are based on all CDS fragments longer than 60 bp from the simulated reads from the 212 test genomes that follow the standard prokaryotic genetic code. Each boxplot shows one performance measurement: CDS-level F1 score, codon-level MCC, start codon F1 score, and stop codon F1 score. Note that the y-axes for the CDS-level and codon-level performance are shown in the range $[0.45, 1.0]$.

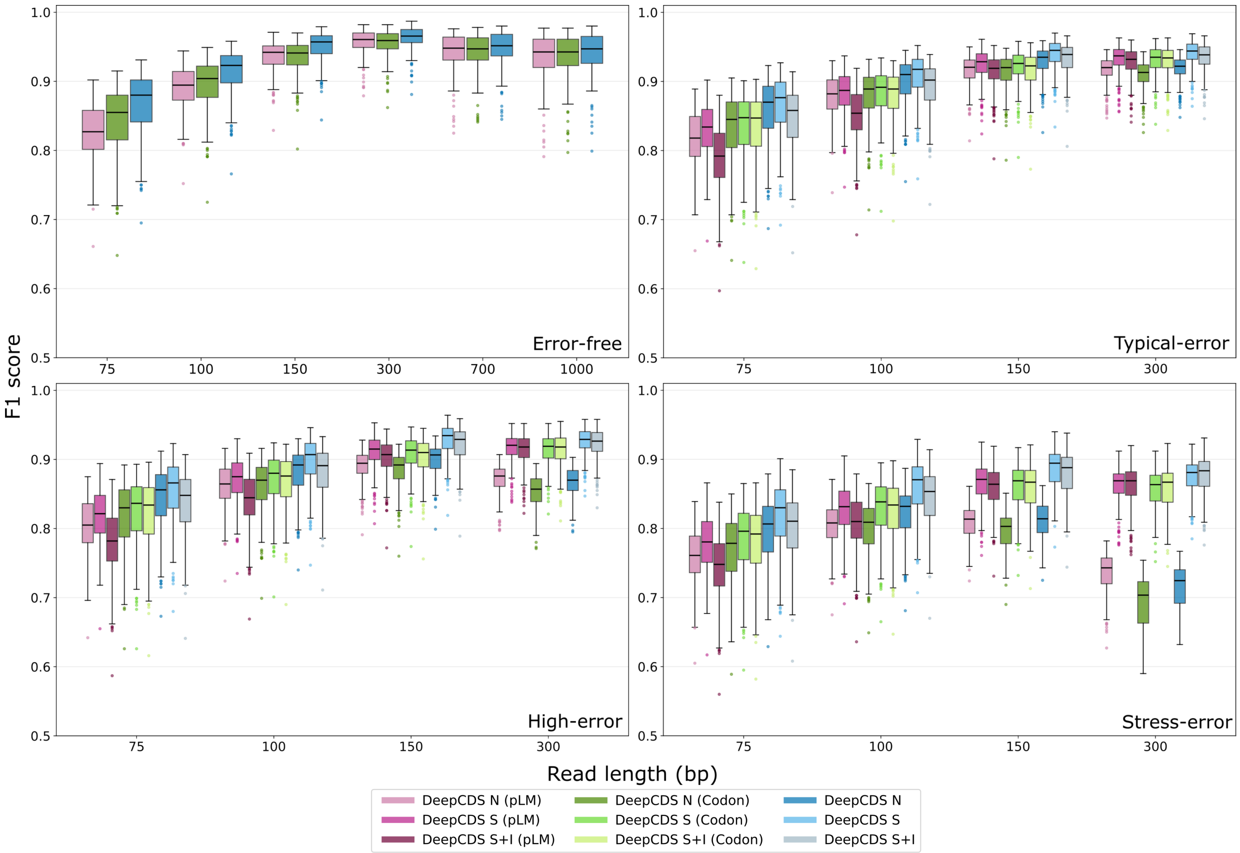


Figure A20: Boxplots visualizing the CDS-level F1 scores measured at the genome level for the test sets of each sequence length and error rate condition for each of the DeepCDS ablation models. Each boxplot is based on the F1 scores obtained from each of the 212 test genomes that follow the standard prokaryotic translation table. The results are based on all CDS fragments longer than 60 bp. Note that the y-axes are shown in the range [0.5;1.0]. (Codon) refers to the ablation models using only the codon encoding as input, and (pLM) refers to the ablation models using only the pLM embeddings as input.

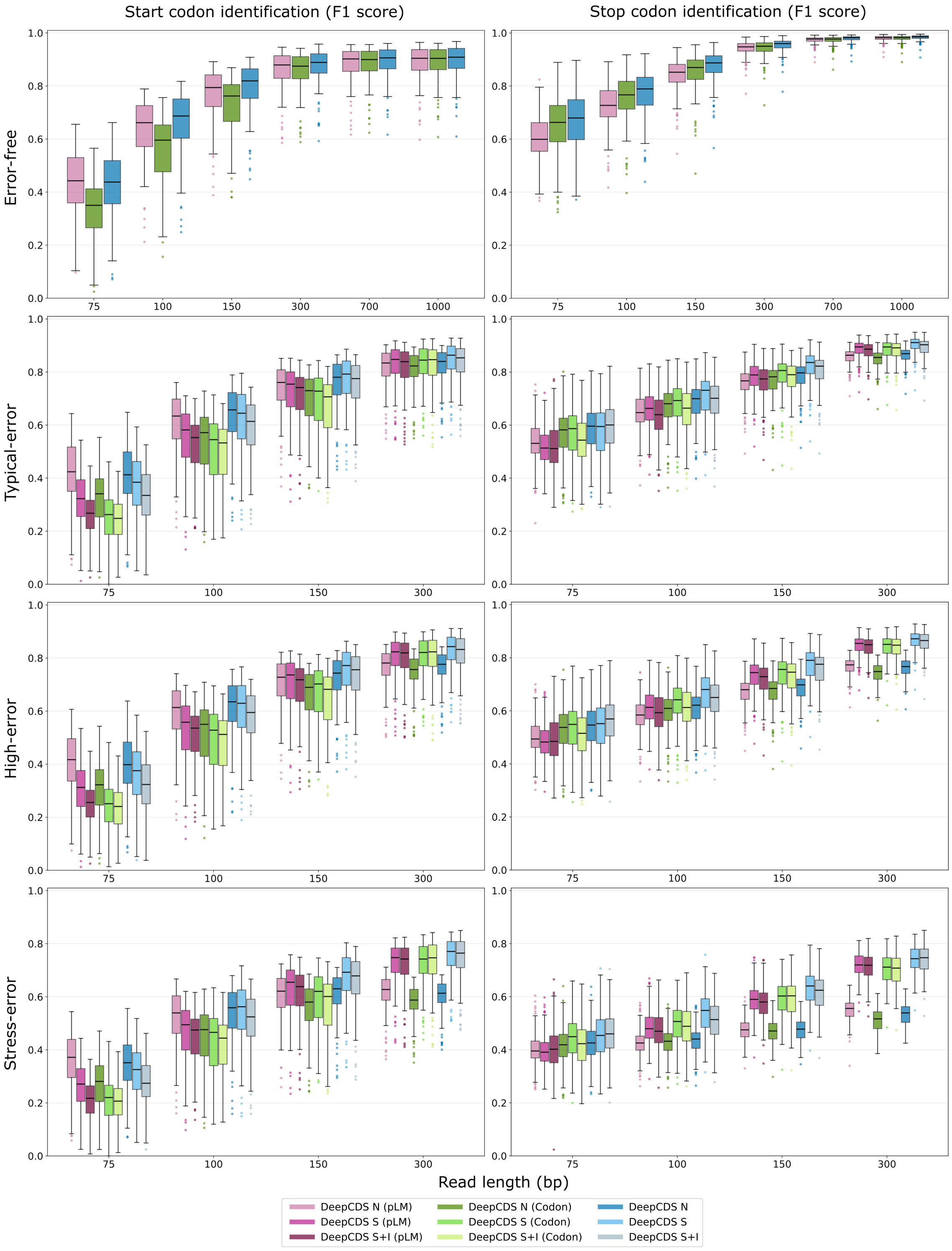


Figure A21: Boxplots visualizing the start and stop codon identification performance, measured as the F1 score, across each sequence length and error rate condition test set. The results are shown at the genome level for the 212 test genomes that follow the standard prokaryotic genetic code; each data point corresponds to the aggregated performance for one genome. The results are based on all CDS fragments longer than 60 bp. (Codon) refers to the ablation models using only the codon encoding as input, and (pLM) refers to the ablation models using only the pLM embeddings as input.

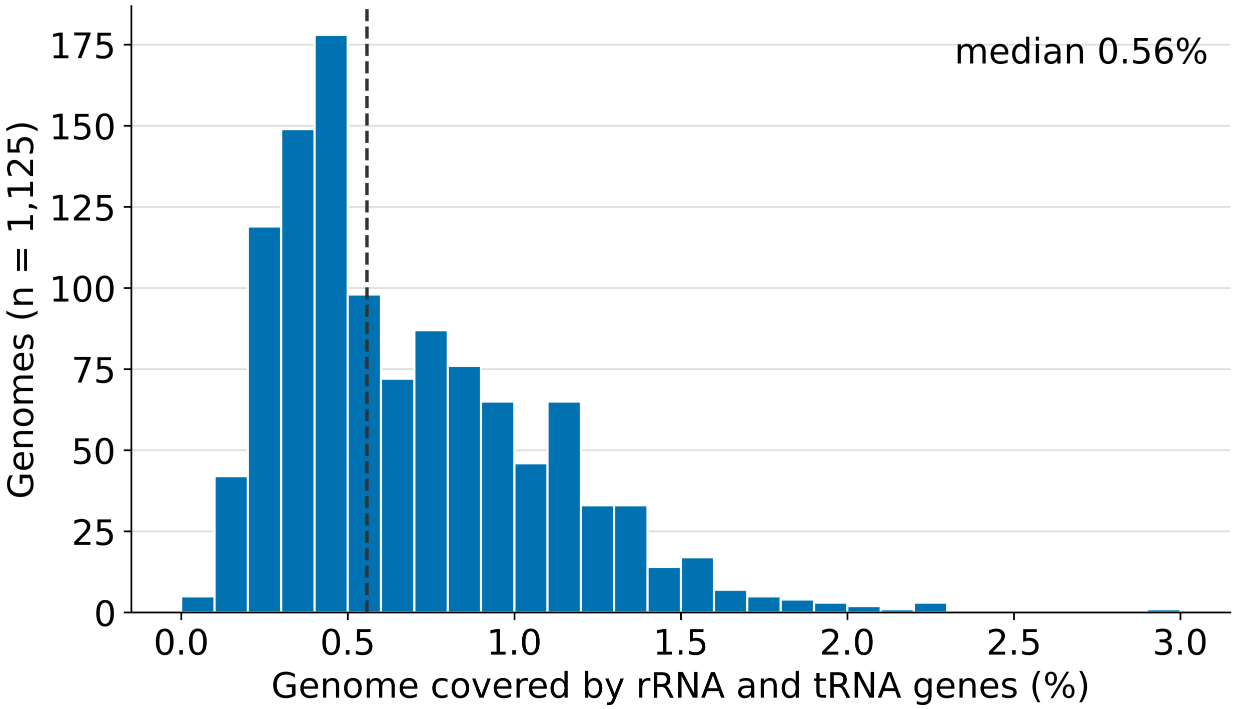


Figure A22: Share of genome positions covered by annotated rRNA and tRNA genes (positions covered by a gene on either strand, relative to the genome length) across the 1,125 genomes in the full dataset. The dashed line shows the median value.

#### A3 Supplementary Tables

Table A1: In deepcds_supplementary_tables.xlsx. Metadata for all genomes in the dataset, as retrieved from NCBI Genome Database [6].

Table A2: In deepcds_supplementary_tables.xlsx. Metadata for all genomes in the dataset: Taxonomic distribution, genomic statistics and data partition.

| **Genome** | **RefSeq accession** | **Genome Size (Mbp)** | **Genes (CDS)** | **Genome density (%)** | **GC–content (%)** | **Translation Table** | **Domain** |
|---|---|---|---|---|---|---|---|
| *A. pernix* K1 | GCF_000011125.1 | 1.67 | 1712 | 88.69 | 56.3 | 11 | Archaea |
| *A. fulgidus* DSM 4304 | GCF_000008665.1 | 2.18 | 2515 | 92.61 | 48.6 | 11 | Archaea |
| *H. salinarum* | GCF_004799605.1 | 2.43 | 2541 | 89.11 | 66.3 | 11 | Archaea |
| *M. vannielii* SB | GCF_000017165.1 | 1.72 | 1725 | 86.67 | 31.3 | 11 | Archaea |
| *M. acetivorans* C2A | GCF_000007345.1 | 5.75 | 4884 | 75.08 | 42.7 | 11 | Archaea |
| *M. stadtmanae* DSM 3091 | GCF_000012545.1 | 1.77 | 1559 | 84.42 | 27.6 | 11 | Archaea |
| *B. aphidicola* str. Sg | GCF_000007365.1 | 0.64 | 593 | 88.86 | 25.3 | 11 | Bacteria |
| *B. subtilis* str. 168 | GCF_000009045.1 | 4.21 | 4240 | 87.34 | 43.5 | 11 | Bacteria |
| *H. pylori* | GCF_025998455.1 | 1.70 | 1584 | 91.54 | 38.5 | 11 | Bacteria |
| *M. tuberculosis* H37Rv | GCF_000195955.2 | 4.41 | 3906 | 89.47 | 65.6 | 11 | Bacteria |
| *H. influenzae* | GCF_020736045.1 | 1.89 | 1808 | 88.42 | 38.2 | 11 | Bacteria |
| *M. capricolum* ATCC 27343 | GCF_000012765.1 | 1.01 | 840 | 89.97 | 23.8 | 4 | Bacteria |
| *C. jeikeium* | GCF_028609885.1 | 2.52 | 2217 | 89.31 | 61.4 | 11 | Bacteria |
| *E. coli* K-12 | GCF_000005845.2 | 4.64 | 4340 | 85.31 | 50.8 | 11 | Bacteria |
| *P. aeruginosa* PAO1 | GCF_000006765.1 | 6.26 | 5573 | 89.17 | 66.6 | 11 | Bacteria |
| *S. aureus* NCTC 8325 | GCF_000013425.1 | 2.82 | 2767 | 83.17 | 32.9 | 11 | Bacteria |

Table A3: Test set statistics based on the RefSeq annotations for the 16 representative genomes. Translation table 11 is the standard prokaryotic genetic code, and translation table 4 translates TGA as tryptophan instead of a stop codon.

| | Training set | Validation set | Test set |
|---|---|---|---|
| **Genomes (n)** | 813 (70/743) | 97 (8/89) | 215 (39/176) |
| **Families (n)** | 202 (16/186) | 31 (6/25) | 16 (6/10) |
| **Alternative genetic code (n)** | 18 | 5 | 3 |
| **GC-content range (%)** | [24.7, 74.9] | [24.4, 72.0] | [23.7, 70.5] |
| **GC-content mean (%)** | 52.0 | 51.1 | 52.0 |
| **GC-content median (%)** | 52.7 | 50.9 | 54.9 |

Table A4: Summary statistics for the complete partitioned dataset. Families (n) denotes the number of taxonomic families per partition according to NCBI's Taxonomy Database [7]. Values in parentheses for genomes and families denote (Archaea/Bacteria) counts. Alternative genetic code (n) is the number of genomes in each partition that use an alternative genetic code instead of the standard prokaryotic code, in which TGA is encoded as tryptophan rather than a stop codon (NCBI genetic code 4). All genomes with an alternative genetic code in this dataset are from the bacterial domain. The GC-content statistics are calculated per genome.

Table A5: In deepcds_supplementary_tables.xlsx. Per-genome read counts in the error-free training and validation sets (simulated with Mason [3]).

Table A6: In deepcds_supplementary_tables.xlsx. Per-genome read counts in the training and validation sets with simulated substitution errors (simulated with Mason [3]).

Table A7: In deepcds_supplementary_tables.xlsx. Per-genome read counts in the training and validation sets with simulated substitution, insertion, and deletion errors (simulated with Mason [3]).

Table A8: In deepcds_supplementary_tables.xlsx. Per-genome read counts in the error-free test sets (simulated with Mason [3]).

Table A9: In deepcds_supplementary_tables.xlsx. Per-genome read counts in the typical-error test sets (simulated with Mason [3]).

Table A10: In deepcds_supplementary_tables.xlsx. Per-genome read counts in the high-error test sets (simulated with Mason [3]).

Table A11: In deepcds_supplementary_tables.xlsx. Per-genome read counts in the stress-error test sets (simulated with Mason [3]).

Table A12: In deepcds_supplementary_tables.xlsx. Per-genome read counts in the test sets simulated with art_modern [4].

| | DeepCDS S+I | DeepCDS S | DeepCDS N |
|---|---|---|---|
| **Dropout rate (after ESM-2 encoding)** | 0 | 0 | 0 |
| **TE Layers** | 6 | 4 | 4 |
| **Attention heads (TE layers)** | 4 | 4 | 4 |
| **Dropout rate (TE layers)** | 0 | 0 | 0 |
| **Activation function (TE layers)** | relu | relu | gelu |
| **Learning rate (From scratch)** | $5.88 \cdot 10^{-6}$ | $3.30 \cdot 10^{-5}$ | $5.46 \cdot 10^{-5}$ |
| **Learning rate (fine-tuning ESM-2)** | $1.37 \cdot 10^{-7}$ | $1.77 \cdot 10^{-6}$ | $1.36 \cdot 10^{-6}$ |
| **Transition weight (Infrequent transitions)** | -3.42 | -3.28 | -1.88 |

Table A13: Selected hyperparameters based on hyperparameter optimization for each model architecture and error type configuration. TE layers = Transformer Encoder layers, specifies hyperparameter settings related to the transformer encoder layers trained from scratch.

| **Setting** | **Value** |
|---|---|
| *Reproducibility* | |
| Random seed | 42, applied to Python (`random`), NumPy, PyTorch (CPU and CUDA), and data loader workers |
| *Optimization* | |
| Optimizer | AdamW, weight decay 0.01 |
| Gradient clipping | Maximum norm 1.0 |
| Batch size | 32 |
| Maximum training length | 20 epochs |
| Learning rate scheduler | ReduceLROnPlateau on the validation loss: factor 0.7, patience 6 validations, minimum learning rate $10^{-7}$ |
| ESM-2 fine-tuning | ESM-2 frozen for the first 3 million training sequences; the last half of the ESM-2 layers was then unfrozen with a linear learning rate warmup over 32,000 training sequences, starting at 10% of the ESM-2 learning rate |
| *Validation, early stopping, and checkpoint selection* | |
| Validation frequency | Every 5,000 (DeepCDS N), 6,000 (DeepCDS S), and 10,000 (DeepCDS S+I) training steps |
| Early stopping | Training stopped after 16 consecutive validations without improvement in the validation loss (negative log-likelihood) |
| Checkpoint selection | Checkpoint with the lowest validation loss |
| *Hyperparameter optimization* | |
| Procedure | Optuna, 30 trials per model, minimizing the validation loss, with Hyperband pruning; configurations that did not converge were excluded |
| Search space | Transformer encoder depth $\{2, 4, 6, 8\}$; attention heads $\{2, 4\}$; dropout after ESM-2 $\{0, 0.1, 0.2, 0.3, 0.4\}$; dropout in transformer layers $\{0, 0.2, 0.3, 0.4, 0.5\}$; activation function {ReLU, GELU}; learning rate (weights trained from scratch) $5 \cdot 10^{-6}$ to $10^{-4}$ (log scale); ESM-2 learning rate 0.01 to 0.5 times the learning rate from scratch (log scale); initial transition score for infrequent valid transitions $[-4, -1]$ |
| Selected values | See Supplementary Table A13 |

Table A14: Settings used for training all DeepCDS model variants. Please note that the hyperparameter search was not seeded.

| Training genomes | Typical-error | High-error | Stress-error |
|---|---|---|---|
| 100 | 0.932 (0.022) | 0.919 (0.024) | 0.875 (0.030) |
| 200 | 0.936 (0.022) | 0.924 (0.024) | 0.877 (0.029) |
| 400 | 0.939 (0.021) | 0.926 (0.023) | 0.882 (0.028) |
| 813 | 0.939 (0.021) | 0.927 (0.023) | 0.884 (0.028) |

Table A15: DeepCDS training set size experiment. Aggregated F1 scores on the 300bp test sets of each error rate condition calculated for DeepCDS S+I model variants trained on smaller subsets of the training genomes, namely 100, 200, and 400, compared to the full training set of 813 genomes. The smaller subsets follow the same phylogenetic distribution, based on the family level, as the full training set. The per-genome standard deviation is stated in parentheses. Performance gains are modest when comparing 100 to 813 training genomes, with performance largely plateauing beyond 200 training genomes. This suggests that the diversity captured by a smaller number of phylogenetically representative genomes is sufficient for near-plateau performance.

| **Family** | **Genome count** | **Genome size (Mbp)** | **Genes (CDS)** | **Genome density (%)** | **GC–content (%)** | **Translation Table** | **Domain** |
|---|---|---|---|---|---|---|---|
| Archaeoglobaceae | 3 | 2.18 [1.90–2.20] | 2,515 [2,184–2,573] | 92.05 [91.99–92.61] | 46.5 [44.1–48.6] | 11 | Archaea |
| Desulfurococcaceae | 6 | 1.35 [1.30–1.67] | 1,490 [1,368–1,712] | 89.82 [88.69–92.49] | 54.7 [44.8–56.7] | 11 | Archaea |
| Halobacteriaceae | 9 | 2.71 [2.08–3.75] | 2,927 [2,149–3,760] | 89.10 [86.53–91.88] | 66.3 [56.4–68.6] | 11 | Archaea |
| Methanobacteriaceae | 8 | 1.81 [1.50–2.94] | 1,792 [1,559–2,438] | 87.64 [77.89–90.51] | 36.0 [27.6–49.6] | 11 | Archaea |
| Methanococcaceae | 2 | 1.72 [1.71–1.72] | 1,774 [1,725–1,824] | 87.70 [86.67–88.74] | 32.1 [31.3–32.9] | 11 | Archaea |
| Methanosarcinaceae | 11 | 3.13 [1.83–5.75] | 2,744 [1,610–4,884] | 76.48 [71.76–89.95] | 41.4 [39.2–43.3] | 11 | Archaea |
| Bacillaceae | 22 | 4.22 [2.70–5.75] | 4,260 [2,702–5,842] | 83.17 [76.15–88.43] | 39.8 [35.7–46.9] | 11 | Bacteria |
| Corynebacteriaceae | 33 | 2.52 [2.28–3.31] | 2,360 [2,019–3,001] | 88.50 [82.23–91.37] | 59.4 [52.2–69.3] | 11 | Bacteria |
| Enterobacteriaceae | 33 | 4.90 [4.36–6.30] | 4,555 [3,985–5,842] | 88.10 [85.31–89.80] | 54.6 [49.9–58.0] | 11 | Bacteria |
| Erwiniaceae | 6 | 4.69 [0.64–4.80] | 4,262 [593–4,391] | 86.30 [86.04–88.86] | 54.7 [25.3–57.7] | 11 | Bacteria |
| Helicobacteraceae | 2 | 1.70 [1.70–1.70] | 1,634 [1,584–1,683] | 92.47 [91.54–93.41] | 37.0 [35.4–38.5] | 11 | Bacteria |
| Mycobacteriaceae | 27 | 6.01 [3.27–8.08] | 5,476 [3,144–7,793] | 91.88 [69.77–93.24] | 66.2 [57.9–70.5] | 11 | Bacteria |
| Mycoplasmataceae | 3 | 1.01 [1.01–1.03] | 840 [829–871] | 89.97 [89.75–91.30] | 23.8 [23.7–23.8] | 4 | Bacteria |
| Pasteurellaceae | 9 | 2.14 [1.89–2.50] | 1,972 [1,808–2,260] | 88.44 [87.96–89.92] | 40.3 [38.2–44.9] | 11 | Bacteria |
| Pseudomonadaceae | 24 | 6.25 [4.04–7.33] | 5,612 [3,771–6,405] | 88.69 [87.03–90.15] | 61.5 [54.4–66.6] | 11 | Bacteria |
| Staphylococcaceae | 17 | 2.57 [2.17–2.82] | 2,429 [2,057–2,767] | 85.32 [83.17–89.87] | 33.2 [32.2–37.7] | 11 | Bacteria |

Table A16: Test set statistics aggregated to the family level based on RefSeq annotations. For each metric, the median value on a genome-level is reported. The bracketed ranges indicate the minimum and maximum observed across genomes in the family.

| | Original | | Cleaned | | |
|---|---|---|---|---|---|
| | **Full dataset** | **Model development species removed** | **Base** | **Excl. uncertain annotations** | **Excl. contigs lacking annotations** |
| **Read pairs** | 16,633,513 | 13,744,053 | 13,629,005 | 13,620,644 | 11,470,050 |
| **Individual reads** | 33,267,026 | 27,488,106 | 26,976,132 | 25,855,334 | 22,658,222 |
| Bacteria | - | - | 21,659,273 | 20,658,353 | 21,636,137 |
| Eukaryota | - | - | 4,795,966 | 4,795,966 | 501,192 |
| Viruses | - | - | 520,893 | 401,015 | 520,893 |
| **Reads with a labeled CDS** | - | - | 10,486,866 | 9,369,179 | 10,486,866 |
| **Distinct genomes represented** | 309 | 266 | 257 | 257 | 247 |
| Bacteria | - | - | 113 | 113 | 113 |
| Eukaryota | - | - | 11 | 11 | 1 |
| Viruses | - | - | 133 | 133 | 133 |

Table A17: Statistics of the simulated CAMI III sample 0 "Toy Longitudinal Human Gut" test set. "Original" refers to the statistics of the complete dataset before preprocessing annotations. "Model development species removed" refers to the dataset after removing genomes identical to, or with an ANI of at least 95% to, any genome in the training or validation sets. "Cleaned" refers to all retained sequences after additionally removing reads overlapping with CDS annotations tagged with "pseudo=true", "pseudogene", "partial=true", or "note=programmed frameshift". "Uncertain annotations" refers to all retained sequences after removing reads that also overlap with CDS annotations tagged with either "ab initio prediction" or "product=hypothetical protein". "Contigs lacking annotations" refers to contigs where no annotations were available.

#### Testing different overlap criteria

All results presented in the manuscript work are based on a complete overlap, i.e., a CDS fragment is only considered a TP if the predicted start and stop coordinates are identical to the ground truth coordinates. However, we also tested performance on less strict overlap criteria. An overlap was measured as the intersection over union (IoU), and the results with overlap 0.8 and 0.5, respectively, are shown on each test set of error rate condition and sequence length. The performance is reported as the aggregated CDS-level F1 score, sensitivity, and precision based on the 212 test genomes that follow the standard prokaryotic genetic code. Aggregated means that the TPs, FNs, FPs, and TNs are summed over the 212 genomes and then the scores are calculated. The results are based on all CDS fragments longer than 60 bp. The numbers marked in bold highlight the highest performance metric for the specific sequence length condition.

| | 75bp | 100bp | 150bp | 300bp | 700bp | 1,000bp |
|---|---|---|---|---|---|---|
| **F1 Score** | | | | | | |
| FGS (Complete) | 0.822 | 0.858 | 0.887 | 0.898 | 0.888 | 0.886 |
| MetaProdigal | 0.827 | 0.867 | 0.907 | 0.942 | 0.960 | 0.965 |
| DeepCDS N | **0.882** | **0.928** | **0.964** | **0.982** | **0.984** | **0.985** |
| **Sensitivity** | | | | | | |
| FGS (Complete) | 0.876 | 0.905 | 0.925 | 0.935 | 0.931 | 0.934 |
| MetaProdigal | 0.868 | 0.904 | 0.929 | 0.952 | 0.963 | 0.967 |
| DeepCDS N | **0.899** | **0.934** | **0.959** | **0.979** | **0.980** | **0.981** |
| **Precision** | | | | | | |
| FGS (Complete) | 0.774 | 0.816 | 0.852 | 0.865 | 0.848 | 0.843 |
| MetaProdigal | 0.789 | 0.834 | 0.887 | 0.932 | 0.957 | 0.964 |
| DeepCDS N | **0.865** | **0.921** | **0.968** | **0.984** | **0.987** | **0.989** |

Table A18: Aggregated CDS-level performance with overlap criterion = **0.8** on the **error-free** test set. Standard errors are omitted from the cells for readability; across all metrics and read lengths, the genome-level cluster bootstrap standard error spans, per model: FGS (Complete) 0.001–0.004; MetaProdigal 0.001–0.003; DeepCDS N 0.000–0.003.

| | 75bp | 100bp | 150bp | 300bp | 700bp | 1,000bp |
|---|---|---|---|---|---|---|
| **F1 Score** | | | | | | |
| FGS (Complete) | 0.822 | 0.865 | 0.900 | 0.916 | 0.911 | 0.910 |
| MetaProdigal | 0.827 | 0.874 | 0.918 | 0.952 | 0.968 | 0.973 |
| DeepCDS N | **0.882** | **0.931** | **0.968** | **0.986** | **0.988** | **0.989** |
| **Sensitivity** | | | | | | |
| FGS (Complete) | 0.876 | 0.913 | 0.939 | 0.953 | 0.956 | 0.959 |
| MetaProdigal | 0.868 | 0.911 | 0.940 | 0.962 | 0.971 | 0.974 |
| DeepCDS N | **0.899** | **0.937** | **0.964** | **0.983** | **0.984** | **0.985** |
| **Precision** | | | | | | |
| FGS (Complete) | 0.774 | 0.823 | 0.865 | 0.881 | 0.871 | 0.865 |
| MetaProdigal | 0.789 | 0.840 | 0.897 | 0.942 | 0.965 | 0.971 |
| DeepCDS N | **0.865** | **0.924** | **0.972** | **0.988** | **0.991** | **0.992** |

Table A19: Aggregated CDS-level performance with overlap criterion = **0.5** on the **error-free** test set. Standard errors are omitted from the cells for readability; across all metrics and read lengths, the genome-level cluster bootstrap standard error spans, per model: FGS (Complete) 0.001–0.004; MetaProdigal 0.001–0.003; DeepCDS N 0.000–0.003.

| | 75bp | 100bp | 150bp | 300bp |
|---|---|---|---|---|
| **F1 Score** | | | | |
| FGS (Complete) | 0.814 | 0.849 | 0.875 | 0.874 |
| FGS (0.5% error rate) | 0.807 | 0.836 | 0.852 | 0.833 |
| FGS (1.0% error rate) | 0.802 | 0.830 | 0.844 | 0.823 |
| MetaProdigal | 0.818 | 0.858 | 0.895 | 0.918 |
| DeepCDS N | 0.874 | 0.919 | 0.953 | 0.962 |
| DeepCDS S | **0.880** | **0.925** | **0.960** | **0.975** |
| DeepCDS S+I | 0.862 | 0.912 | 0.954 | 0.971 |
| **Sensitivity** | | | | |
| FGS (Complete) | 0.866 | 0.895 | 0.914 | 0.916 |
| FGS (0.5% error rate) | 0.860 | 0.885 | 0.891 | 0.861 |
| FGS (1.0% error rate) | 0.855 | 0.879 | 0.883 | 0.849 |
| MetaProdigal | 0.858 | 0.894 | 0.917 | 0.933 |
| DeepCDS N | 0.887 | 0.923 | 0.947 | 0.962 |
| DeepCDS S | 0.898 | 0.930 | 0.955 | **0.974** |
| DeepCDS S+I | **0.919** | **0.940** | **0.956** | 0.969 |
| **Precision** | | | | |
| FGS (Complete) | 0.768 | 0.807 | 0.839 | 0.835 |
| FGS (0.5% error rate) | 0.760 | 0.793 | 0.816 | 0.807 |
| FGS (1.0% error rate) | 0.755 | 0.786 | 0.808 | 0.800 |
| MetaProdigal | 0.782 | 0.825 | 0.874 | 0.904 |
| DeepCDS N | 0.861 | 0.916 | 0.960 | 0.963 |
| DeepCDS S | **0.864** | **0.921** | **0.966** | **0.977** |
| DeepCDS S+I | 0.811 | 0.885 | 0.952 | 0.973 |

Table A20: Aggregated CDS-level performance with overlap criterion = **0.8** on the **typical-error** test set. Standard errors are omitted from the cells for readability; across all metrics and read lengths, the genome-level cluster bootstrap standard error spans, per model: FGS (Complete) 0.001–0.004; FGS (0.5% error rate) and FGS (1.0% error rate): 0.002–0.004; MetaProdigal, DeepCDS N, DeepCDS S and DeepCDS S+I 0.001–0.003.

| | 75bp | 100bp | 150bp | 300bp |
|---|---|---|---|---|
| **F1 Score** | | | | |
| FGS (Complete) | 0.814 | 0.858 | 0.893 | 0.903 |
| FGS (0.5% error rate) | 0.810 | 0.852 | 0.885 | 0.895 |
| FGS (1.0% error rate) | 0.807 | 0.849 | 0.882 | 0.892 |
| MetaProdigal | 0.819 | 0.866 | 0.910 | 0.940 |
| DeepCDS N | 0.874 | 0.924 | 0.963 | 0.977 |
| DeepCDS S | **0.881** | **0.930** | **0.967** | **0.982** |
| DeepCDS S+I | 0.863 | 0.918 | 0.962 | 0.980 |
| **Sensitivity** | | | | |
| FGS (Complete) | 0.866 | 0.905 | 0.933 | 0.947 |
| FGS (0.5% error rate) | 0.863 | 0.901 | 0.926 | 0.925 |
| FGS (1.0% error rate) | 0.861 | 0.899 | 0.923 | 0.919 |
| MetaProdigal | 0.858 | 0.902 | 0.933 | 0.955 |
| DeepCDS N | 0.887 | 0.928 | 0.956 | 0.976 |
| DeepCDS S | 0.898 | 0.935 | 0.961 | **0.980** |
| DeepCDS S+I | **0.920** | **0.946** | **0.964** | 0.978 |
| **Precision** | | | | |
| FGS (Complete) | 0.768 | 0.816 | 0.856 | 0.863 |
| FGS (0.5% error rate) | 0.763 | 0.807 | 0.848 | 0.866 |
| FGS (1.0% error rate) | 0.760 | 0.804 | 0.845 | 0.866 |
| MetaProdigal | 0.783 | 0.833 | 0.889 | 0.926 |
| DeepCDS N | 0.861 | 0.921 | 0.969 | 0.977 |
| DeepCDS S | **0.864** | **0.925** | **0.972** | **0.983** |
| DeepCDS S+I | 0.812 | 0.891 | 0.960 | 0.981 |

Table A21: Aggregated CDS-level performance with overlap criterion = **0.5** on the **typical-error** test set. Standard errors are omitted from the cells for readability; across all metrics and read lengths, the genome-level cluster bootstrap standard error spans, per model: FGS (Complete) 0.001–0.004; FGS (0.5% error rate) and FGS (1.0% error rate): 0.002–0.004; MetaProdigal and DeepCDS S+I 0.001–0.003; DeepCDS N and DeepCDS S 0.000–0.003.

| | 75bp | 100bp | 150bp | 300bp |
|---|---|---|---|---|
| **F1 Score** | | | | |
| FGS (Complete) | 0.800 | 0.832 | 0.848 | 0.821 |
| FGS (0.5% error rate) | 0.793 | 0.820 | 0.828 | 0.793 |
| FGS (1.0% error rate) | 0.789 | 0.814 | 0.820 | 0.787 |
| MetaProdigal | 0.805 | 0.841 | 0.870 | 0.867 |
| DeepCDS N | 0.860 | 0.903 | 0.930 | 0.921 |
| DeepCDS S | **0.870** | **0.916** | **0.952** | **0.968** |
| DeepCDS S+I | 0.852 | 0.903 | 0.946 | 0.964 |
| **Sensitivity** | | | | |
| FGS (Complete) | 0.850 | 0.876 | 0.887 | 0.871 |
| FGS (0.5% error rate) | 0.844 | 0.867 | 0.867 | 0.827 |
| FGS (1.0% error rate) | 0.840 | 0.861 | 0.859 | 0.817 |
| MetaProdigal | 0.842 | 0.874 | 0.891 | 0.887 |
| DeepCDS N | 0.868 | 0.900 | 0.918 | 0.922 |
| DeepCDS S | 0.883 | 0.917 | 0.943 | **0.965** |
| DeepCDS S+I | **0.907** | **0.929** | **0.945** | 0.962 |
| **Precision** | | | | |
| FGS (Complete) | 0.757 | 0.792 | 0.813 | 0.776 |
| FGS (0.5% error rate) | 0.749 | 0.778 | 0.792 | 0.762 |
| FGS (1.0% error rate) | 0.744 | 0.772 | 0.785 | 0.759 |
| MetaProdigal | 0.771 | 0.811 | 0.850 | 0.848 |
| DeepCDS N | 0.853 | 0.906 | 0.942 | 0.920 |
| DeepCDS S | **0.858** | **0.915** | **0.960** | **0.970** |
| DeepCDS S+I | 0.804 | 0.878 | 0.946 | 0.967 |

Table A22: Aggregated CDS-level performance with overlap criterion = **0.8** on the **high-error** test set. Standard errors are omitted from the cells for readability; across all metrics and read lengths, the genome-level cluster bootstrap standard error spans, per model: FGS (Complete) 0.001–0.004; FGS (0.5% error rate) and FGS (1.0% error rate): 0.002–0.004; MetaProdigal, DeepCDS N, DeepCDS S and DeepCDS S+I 0.001–0.003.

| | 75bp | 100bp | 150bp | 300bp |
|---|---|---|---|---|
| **F1 Score** | | | | |
| FGS (Complete) | 0.801 | 0.844 | 0.879 | 0.878 |
| FGS (0.5% error rate) | 0.797 | 0.839 | 0.872 | 0.877 |
| FGS (1.0% error rate) | 0.794 | 0.836 | 0.870 | 0.876 |
| MetaProdigal | 0.806 | 0.852 | 0.895 | 0.915 |
| DeepCDS N | 0.861 | 0.912 | 0.952 | 0.960 |
| DeepCDS S | **0.871** | **0.922** | **0.961** | **0.978** |
| DeepCDS S+I | 0.853 | 0.910 | 0.957 | 0.977 |
| **Sensitivity** | | | | |
| FGS (Complete) | 0.851 | 0.889 | 0.919 | 0.931 |
| FGS (0.5% error rate) | 0.848 | 0.886 | 0.914 | 0.914 |
| FGS (1.0% error rate) | 0.845 | 0.884 | 0.911 | 0.910 |
| MetaProdigal | 0.843 | 0.885 | 0.916 | 0.936 |
| DeepCDS N | 0.868 | 0.910 | 0.940 | 0.962 |
| DeepCDS S | 0.884 | 0.923 | 0.952 | **0.976** |
| DeepCDS S+I | **0.908** | **0.936** | **0.957** | 0.974 |
| **Precision** | | | | |
| FGS (Complete) | 0.757 | 0.804 | 0.842 | 0.830 |
| FGS (0.5% error rate) | 0.752 | 0.796 | 0.835 | 0.842 |
| FGS (1.0% error rate) | 0.749 | 0.792 | 0.832 | 0.845 |
| MetaProdigal | 0.772 | 0.821 | 0.874 | 0.895 |
| DeepCDS N | 0.853 | 0.915 | 0.964 | 0.959 |
| DeepCDS S | **0.858** | **0.921** | **0.970** | **0.981** |
| DeepCDS S+I | 0.805 | 0.885 | 0.957 | 0.980 |

Table A23: Aggregated CDS-level performance with overlap criterion = **0.5** on the **high-error** test set. Standard errors are omitted from the cells for readability; across all metrics and read lengths, the genome-level cluster bootstrap standard error spans, per model: FGS (Complete) 0.001–0.004; FGS (0.5% error rate) and FGS (1.0% error rate): 0.002–0.004; MetaProdigal, DeepCDS N, DeepCDS S and DeepCDS S+I 0.001–0.003.

| | 75bp | 100bp | 150bp | 300bp |
|---|---|---|---|---|
| **F1 Score** | | | | |
| FGS (Complete) | 0.757 | 0.780 | 0.768 | 0.675 |
| FGS (0.5% error rate) | 0.750 | 0.768 | 0.753 | 0.675 |
| FGS (1.0% error rate) | 0.746 | 0.762 | 0.748 | 0.677 |
| MetaProdigal | 0.761 | 0.789 | 0.793 | 0.723 |
| DeepCDS N | 0.816 | 0.851 | 0.855 | 0.798 |
| DeepCDS S | **0.836** | **0.884** | **0.919** | **0.939** |
| DeepCDS S+I | 0.819 | 0.870 | 0.913 | 0.937 |
| **Sensitivity** | | | | |
| FGS (Complete) | 0.799 | 0.817 | 0.806 | 0.740 |
| FGS (0.5% error rate) | 0.794 | 0.809 | 0.791 | 0.722 |
| FGS (1.0% error rate) | 0.789 | 0.804 | 0.786 | 0.719 |
| MetaProdigal | 0.790 | 0.815 | 0.809 | 0.753 |
| DeepCDS N | 0.805 | 0.829 | 0.827 | 0.799 |
| DeepCDS S | 0.835 | 0.872 | 0.898 | **0.932** |
| DeepCDS S+I | **0.863** | **0.886** | **0.903** | 0.928 |
| **Precision** | | | | |
| FGS (Complete) | 0.720 | 0.745 | 0.734 | 0.620 |
| FGS (0.5% error rate) | 0.712 | 0.731 | 0.718 | 0.634 |
| FGS (1.0% error rate) | 0.707 | 0.725 | 0.714 | 0.639 |
| MetaProdigal | 0.734 | 0.765 | 0.777 | 0.695 |
| DeepCDS N | 0.827 | 0.874 | 0.886 | 0.797 |
| DeepCDS S | **0.836** | **0.896** | **0.941** | **0.947** |
| DeepCDS S+I | 0.779 | 0.854 | 0.923 | 0.946 |

Table A24: Aggregated CDS-level performance with overlap criterion = **0.8** on the **stress-error** test set. Standard errors are omitted from the cells for readability; across all metrics and read lengths, the genome-level cluster bootstrap standard error spans, per model: FGS (Complete) and FGS (0.5% error rate) 0.002–0.004; FGS (1.0% error rate) 0.003–0.004; MetaProdigal 0.002–0.003; DeepCDS N and DeepCDS S+I 0.001–0.003; DeepCDS S 0.001–0.004.

| | 75bp | 100bp | 150bp | 300bp |
|---|---|---|---|---|
| **F1 Score** | | | | |
| FGS (Complete) | 0.759 | 0.801 | 0.833 | 0.802 |
| FGS (0.5% error rate) | 0.755 | 0.797 | 0.830 | 0.819 |
| FGS (1.0% error rate) | 0.753 | 0.794 | 0.829 | 0.822 |
| MetaProdigal | 0.763 | 0.806 | 0.845 | 0.841 |
| DeepCDS N | 0.817 | 0.872 | 0.914 | 0.906 |
| DeepCDS S | **0.837** | **0.894** | **0.940** | **0.965** |
| DeepCDS S+I | 0.821 | 0.883 | 0.936 | 0.964 |
| **Sensitivity** | | | | |
| FGS (Complete) | 0.801 | 0.840 | 0.875 | 0.880 |
| FGS (0.5% error rate) | 0.799 | 0.839 | 0.873 | 0.876 |
| FGS (1.0% error rate) | 0.796 | 0.837 | 0.871 | 0.873 |
| MetaProdigal | 0.792 | 0.832 | 0.863 | 0.876 |
| DeepCDS N | 0.806 | 0.849 | 0.883 | 0.907 |
| DeepCDS S | 0.837 | 0.882 | 0.918 | **0.957** |
| DeepCDS S+I | **0.865** | **0.899** | **0.926** | 0.955 |
| **Precision** | | | | |
| FGS (Complete) | 0.721 | 0.766 | 0.796 | 0.737 |
| FGS (0.5% error rate) | 0.716 | 0.758 | 0.792 | 0.769 |
| FGS (1.0% error rate) | 0.713 | 0.755 | 0.791 | 0.777 |
| MetaProdigal | 0.736 | 0.781 | 0.828 | 0.809 |
| DeepCDS N | 0.828 | 0.895 | 0.946 | 0.905 |
| DeepCDS S | **0.838** | **0.906** | **0.962** | 0.972 |
| DeepCDS S+I | 0.781 | 0.867 | 0.946 | **0.973** |

Table A25: Aggregated CDS-level performance with overlap criterion = **0.5** on the **stress-error** test set. Standard errors are omitted from the cells for readability; across all metrics and read lengths, the genome-level cluster bootstrap standard error spans, per model: FGS (Complete) and DeepCDS S 0.001–0.004; FGS (0.5% error rate) 0.002–0.004; FGS (1.0% error rate) 0.002–0.005; MetaProdigal, DeepCDS N, and DeepCDS S+I 0.001–0.003.

| | 75bp | 100bp | 150bp | 300bp | 700bp | 1,000bp |
|---|---|---|---|---|---|---|
| **ATG (canonical) start codons** | | | | | | |
| True start codon occurrences | 86 033 | 187 044 | 285 304 | 376 859 | 419 835 | 423 591 |
| **F1 score** | | | | | | |
| MetaProdigal | 0.143 ± 0.002 | 0.471 ± 0.005 | 0.633 ± 0.005 | 0.789 ± 0.004 | 0.867 ± 0.003 | 0.880 ± 0.003 |
| DeepCDS N | **0.471** ± 0.006 | **0.710** ± 0.005 | **0.837** ± 0.004 | **0.909** ± 0.003 | **0.926** ± 0.002 | **0.930** ± 0.002 |
| **Precision** | | | | | | |
| MetaProdigal | 0.415 ± 0.008 | 0.536 ± 0.008 | 0.649 ± 0.007 | 0.792 ± 0.005 | 0.869 ± 0.004 | 0.881 ± 0.004 |
| DeepCDS N | **0.762** ± 0.006 | **0.836** ± 0.004 | **0.880** ± 0.003 | **0.912** ± 0.003 | **0.923** ± 0.003 | **0.927** ± 0.003 |
| **Sensitivity** | | | | | | |
| MetaProdigal | 0.087 ± 0.001 | 0.420 ± 0.004 | 0.618 ± 0.005 | 0.787 ± 0.003 | 0.864 ± 0.003 | 0.880 ± 0.003 |
| DeepCDS N | **0.340** ± 0.006 | **0.617** ± 0.005 | **0.797** ± 0.004 | **0.906** ± 0.003 | **0.929** ± 0.002 | **0.933** ± 0.002 |
| **GTG start codons** | | | | | | |
| True start codon occurrences | 13 332 | 29 077 | 44 289 | 58 150 | 64 135 | 64 731 |
| **F1 score** | | | | | | |
| MetaProdigal | 0.037 ± 0.002 | 0.245 ± 0.005 | 0.401 ± 0.005 | 0.590 ± 0.006 | 0.708 ± 0.006 | 0.728 ± 0.007 |
| DeepCDS N | **0.155** ± 0.006 | **0.459** ± 0.008 | **0.657** ± 0.008 | **0.778** ± 0.006 | **0.811** ± 0.006 | **0.817** ± 0.005 |
| **Precision** | | | | | | |
| MetaProdigal | 0.292 ± 0.017 | 0.389 ± 0.008 | 0.495 ± 0.009 | 0.655 ± 0.007 | 0.754 ± 0.006 | 0.762 ± 0.006 |
| DeepCDS N | **0.739** ± 0.013 | **0.789** ± 0.006 | **0.820** ± 0.005 | **0.845** ± 0.005 | **0.847** ± 0.005 | **0.848** ± 0.005 |
| **Sensitivity** | | | | | | |
| MetaProdigal | 0.020 ± 0.001 | 0.178 ± 0.006 | 0.337 ± 0.008 | 0.537 ± 0.008 | 0.667 ± 0.008 | 0.697 ± 0.009 |
| DeepCDS N | **0.087** ± 0.004 | **0.323** ± 0.008 | **0.547** ± 0.010 | **0.721** ± 0.008 | **0.777** ± 0.007 | **0.789** ± 0.007 |
| **TTG start codons** | | | | | | |
| True start codon occurrences | 4 385 | 9 634 | 14 658 | 19 217 | 21 236 | 21 087 |
| **F1 score** | | | | | | |
| MetaProdigal | 0.016 ± 0.003 | 0.144 ± 0.011 | 0.282 ± 0.014 | 0.494 ± 0.014 | 0.627 ± 0.011 | 0.651 ± 0.011 |
| DeepCDS N | **0.082** ± 0.006 | **0.393** ± 0.013 | **0.601** ± 0.013 | **0.738** ± 0.012 | **0.769** ± 0.011 | **0.777** ± 0.011 |
| **Precision** | | | | | | |
| MetaProdigal | 0.044 ± 0.010 | 0.142 ± 0.015 | 0.281 ± 0.020 | 0.524 ± 0.019 | 0.696 ± 0.014 | 0.719 ± 0.012 |
| DeepCDS N | **0.688** ± 0.028 | **0.768** ± 0.010 | **0.801** ± 0.007 | **0.832** ± 0.007 | **0.836** ± 0.007 | **0.838** ± 0.007 |
| **Sensitivity** | | | | | | |
| MetaProdigal | 0.010 ± 0.002 | 0.147 ± 0.010 | 0.283 ± 0.012 | 0.468 ± 0.013 | 0.571 ± 0.013 | 0.595 ± 0.014 |
| DeepCDS N | **0.044** ± 0.004 | **0.264** ± 0.012 | **0.481** ± 0.016 | **0.662** ± 0.017 | **0.712** ± 0.017 | **0.725** ± 0.016 |
| **Other non-canonical start codons** | | | | | | |
| True start codon occurrences | 988 | 2 113 | 3 210 | 4 101 | 4 482 | 4 559 |
| **F1 score** | | | | | | |
| MetaProdigal | **0.000** ± 0.000 | 0.000 ± 0.000 | 0.000 ± 0.000 | 0.000 ± 0.000 | 0.000 ± 0.000 | 0.000 ± 0.000 |
| DeepCDS N | **0.000** ± 0.000 | **0.006** ± 0.002 | **0.032** ± 0.005 | **0.091** ± 0.008 | **0.120** ± 0.008 | **0.126** ± 0.009 |
| **Precision** | | | | | | |
| MetaProdigal | **0.000** ± 0.000 | 0.000 ± 0.000 | 0.000 ± 0.000 | 0.000 ± 0.000 | 0.000 ± 0.000 | 0.000 ± 0.000 |
| DeepCDS N | **0.000** ± 0.000 | **0.462** ± 0.145 | **0.417** ± 0.053 | **0.349** ± 0.023 | **0.312** ± 0.018 | **0.311** ± 0.017 |
| **Sensitivity** | | | | | | |
| MetaProdigal | **0.000** ± 0.000 | 0.000 ± 0.000 | 0.000 ± 0.000 | 0.000 ± 0.000 | 0.000 ± 0.000 | 0.000 ± 0.000 |
| DeepCDS N | **0.000** ± 0.000 | **0.003** ± 0.001 | **0.017** ± 0.002 | **0.052** ± 0.005 | **0.075** ± 0.006 | **0.079** ± 0.006 |

Table A26: Start codon prediction performance calculated separately on canonical (ATG) versus groups of non-canonical (GTG, TTG, and others) start codons on the error-free test sets simulated at different sequence lengths, including start codons at sequence borders. The scores are based on summed counts of TPs, FPs, and FNs, aggregated across the 212 test genomes that follow the standard prokaryotic genetic code. The standard error from a genome-level cluster bootstrap is reported (±). For each test set, the best performance is marked in bold. Because FragGeneScan does not explicitly annotate start codon positions, we can not assess whether border positions represent a true start codon position or an internal part of a CDS region. For this reason, FragGeneScan is omitted from these results. It should be noted that the number of occurrences of true start codons increases with test set sequence length; this is due to the fact that many CDS parts in the test sets become shorter than 61 bp and for that reason are excluded from the results, as FragGeneScan and MetaProdigal do not predict shorter stretches of CDSs.

| | 75bp | 100bp | 150bp | 300bp | 700bp | 1,000bp |
|---|---|---|---|---|---|---|
| **MCC** | | | | | | |
| FGS (Complete) | 0.800 | 0.849 | 0.898 | 0.942 | 0.959 | 0.962 |
| MetaProdigal | 0.805 | 0.858 | 0.914 | 0.965 | 0.984 | 0.987 |
| DeepCDS N | **0.866** | **0.922** | **0.967** | **0.990** | **0.994** | **0.995** |
| **F1 Score** | | | | | | |
| FGS (Complete) | 0.813 | 0.841 | 0.863 | 0.855 | 0.796 | 0.766 |
| MetaProdigal | 0.819 | 0.855 | 0.889 | 0.913 | 0.910 | 0.906 |
| DeepCDS N | **0.876** | **0.919** | **0.953** | **0.965** | **0.952** | **0.946** |
| **Sensitivity** | | | | | | |
| FGS (Complete) | 0.866 | 0.887 | 0.900 | 0.889 | 0.835 | 0.807 |
| MetaProdigal | 0.861 | 0.891 | 0.911 | 0.922 | 0.913 | 0.907 |
| DeepCDS N | **0.894** | **0.926** | **0.949** | **0.962** | **0.949** | **0.942** |
| **Precision** | | | | | | |
| FGS (Complete) | 0.766 | 0.800 | 0.829 | 0.822 | 0.761 | 0.728 |
| MetaProdigal | 0.782 | 0.821 | 0.869 | 0.903 | 0.908 | 0.904 |
| DeepCDS N | **0.860** | **0.913** | **0.958** | **0.967** | **0.955** | **0.949** |

Table A27: Aggregated CDS-level (F1 score, Sensitivity, and Precision) and codon-level (MCC) performance metrics on the error-free test sets of different sequence lengths, based on the 212 test genomes that follow the standard prokaryotic genetic code. Aggregated means that the TPs, FNs, FPs, and TNs are summed over all the genomes, from which the scores are calculated. The FGS and DeepCDS models are those trained on data without sequencing errors. The results are based on all CDS fragments longer than 60 bp. The numbers marked in bold highlight the highest performance metric for the specific sequence length condition. Standard errors are omitted from the cells for readability; across all metrics and read lengths, the genome-level cluster bootstrap standard error spans, per model: FGS (Complete) 0.001–0.004; MetaProdigal 0.001–0.003; DeepCDS N 0.000–0.003.

| | FGS (Complete) | FGS (0.5% error rate) | FGS (1% error rate) | MetaProdigal | DeepCDS N | DeepCDS S | DeepCDS S+I |
|---|---|---|---|---|---|---|---|
| **CDS-level F1 score** | | | | | | | |
| *A. pernix* (◇) | 0.609 | 0.544 | 0.520 | 0.879 | 0.875 | **0.892** | 0.878 |
| *A. fulgidus* (◇) | 0.712 | 0.642 | 0.618 | 0.854 | 0.870 | **0.890** | 0.877 |
| *H. salinarum* (◇) | 0.836 | 0.791 | 0.774 | 0.874 | 0.931 | **0.949** | 0.944 |
| *M. vannielii* (◇) | 0.827 | 0.779 | 0.764 | 0.860 | 0.913 | **0.939** | 0.935 |
| *M. acetivorans* (◇) | 0.739 | 0.691 | 0.675 | 0.802 | 0.862 | **0.875** | 0.865 |
| *M. stadtmanae* (◇) | 0.830 | 0.797 | 0.785 | 0.879 | 0.917 | **0.937** | 0.933 |
| *B. aphidicola* | 0.841 | 0.823 | 0.814 | 0.862 | 0.887 | **0.926** | 0.925 |
| *B. subtilis* | 0.792 | 0.761 | 0.750 | 0.842 | 0.921 | **0.947** | 0.942 |
| *H. pylori* | 0.829 | 0.790 | 0.778 | 0.849 | 0.900 | **0.928** | 0.923 |
| *M. tuberculosis* | 0.778 | 0.702 | 0.675 | 0.825 | 0.885 | **0.902** | 0.888 |
| *H. influenzae* | 0.840 | 0.806 | 0.795 | 0.883 | 0.925 | **0.950** | 0.945 |
| *M. capricolum* (∗) | 0.429 | 0.411 | 0.407 | 0.873 | 0.913 | **0.933** | 0.928 |
| *C. jeikeium* | 0.833 | 0.802 | 0.791 | 0.873 | 0.930 | **0.946** | 0.945 |
| *E. coli* | 0.792 | 0.755 | 0.744 | 0.866 | 0.914 | **0.936** | 0.929 |
| *P. aeruginosa* | 0.856 | 0.820 | 0.811 | 0.894 | 0.942 | **0.964** | 0.962 |
| *S. aureus* | 0.815 | 0.776 | 0.767 | 0.865 | 0.915 | **0.941** | 0.935 |
| **Aggregated** | 0.785 | 0.741 | 0.727 | 0.857 | 0.906 | **0.927** | 0.920 |
| **CDS-level Sensitivity** | | | | | | | |
| *A. pernix* (◇) | 0.624 | 0.550 | 0.523 | **0.890** | 0.865 | 0.882 | 0.864 |
| *A. fulgidus* (◇) | 0.734 | 0.643 | 0.615 | 0.857 | 0.864 | **0.882** | 0.870 |
| *H. salinarum* (◇) | 0.861 | 0.807 | 0.788 | 0.886 | 0.928 | **0.946** | 0.940 |
| *M. vannielii* (◇) | 0.844 | 0.788 | 0.772 | 0.867 | 0.913 | **0.941** | 0.938 |
| *M. acetivorans* (◇) | 0.819 | 0.756 | 0.737 | 0.817 | 0.845 | **0.859** | 0.850 |
| *M. stadtmanae* (◇) | 0.844 | 0.804 | 0.789 | 0.888 | 0.914 | **0.933** | 0.931 |
| *B. aphidicola* | 0.851 | 0.829 | 0.819 | 0.866 | 0.878 | 0.916 | **0.919** |
| *B. subtilis* | 0.853 | 0.805 | 0.789 | 0.858 | 0.920 | **0.945** | 0.941 |
| *H. pylori* | 0.849 | 0.799 | 0.785 | 0.845 | 0.893 | **0.921** | 0.918 |
| *M. tuberculosis* | 0.797 | 0.711 | 0.682 | 0.834 | 0.873 | **0.888** | 0.876 |
| *H. influenzae* | 0.869 | 0.826 | 0.813 | 0.887 | 0.928 | **0.952** | 0.946 |
| *M. capricolum* (∗) | 0.449 | 0.430 | 0.423 | 0.885 | 0.912 | **0.932** | 0.928 |
| *C. jeikeium* | 0.867 | 0.828 | 0.815 | 0.890 | 0.927 | **0.944** | 0.941 |
| *E. coli* | 0.847 | 0.795 | 0.780 | 0.880 | 0.921 | **0.942** | 0.935 |
| *P. aeruginosa* | 0.892 | 0.844 | 0.833 | 0.921 | 0.950 | **0.971** | 0.970 |
| *S. aureus* | 0.850 | 0.803 | 0.792 | 0.884 | 0.916 | **0.943** | 0.936 |
| **Aggregated** | 0.823 | 0.767 | 0.750 | 0.870 | 0.903 | **0.924** | 0.917 |
| **CDS-level Precision** | | | | | | | |
| *A. pernix* (◇) | 0.595 | 0.538 | 0.516 | 0.868 | 0.885 | **0.903** | 0.892 |
| *A. fulgidus* (◇) | 0.691 | 0.641 | 0.620 | 0.852 | 0.877 | **0.897** | 0.885 |
| *H. salinarum* (◇) | 0.813 | 0.776 | 0.760 | 0.862 | 0.934 | **0.952** | 0.949 |
| *M. vannielii* (◇) | 0.811 | 0.769 | 0.757 | 0.853 | 0.914 | **0.937** | 0.933 |
| *M. acetivorans* (◇) | 0.673 | 0.636 | 0.623 | 0.787 | 0.881 | **0.892** | 0.881 |
| *M. stadtmanae* (◇) | 0.817 | 0.791 | 0.781 | 0.870 | 0.920 | **0.940** | 0.935 |
| *B. aphidicola* | 0.830 | 0.818 | 0.810 | 0.858 | 0.895 | **0.936** | 0.931 |
| *B. subtilis* | 0.740 | 0.722 | 0.714 | 0.826 | 0.922 | **0.949** | 0.942 |
| *H. pylori* | 0.810 | 0.781 | 0.771 | 0.854 | 0.906 | **0.934** | 0.929 |
| *M. tuberculosis* | 0.759 | 0.692 | 0.668 | 0.815 | 0.897 | **0.916** | 0.902 |
| *H. influenzae* | 0.813 | 0.787 | 0.778 | 0.878 | 0.922 | **0.947** | 0.943 |
| *M. capricolum* (∗) | 0.410 | 0.394 | 0.393 | 0.862 | 0.914 | **0.933** | 0.928 |
| *C. jeikeium* | 0.802 | 0.777 | 0.768 | 0.857 | 0.932 | 0.949 | **0.950** |
| *E. coli* | 0.744 | 0.719 | 0.711 | 0.852 | 0.907 | **0.930** | 0.923 |
| *P. aeruginosa* | 0.822 | 0.798 | 0.790 | 0.868 | 0.933 | **0.956** | 0.955 |
| *S. aureus* | 0.782 | 0.751 | 0.743 | 0.846 | 0.914 | **0.939** | 0.934 |
| **Aggregated** | 0.751 | 0.717 | 0.705 | 0.845 | 0.909 | **0.930** | 0.924 |
| **Codon-level MCC** | | | | | | | |
| *A. pernix* (◇) | 0.759 | 0.722 | 0.707 | 0.971 | 0.976 | **0.979** | 0.976 |
| *A. fulgidus* (◇) | 0.839 | 0.821 | 0.813 | 0.968 | 0.973 | **0.978** | 0.976 |
| *H. salinarum* (◇) | 0.942 | 0.934 | 0.931 | 0.962 | 0.985 | **0.989** | **0.989** |
| *M. vannielii* (◇) | 0.954 | 0.945 | 0.941 | 0.964 | 0.984 | **0.991** | 0.989 |
| *M. acetivorans* (◇) | 0.874 | 0.852 | 0.845 | 0.918 | 0.958 | **0.960** | 0.958 |
| *M. stadtmanae* (◇) | 0.947 | 0.945 | 0.942 | 0.969 | 0.985 | 0.990 | **0.991** |
| *B. aphidicola* | 0.964 | 0.965 | 0.964 | 0.966 | 0.979 | **0.987** | **0.987** |
| *B. subtilis* | 0.936 | 0.929 | 0.926 | 0.949 | 0.986 | **0.991** | **0.991** |
| *H. pylori* | 0.959 | 0.956 | 0.952 | 0.962 | 0.981 | **0.988** | 0.987 |
| *M. tuberculosis* | 0.911 | 0.892 | 0.886 | 0.924 | 0.966 | **0.970** | 0.966 |
| *H. influenzae* | 0.961 | 0.956 | 0.952 | 0.971 | 0.988 | **0.993** | 0.992 |
| *M. capricolum* (∗) | 0.804 | 0.869 | 0.874 | 0.966 | 0.988 | **0.991** | 0.990 |
| *C. jeikeium* | 0.934 | 0.928 | 0.925 | 0.950 | 0.985 | **0.988** | 0.987 |
| *E. coli* | 0.922 | 0.914 | 0.911 | 0.947 | 0.972 | **0.976** | 0.974 |
| *P. aeruginosa* | 0.952 | 0.951 | 0.950 | 0.960 | 0.987 | **0.990** | **0.990** |
| *S. aureus* | 0.953 | 0.949 | 0.945 | 0.962 | 0.985 | **0.989** | 0.988 |
| **Aggregated** | 0.916 | 0.907 | 0.903 | 0.951 | 0.978 | **0.981** | 0.980 |

Table A28: CDS-level (F1 Score, Sensitivity, and Precision) and codon-level (MCC) performance metrics per organism on the 300bp typical-error test set, evaluated on 16 genomes commonly used in prokaryotic CDS prediction benchmarks (one genome per family in the test set). Bold indicates best-performing model per genome and metric. The aggregated score corresponds to the value obtained summing the counts of TPs, FPs, and FNs over these 16 organisms. The (◇) marks archaeal organisms, and the (∗) marks usage of an alternative genetic code (NCBI genetic code 4). The numbers marked in bold highlight the model achieving the highest performance per genome and metric. The corresponding CDS-level F1 scores aggregated to the family level are shown in Supplementary Figure A16.

| Model | Annotated start position predicted | | | Ratio |
|---|---|---|---|---|
| | Longest | Shortest | Neither | longest/shortest |
| **Error-free (26,113 reads)** | | | | |
| FGS (Complete) | 36.0% | 25.1% | 38.9% | 1.43 ± 0.07 |
| MetaProdigal | 40.9% | 24.5% | 34.6% | 1.67 ± 0.09 |
| DeepCDS N | 49.1% | 32.9% | 18.0% | 1.49 ± 0.09 |
| **Typical-error (25,971 reads)** | | | | |
| FGS (Complete) | 35.3% | 24.7% | 40.0% | 1.43 ± 0.07 |
| FGS (0.5% error rate) | 29.5% | 18.1% | 52.4% | 1.63 ± 0.09 |
| FGS (1.0% error rate) | 27.7% | 16.9% | 55.4% | 1.64 ± 0.09 |
| MetaProdigal | 39.9% | 23.9% | 36.1% | 1.67 ± 0.09 |
| DeepCDS N | 47.6% | 32.2% | 20.2% | 1.48 ± 0.09 |
| DeepCDS S | 47.4% | 32.6% | 20.0% | 1.45 ± 0.08 |
| DeepCDS S+I | 46.6% | 30.9% | 22.5% | 1.51 ± 0.09 |
| **High-error (25,933 reads)** | | | | |
| FGS (Complete) | 33.5% | 24.3% | 42.3% | 1.38 ± 0.06 |
| FGS (0.5% error rate) | 28.1% | 17.8% | 54.1% | 1.58 ± 0.08 |
| FGS (1.0% error rate) | 26.4% | 16.5% | 57.1% | 1.60 ± 0.08 |
| MetaProdigal | 37.8% | 23.0% | 39.2% | 1.64 ± 0.08 |
| DeepCDS N | 45.0% | 31.0% | 24.0% | 1.45 ± 0.08 |
| DeepCDS S | 46.1% | 32.0% | 21.9% | 1.44 ± 0.08 |
| DeepCDS S+I | 45.5% | 30.3% | 24.2% | 1.50 ± 0.09 |
| **Stress-error (25,775 reads)** | | | | |
| FGS (Complete) | 29.4% | 22.1% | 48.5% | 1.33 ± 0.05 |
| FGS (0.5% error rate) | 24.6% | 16.4% | 59.1% | 1.51 ± 0.07 |
| FGS (1.0% error rate) | 23.1% | 15.3% | 61.6% | 1.50 ± 0.07 |
| MetaProdigal | 31.6% | 20.7% | 47.7% | 1.53 ± 0.07 |
| DeepCDS N | 37.8% | 27.0% | 35.2% | 1.40 ± 0.07 |
| DeepCDS S | 41.4% | 29.7% | 28.9% | 1.40 ± 0.07 |
| DeepCDS S+I | 40.7% | 27.9% | 31.4% | 1.46 ± 0.08 |

Table A29: Prediction of CDS start positions in the 300bp test sets for reads containing two annotated start positions (RefSeq vs. GenBank) with the same stop codon, in the 212 test genomes using the standard prokaryotic genetic code. Only reads in which both versions of the CDS were longer than 60 bp within the read are included. The "Annotated start position predicted" columns report the percentage of reads in which the model predicted the start position of the longer version, the shorter version, or neither. Ratio: number of reads with the longer start predicted divided by the number with the shorter start predicted. The ± marks the standard error from a genome-level cluster bootstrap (1,000 replicates).

| | FGS (Complete) | FGS (0.5% err) | FGS(1% err) | MetaProdigal | DeepCDS N | DeepCDS S | DeepCDS S+I |
|---|---|---|---|---|---|---|---|
| **Archaea** | | | | | | | |
| Error-free | -0.21 | - | - | 0.02 | 0.04 | - | - |
| Typical-error | -0.11 | -0.14 | -0.14 | 0.23 | 0.07 | -0.01 | -0.01 |
| High-error | 0.04 | -0.01 | -0.02 | 0.43* | 0.16 | 0.02 | 0.01 |
| Stress-error | 0.14 | 0.11 | 0.11 | 0.53* | 0.36* | 0.06 | 0.04 |
| **Bacteria** | | | | | | | |
| Error-free | 0.06 | - | - | -0.18* | -0.16* | - | - |
| Typical-error | 0.35* | 0.13 | 0.09 | 0.17* | 0.26* | 0.00 | 0.03 |
| High-error | 0.69* | 0.45* | 0.38* | 0.45* | 0.51* | 0.09 | 0.11 |
| Stress-error | 0.92* | 0.84* | 0.79* | 0.80* | 0.78* | 0.34* | 0.33* |

Table A30: Spearman correlation ($\rho$) between genomic GC-content and CDS-level F1 score across the 215 test genomes, reported for each error-rate condition at 300bp sequence length separately for archaeal and bacterial genomes. For the error-free test set, we only measured performance for the model variants trained on error-free sequences (i.e., FGS (Complete), MetaProdigal, and DeepCDS N). $^{*}p < 0.05$, marking a statistically significant correlation.

| Sequence length (bp) | Inference batch size | Model | GPU Reads/s | GPU Peak GPU memory (GB) | CPU Reads/s | CPU Peak RAM (GB) |
|---|---|---|---|---|---|---|
| 75 | 32 | DeepCDS S+I | 1,903 | 0.9 | 122.3 | 1.0 |
| | 128 | DeepCDS S+I | 2,961 | 1.2 | 142.6 | 1.5 |
| | 512 | DeepCDS S+I | 3,152 | 2.3 | 141.7 | 2.3 |
| | 2,048 | DeepCDS S+I | 3,158 | 6.5 | – | – |
| 100 | 32 | DeepCDS S+I | 1,629 | 0.9 | 97.0 | 1.2 |
| | 128 | DeepCDS S+I | 2,187 | 1.2 | 107.7 | 1.6 |
| | 512 | DeepCDS S+I | 2,376 | 2.7 | 108.9 | 2.4 |
| | 2,048 | DeepCDS S+I | 2,424 | 8.2 | – | – |
| 150 | 32 | DeepCDS S+I | 1,302 | 0.9 | 75.9 | 1.3 |
| | 128 | DeepCDS S+I | 1,673 | 1.5 | 80.2 | 1.9 |
| | 512 | DeepCDS S+I | 1,722 | 3.4 | 80.8 | 3.1 |
| | 2,048 | DeepCDS S+I | 1,752 | 11.3 | – | – |
| 300 | 32 | DeepCDS S+I | 712 | 1.2 | 33.6 | 1.4 |
| | 128 | DeepCDS S+I | 836 | 2.5 | 37.6 | 2.1 |
| | 512 | DeepCDS S+I | 848 | 7.4 | 41.9 | 5.5 |
| | 2,048 | DeepCDS S+I | 842 | 27.2 | – | – |
| 700 | 32 | DeepCDS N | 316 | 1.2 | 15.5 | 1.5 |
| | 128 | DeepCDS N | 333 | 2.4 | 15.2 | 2.1 |
| | 512 | DeepCDS N | 335 | 7.2 | 13.3 | 4.8 |
| | 2,048 | DeepCDS N | 326 | 26.5 | – | – |
| 1,000 | 32 | DeepCDS N | 189 | 1.2 | 9.4 | 1.6 |
| | 128 | DeepCDS N | 200 | 2.4 | 8.3 | 2.1 |
| | 512 | DeepCDS N | 204 | 7.2 | 6.8 | 4.8 |
| | 2,048 | DeepCDS N | 199 | 26.5 | – | – |

Table A31: Inference throughput and peak memory of DeepCDS on the test reads, predicting on both strands of each sequence, for each sequence length and a range of inference batch sizes. The GPU runs were executed on an NVIDIA H100 NVL. The CPU runs were executed on an AMD EPYC 9684X (32 threads). Reads/s mark the median of three timed passes over 100,000 (GPU) or 10,000 (CPU) sequences, after one warm-up batch, excluding model loading and file input/output. CPU runs were limited to batch sizes up to 512, as CPU throughput increased only marginally above a batch size of 128. We ran DeepCDS S+I on the read-length sequences, because it is the DeepCDS version with most parameters, thus giving an upper bound on the runtime of the three models. DeepCDS N was used for sequences longer than typical read lengths, with a sliding-window stride of 70 codons, as used in the benchmark. Please note that these measurements exclude model loading and file input/output, and were obtained on a different GPU (NVIDIA H100 NVL), they are not directly comparable to the end-to-end runtimes on the CAMI III sample, which were measured on an NVIDIA L40S GPU.

| CDS predicted by | Reads (%) | CDS with a UniRef90 hit (%) | | | | |
|---|---|---|---|---|---|---|
| | | N | S | S+I | MP | FGS |
| N + S + S+I + MP + FGS | 74.92 | 85.4 | **85.7** | 84.7 | 82.1 | 82.9 |
| S + S+I + MP + FGS | 2.10 | – | **62.9** | 59.9 | 50.4 | 51.6 |
| N + S + S+I + FGS | 1.44 | 69.9 | **70.2** | 68.2 | – | 58.3 |
| N + S+I + MP + FGS | 1.34 | **54.1** | – | 52.6 | 48.2 | 46.9 |
| N + S + S+I + MP | 0.70 | 69.5 | **69.7** | 68.8 | 62.8 | – |
| N + S + MP + FGS | 0.52 | 58.9 | **59.7** | – | 48.9 | 50.0 |
| S+I + MP + FGS | 2.52 | – | – | **43.4** | 38.2 | 37.6 |
| S + MP + FGS | 0.47 | – | **55.4** | – | 41.7 | 43.3 |
| N + MP + FGS | 0.36 | **42.5** | – | – | 37.6 | 37.4 |
| N + S + S+I | 0.29 | 60.5 | **60.6** | 59.2 | – | – |
| S + S+I + FGS | 0.24 | – | **51.9** | 49.5 | – | 38.4 |
| N + S+I + FGS | 0.23 | **43.4** | – | 42.0 | – | 34.6 |
| S + S+I + MP | 0.09 | – | **51.5** | 50.0 | 34.3 | – |
| N + S+I + MP | 0.06 | **42.3** | – | 41.5 | 36.7 | – |
| N + S + FGS | 0.05 | **50.3** | **50.3** | – | – | 40.9 |
| N + S + MP | 0.02 | 47.1 | **47.2** | – | 40.2 | – |
| MP + FGS | 4.27 | – | – | – | **23.7** | 23.6 |
| S+I + FGS | 0.81 | – | – | **30.2** | – | 25.5 |
| S+I + MP | 0.17 | – | – | **31.4** | 26.6 | – |
| S + S+I | 0.10 | – | **38.1** | 37.7 | – | – |
| N + FGS | 0.08 | **33.4** | – | – | – | 27.3 |
| N + S+I | 0.06 | **37.7** | – | 36.3 | – | – |
| S + FGS | 0.06 | – | **40.4** | – | – | 31.2 |
| S + MP | 0.03 | – | **34.1** | – | 26.6 | – |
| N + MP | 0.03 | **31.1** | – | – | 27.4 | – |
| N + S | 0.02 | 45.8 | **45.9** | – | – | – |
| FGS | 3.15 | – | – | – | – | 13.3 |
| MP | 0.91 | – | – | – | 11.9 | – |
| S+I | 0.33 | – | – | 20.0 | – | – |
| S | 0.04 | – | 25.2 | – | – | – |
| N | 0.03 | 32.3 | – | – | – | – |

Table A32: Share of predicted CDSs with a UniRef90 hit per combination of models on the real gut metagenome sample MH0161. Reads are grouped by which of DeepCDS N, S, and S+I (referred to as N, S, S+I), MetaProdigal (referred to as MP), and FGS (Complete) (referred to as FGS) predict at least one CDS longer than 60 bp on either strand. Of the FGS variants, only FGS (Complete) is included for readability, as it was the overall best-performing FGS variant. **Reads (%)**: share of all reads in the full dataset that fall into the group. **CDS with a UniRef90 hit (%)**: share of the CDSs predicted by each model on these reads with a UniRef90 hit (E-value $\leq 10^{-3}$). –: the model predicts no CDS on these reads. Bold marks the highest share in each row.

| | MetaProdigal | FGS (Complete) | DeepCDS N |
|---|---|---|---|
| FPs on sequences with ≥90% identity | 9,186 | 21,447 | 722 |
| overlapping rRNA or tRNA genes | 7,478 (81.4%) | 17,567 (81.9%) | 240 (33.2%) |
| overlapping rRNA genes | 5,913 (64.4%) | 14,218 (66.3%) | 100 (13.9%) |
| overlapping tRNA genes | 1,565 (17.0%) | 3,349 (15.6%) | 140 (19.4%) |

Table A33: FP CDS predictions on test sequences overlapping annotated rRNA and tRNA genes. FP CDSs predicted on the 300bp error-free test sequences (212 genomes using the standard prokaryotic genetic code) with ≥90% nucleotide identity to the best hit among the training and validation sequences ($E \leq 10$, using MMseqs2 [5]), counted on sequences that overlap an annotated rRNA or tRNA gene by more than 60 bp (gene on either strand; rRNA takes precedence over tRNA). Percentages are relative to each tool's total number of FPs on these sequences.

| | 60bp | 75bp | 100bp | 150bp | 300bp |
|---|---|---|---|---|---|
| **F1 Score** | | | | | |
| DeepCDS N | 0.815 | 0.862 | 0.906 | 0.941 | 0.955 |
| **Sensitivity** | | | | | |
| DeepCDS N | 0.835 | 0.868 | 0.906 | 0.933 | 0.950 |
| **Precision** | | | | | |
| DeepCDS N | 0.795 | 0.856 | 0.907 | 0.950 | 0.959 |

Table A34: Aggregated CDS-level performance based on the error-free test set of varying sequence lengths for the 212 test genomes that follow the standard prokaryotic translation table. The results show performance based on prediction of CDS fragments longer than 30 bp. Note that FGS and MetaProdigal only predict fragments of length 60 bp or longer, and are not included in the table. Standard errors are omitted from the cells for readability; across all metrics and read lengths, the genome-level cluster bootstrap standard error span is 0.001–0.004.

| | 60bp | 75bp | 100bp | 150bp | 300bp |
|---|---|---|---|---|---|
| **F1 Score** | | | | | |
| DeepCDS N | 0.805 | 0.851 | 0.892 | 0.919 | 0.912 |
| DeepCDS S | **0.813** | **0.858** | **0.900** | **0.930** | **0.933** |
| DeepCDS S+I | 0.793 | 0.839 | 0.884 | 0.923 | 0.928 |
| **Sensitivity** | | | | | |
| DeepCDS N | 0.822 | 0.855 | 0.889 | 0.910 | 0.911 |
| DeepCDS S | 0.828 | 0.862 | 0.896 | 0.920 | **0.930** |
| DeepCDS S+I | **0.864** | **0.884** | **0.905** | **0.921** | 0.924 |
| **Precision** | | | | | |
| DeepCDS N | 0.789 | 0.848 | 0.894 | 0.928 | 0.914 |
| DeepCDS S | **0.799** | **0.855** | **0.904** | **0.940** | **0.937** |
| DeepCDS S+I | 0.732 | 0.799 | 0.865 | 0.924 | 0.932 |

Table A35: Aggregated CDS-level performance based on the typical-error test set of varying sequence lengths for the 212 test genomes that follow the standard prokaryotic translation table. The results show performance based on prediction of CDS fragments longer than 30 bp. Note that FGS and MetaProdigal only predict fragments of length 60 bp or longer, and are not included in the table. The numbers marked in bold highlight the highest performance metric for the specific sequence length condition. Standard errors are omitted from the cells for readability; across all metrics and read lengths, the genome-level cluster bootstrap standard error spans 0.001–0.004 for all models.

| | 60bp | 75bp | 100bp | 150bp | 300bp |
|---|---|---|---|---|---|
| **F1 Score** | | | | | |
| DeepCDS N | 0.792 | 0.835 | 0.872 | 0.890 | 0.862 |
| DeepCDS S | **0.802** | **0.847** | **0.889** | **0.918** | **0.919** |
| DeepCDS S+I | 0.782 | 0.828 | 0.873 | 0.911 | 0.916 |
| **Sensitivity** | | | | | |
| DeepCDS N | 0.806 | 0.834 | 0.865 | 0.878 | 0.863 |
| DeepCDS S | 0.814 | 0.847 | 0.881 | 0.905 | **0.915** |
| DeepCDS S+I | **0.850** | **0.870** | **0.891** | **0.907** | 0.910 |
| **Precision** | | | | | |
| DeepCDS N | 0.779 | 0.837 | 0.879 | 0.903 | 0.861 |
| DeepCDS S | **0.791** | **0.848** | **0.897** | **0.932** | **0.924** |
| DeepCDS S+I | 0.724 | 0.790 | 0.856 | 0.915 | 0.921 |

Table A36: Aggregated CDS-level performance based on the high-error test set of varying sequence lengths for the 212 test genomes that follow the standard prokaryotic translation table. The results show performance based on prediction of CDS fragments longer than 30 bp. Note that FGS and MetaProdigal only predict fragments of length 60 bp or longer, and are not included in the table. The numbers marked in bold highlight the highest performance metric for the specific sequence length condition. Standard errors are omitted from the cells for readability; across all metrics and read lengths, the genome-level cluster bootstrap standard error spans 0.001–0.004 for all models.

| | 60bp | 75bp | 100bp | 150bp | 300bp |
|---|---|---|---|---|---|
| **F1 Score** | | | | | |
| DeepCDS N | 0.748 | 0.784 | 0.808 | 0.801 | 0.717 |
| DeepCDS S | **0.765** | **0.810** | **0.851** | **0.877** | 0.871 |
| DeepCDS S+I | 0.747 | 0.791 | 0.834 | 0.869 | **0.873** |
| **Sensitivity** | | | | | |
| DeepCDS N | 0.749 | 0.769 | 0.787 | 0.777 | 0.720 |
| DeepCDS S | 0.765 | 0.797 | 0.831 | 0.853 | 0.862 |
| DeepCDS S+I | **0.806** | **0.824** | **0.843** | **0.856** | **0.862** |
| **Precision** | | | | | |
| DeepCDS N | 0.747 | 0.800 | 0.830 | 0.826 | 0.714 |
| DeepCDS S | **0.766** | **0.823** | **0.872** | **0.903** | 0.880 |
| DeepCDS S+I | 0.695 | 0.760 | 0.824 | 0.883 | **0.884** |

Table A37: Aggregated CDS-level performance based on the stress-error test set of varying sequence lengths for the 212 test genomes that follow the standard prokaryotic translation table. The results show performance based on prediction of CDS fragments longer than 30 bp. Note that FGS and MetaProdigal only predict fragments of length 60 bp or longer, and are not included in the table. The numbers marked in bold highlight the highest performance metric for the specific sequence length condition. Standard errors are omitted from the cells for readability; across all metrics and read lengths, the genome-level cluster bootstrap standard error spans, per model: DeepCDS N 0.001–0.004; DeepCDS S 0.001–0.004; DeepCDS S+I 0.002–0.004.
